# Extending Mendelian randomisation to founder-haplotype data in Diversity Outbred mice

**DOI:** 10.64898/2026.11.10.763681

**Authors:** Xiaoyang Dai, Gibran Hemani, Larry D. Mesner, Gina M. Calabrese, Charles R. Farber, Gary Churchill, George Davey Smith

## Abstract

Observational associations can reflect confounding or reverse causation, which Mendelian randomisation (MR) avoids by using genetic variation to estimate causal effects. In Diversity Outbred (DO) mice such estimates can be tested experimentally, but applying human summary-data MR, with one scalar association per variant, ignores the differing effects of their eight founder haplotypes. We introduce Founder GLS-IVW, which uses each locus’s founder-effect pattern for causal estimation and diagnosis. At *Qsox1*, it recovered the direction of effect seen in knockout mice, which one scalar coding failed to detect. Across 4,596 trait pairs, scalar and founder analyses disagreed for 29.7%; in seven of eight, only the founder analysis found an effect, using information scalar coding discards. Because exposure and outcome may be measured in different or partly overlapping animals, the method supports two-sample MR across separate experiments. Our R package qtl2mr, built on the widely used R/qtl2, helps prioritise mechanistic experiments.

---

Because observational associations between traits can be distorted by confounding and reverse causation, Mendelian randomisation (MR), which uses the random inheritance of alleles to investigate exposure–outcome relationships, has become an important framework for investigating the aetiology of complex traits ^1,2^. In humans, its assumptions are difficult to test ^3^, and tissues such as bone and brain are rarely available at scale. Mouse populations offer a complementary setting: environments can be controlled, tissues sampled and predictions tested experimentally^4^, and their deep molecular and physiological phenotyping reveals more trait relationships than intervention can examine. MR in these resources could help to prioritise experiments and connect genetic evidence to biological mechanisms.

The genetic structure of Diversity Outbred (DO) mice presents a distinct challenge. Each animal is a mosaic of eight founder strains, and a local quantitative trait locus (QTL) can carry a pattern of effects across founder haplotypes ^5–10^. Simply extending human summary-data MR to these mice, with one scalar association for each genetic variant ^11,12^, ignores this founder-haplotype structure. A biallelic code groups founders into two classes, potentially combining opposing effects and obscuring an otherwise informative local association. A founder-aware scalar coding can preserve that signal when a suitable variant or grouping is known, but still commits the analysis to one representation.

At the *Qsox1* locus studied here, alternative scalar codings of the same strong expression QTL (eQTL) produce markedly different first-stage associations. The problem is therefore how to retain the genetic information on which inference depends.

Founder-haplotype effects provide both an instrument and a way to assess its consistency. Under a valid instrument model with a common linear exposure effect, the outcome pattern is proportional to the exposure pattern, *b*_*Y*_ = *θb*_*X*_. The scaling factor *θ* is the effect targeted by MR. Founders can differ in their effects on the exposure without violating this model; the common-effect assumption concerns the effect of the exposure on the outcome across haplotype backgrounds (IV4; Fig. 1c)^13,14^. A minimised lack-of-fit statistic, *Q*^min^, summarises departures from proportionality, and founder residuals locate the backgrounds contributing to them, although neither identifies the biological mechanism.

**Fig. 1.**
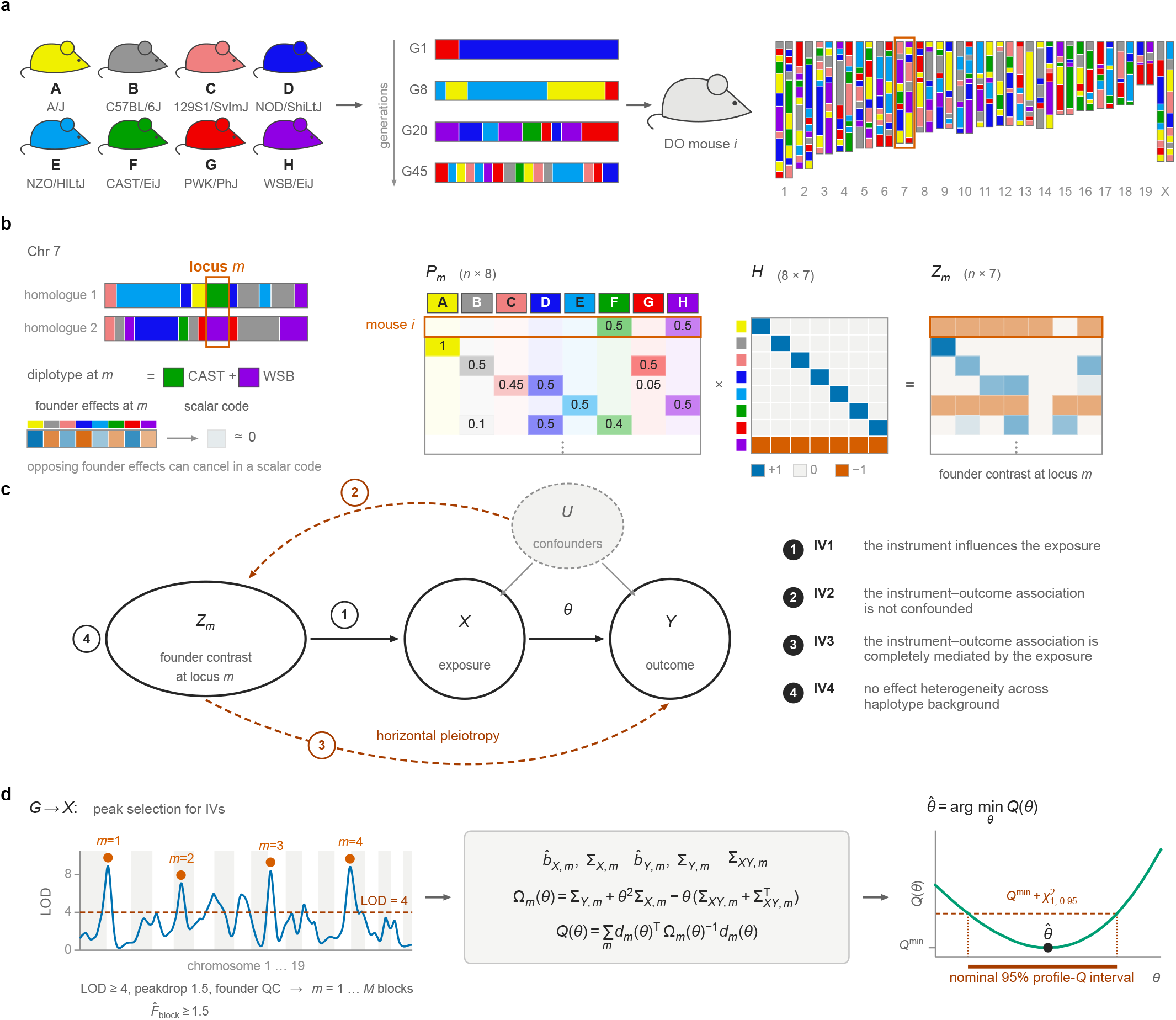
Founder-haplotype Mendelian randomisation in Diversity Outbred mice. **a**, The eight founder strains of the Diversity Outbred (DO) population (A–H) recombine over generations of outbreeding, so each DO mouse *i* carries a mosaic of founder haplotypes. Chromosome 7 is highlighted. **b**, At a locus *m* on chromosome 7, mouse *i* carries CAST/EiJ on one homologue and WSB/EiJ on the other. When founders have opposing effects, a scalar (biallelic) genotype code can cancel them. The founder-haplotype probabilities *P*_*m*_ (*n ×* 8; each row sums to 1) are multiplied by an 8 *×* 7 sum-to-zero contrast matrix *H* to give the seven-dimensional founder-contrast block *Z* _*m*_ = *P*_*m*_ *H* (*n ×* 7), which keeps the full pattern of founder effects. The *H* drawn is illustrative (each founder against WSB/EiJ). The analyses use the orthonormal Helmert basis; the estimate, its interval and *Q* ^min^ are basis-invariant (Methods). **c**, *Z* _*m*_ enters the standard instrumental-variable structure *Z* _*m*_ → *X* → *Y*, in which *θ* is the causal effect of the exposure *X* on the outcome *Y*, and *U* represents common causes of *X* and *Y*. Numbered labels mark the four instrument assumptions IV1–IV4 (key at right); IV4 requires the effect of *X* on *Y* not to vary across haplotype backgrounds, whereas founders may differ in their effects on *X*. Dashed arrows, possible violations of IV2 (instrument–outcome confounding) and IV3 (horizontal pleiotropy). **d**, Instruments are selected from the exposure genome scan (*G* → *X* ) as peaks with LOD ≥ 4 and a peakdrop of 1.5 that pass founder QC, keeping blocks with 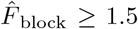; this gives blocks *m* = 1, …, *M* (Methods). The LOD trace is illustrative. For each block, the matched founder-contrast estimates 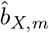 and 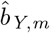, their covariances Σ_*X,m*_ and Σ_*Y,m*_, and the exposure–outcome cross-covariance Σ_*XY,m*_ define the overlap-aware second-order objective *Q* (*θ*). Its minimiser is the estimate 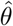. The nominal 95% profile-*Q* interval (approximate; Discussion) contains every *θ* with *Q* (*θ*) ≤ *Q* ^min^ + *χ*^2^_1,0.95_. Founder colours are the standard Collaborative Cross colours; notation, Methods.

Here we introduce Founder GLS-IVW (founder-block generalised least-squares inverse-variance-weighted MR), which combines kinship-adjusted founder-effect reconstruction with overlap-aware second-order estimation ^15–17^. QTL-anchored causal inference has a long history in model-organism populations ^18,19^, including single-SNP summary-data MR in outbred heterogeneous stock rats ^20^ and founder-haplotype mediation analyses in DO mice ^21,22^; the closest DO method, bmediatR, performs Bayesian model selection for mediation rather than MR estimation ^23^. Each trait is fitted on its own sample, so the exposure and outcome can be measured in the same, different or partly overlapping mice, and the covariance between their founder-effect estimates is built from the mice measured for both traits. We demonstrate how representation changes inference at *Qsox1*, use simulations to characterise estimation, interval coverage and diagnostics, and extend the analysis to multiple loci. Across a landscape of 4,596 trait pairs, we then compare Founder GLS-IVW with the direct extension of human MR, the SNP-level inverse-variance-weighted estimate, to measure what that extension may miss and to separate the contribution of the founder representation from that of overlap-aware estimation. The qtl2mr package implements these analyses for deeply phenotyped founder-based resources.

## Results

### Founder-haplotype Mendelian randomisation

Figure 1 outlines the approach. Each DO mouse carries a mosaic of the eight founder genomes (Fig. 1a), and at a given locus qtl2 records the probability that each founder contributed the local haplotype ^8^. Because a single allele code can make opposing founder effects cancel, we transform the eight probabilities into seven founder contrasts that keep the full pattern (Fig. 1b). This founder block is the instrument, under the usual instrumental-variable assumptions (Fig. 1c). For each locus that passes a logarithm-of-odds (LOD) threshold and peak selection, we estimate the founder effects on the exposure and on the outcome and find the single scaling factor, the MR estimate, that best matches the two patterns, allowing for the uncertainty in both and for animals measured for both traits (second-order, SO, weighting; Fig. 1d). What it cannot explain is summarised by the lack-of-fit statistic *Q*^min^ and by founder-level residuals (Methods).

### Scalar coding can conceal a strong local signal

We used the published data of the Farber laboratory DO bone cohort, a systems-genetics study of bone morphology and strength ^24^: 619 phenotyped mice, 192 with cortical-bone RNA-seq. Standardised *Qsox1* expression has a founder eQTL peak (LOD = 24.42) at the GigaMUGA^25^ marker JAX00272363, in a region that also contains bone-trait QTL (Extended Data Fig. 1); this overlap motivates the analysis but does not by itself establish mediation by *Qsox1*.

Alternative scalar codings captured very different amounts of this signal (Fig. 2a,b). The GigaMUGA 0/1/2 marker gave a first-stage *F* = 3.87 (*P* = 0.049), whereas the founder-aware scan1snps dosage rs232318063 gave *F* = 101.70 (*P* = 6.46 *×* 10^−24^). The seven-contrast founder block gave a per-contrast first-stage statistic of *F*_block_ = 15.36 (Methods). The founder effects explain the difference: CAST/EiJ increases *Qsox1* expression, whereas PWK/PhJ and WSB/EiJ decrease it (mean-centred effects of 0.94, −0.58 and −1.02 expression standard deviations, SD). The GigaMUGA code groups all three together, so their opposing effects cancel, whereas the scan1snps code separates CAST/EiJ from the other two.

**Fig. 2.**
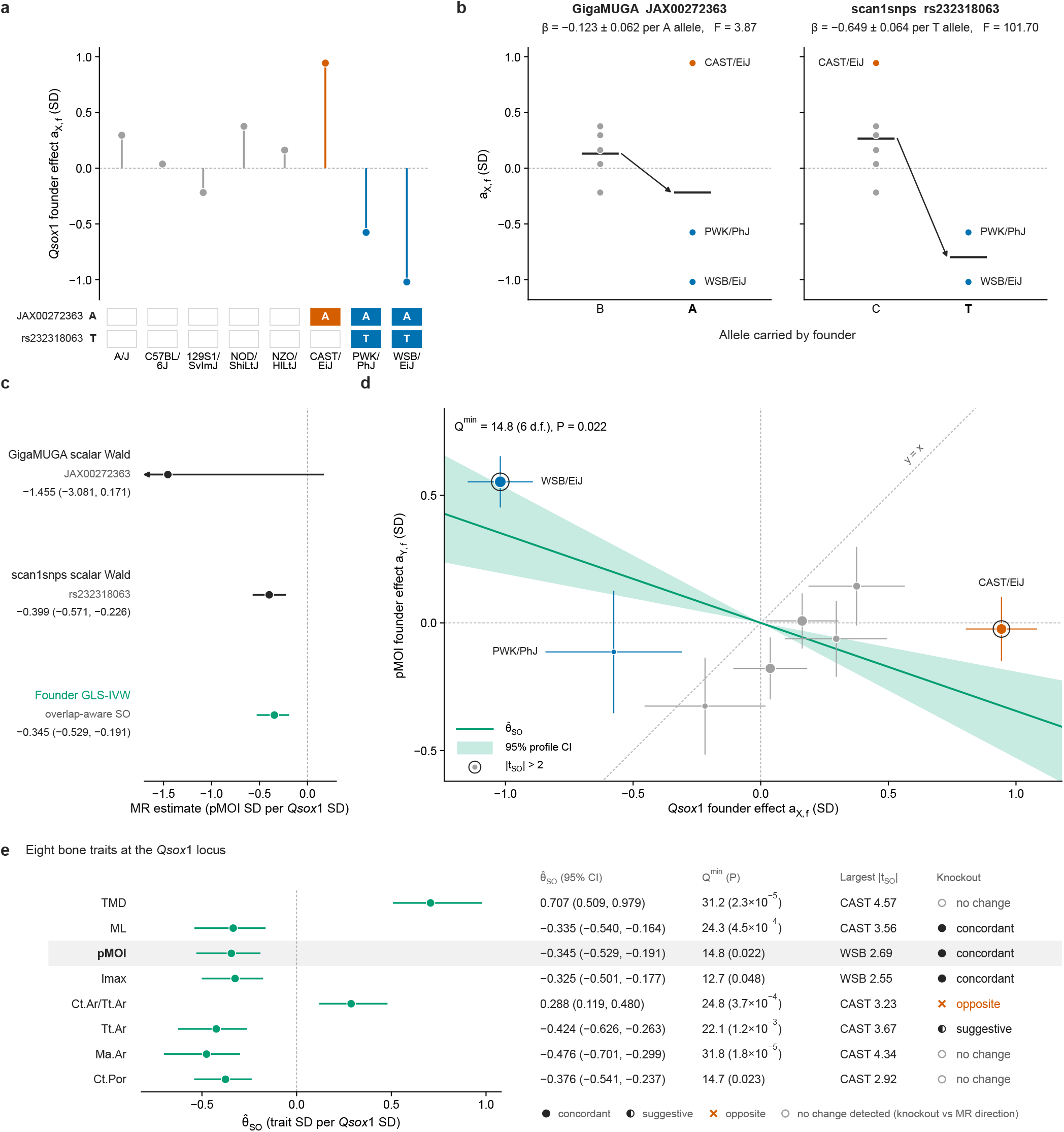
Coding-dependent information loss, founder-block MR and within-locus diagnostics at the *Qsox1* locus. **a**, Mean-centred *Qsox1* founder effects *a*_X,f_ (expression SD) at the lead eQTL marker JAX00272363. Below, founders carrying the allele counted by each scalar dosage: A at GigaMUGA JAX00272363 and T at scan1snps rs232318063. **b**, Founder effects from **a**, grouped by allele. Bars, unweighted group means; arrow, their difference (illustrative, not the fitted coefficient). *β±* s.e. (per copy of the counted allele) and *F* come from the fitted scalar first stage (*n* = 192 mice). **c**, Estimates of *Qsox1* → pMOI (pMOI, polar moment of inertia measured by *µ*CT), in pMOI SD per *Qsox1* SD, with 95% CIs. Scalar rows are Wald ratios with estimate *±*1.96 s.e. intervals. The founder row is the overlap-aware second-order (SO) Founder GLS-IVW estimate with its nominal profile-*Q* interval. Arrow, interval cut off at the axis limit. **d**, pMOI founder effects against *Qsox1* founder effects. Crosshairs show *±*1 marginal s.e.; point area is proportional to 1/Var(*a*_X,f_). Line, 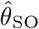 through the origin; tint, its nominal profile-*Q* 95% CI; dashed lines, *x* = 0, *y* = 0 and *y* = *x*. Rings mark founders whose one-founder-versus-rest residual has |*t* _SO_| *>* 2. *Q* ^min^, over-identification statistic with an approximate *χ*^2^_6_ reference. Exposure *n* = 192; outcome *n* = 611; 187 mice overlap. **e**, Eight bone traits at the *Qsox1* locus: 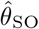 with its 95% profile-*Q* CI (trait SD per *Qsox1* SD), *Q* ^min^ with its approximate *χ*^2^_6_ *P* value, the founder with the largest |*t* _SO_|, and the change after *Qsox1* loss of function in published experiments ^24^ relative to the MR direction (an increase after loss of function is concordant with a negative estimate; details in Extended Data Fig. 2b). *P* values are nominal, and the eight traits are correlated. Outcome *n* = 611 for all traits except ML (591); 187 of these mice overlap with the exposure sample (169 for ML). TMD, cortical tissue mineral density; ML, medial–lateral femoral width; Imax, maximum moment of inertia; Ct.Ar/Tt.Ar, cortical area fraction; Tt.Ar, total area; Ma.Ar, medullary area; Ct.Por, cortical porosity.

The choice also changed the MR result (Fig. 2c). For *Qsox1* expression and polar moment of inertia (pMOI; *n* = 611, including 187 mice with expression data), the GigaMUGA Wald ratio was −1.455 pMOI SD per *Qsox1* SD (95% CI −3.081 to 0.171). In contrast, the scan1snps ratio, −0.399 (−0.571 to −0.226), and Founder GLS-IVW estimate, −0.345 (−0.529 to −0.191), agreed in direction and magnitude and excluded zero. A scalar coding can therefore recover the association only if its grouping preserves the relevant founder pattern; Founder GLS-IVW retains that pattern without choosing a grouping.

The knockout experiment of the original study is the most direct test of these estimates. Male *Qsox1* -mutant mice lacking QSOX1 activity had higher pMOI than wild-type mice ^24^, the direction implied by a negative estimate. Founder GLS-IVW recovered this experimentally supported effect, as did the scan1snps coding, whereas the GigaMUGA coding did not: how founders are grouped into a scalar code decides whether the effect is seen at all. The founder block requires no such choice.

### Founder effects provide within-locus diagnostics

Founder effects also show when one causal effect cannot explain a locus. If the exposure acts on the outcome through a single common effect, each founder’s effect on the outcome is the same multiple of its effect on the exposure, so the founder effects lie on a line through the origin (Fig. 2d). The lack-of-fit statistic *Q*^min^ tests this proportionality, and one-founder-versus-rest residual scores show which founder haplotypes depart from it. A scalar analysis, with one association per locus, can report only the estimate.

At *Qsox1* we examined eight femoral traits: two measures of resistance to twisting and bending (pMOI and the maximum moment of inertia), three measures of bone size (medial–lateral width, total cross-sectional area and the area of the marrow cavity), the fraction of the cross-section occupied by cortical bone, and cortical mineral density and porosity. All eight showed nominal lack of fit (*Q*^min^ *P* = 1.8 *×* 10^−5^ to 0.048; Fig. 2e and Extended Data Fig. 2a), and in every trait the largest residual came from one of two founders, CAST/EiJ (six traits) or WSB/EiJ (pMOI and the maximum moment of inertia); for pMOI, CAST/EiJ raised *Qsox1* expression most yet had almost no effect on pMOI. This points to more than one effect at the locus, in agreement with the original study, in which an *Ier5* eQTL colocalised with all eight trait QTL and conditional mapping indicated at least two WSB/EiJ-driven loci^24^. The knockout results fit the same picture. Width and the maximum moment of inertia, and suggestively total area, changed in the directions implied by their estimates, whereas the cortical fraction changed in the opposite direction and no change was detected in marrow area, mineral density or porosity (Fig. 2e; Extended Data Fig. 2b). The two traits that disagreed most clearly, cortical fraction and marrow area, had strong evidence of lack of fit (*Q*^min^ *P* = 3.7 *×* 10^−4^ and 1.8 *×* 10^−5^): in DO mice the WSB/EiJ haplotype enlarged the marrow cavity, which loss of QSOX1 activity did not.

Founder MR therefore reports more than an estimate. It shows whether the founder pattern supports a single causal effect and, if not, which haplotypes depart from it. At *Qsox1*, these diagnostics, computed from the DO data alone, agreed with what colocalisation and knockout experiments showed: the locus carries more than one effect, and a single *Qsox1* -mediated effect does not explain every trait. Such signals can be flagged before experiments are designed.

### Simulations show calibration at moderate strength and approximate intervals at weak strength

We evaluated Founder GLS-IVW in simulations in which the causal effect is known (Fig. 3a). Single-locus calibration experiments kept the genotypes and kinship of real DO mice and simulated only founder effects and noise, the noise drawn from the covariance model used in fitting; multilocus experiments used synthetic designs or summary-level coefficient blocks. These experiments therefore test the estimator when its covariance model is correct. Instrument strength was set by *F*_target_, the population founder-block signal per contrast; the fitted statistic *F*_block_ also includes sampling variation, and at *F*_target_ = 1.5 it averages about 2.5 (median 2.95 for the real blocks analysed below); results for every setting are in Supplementary Note 4.

**Fig. 3.**
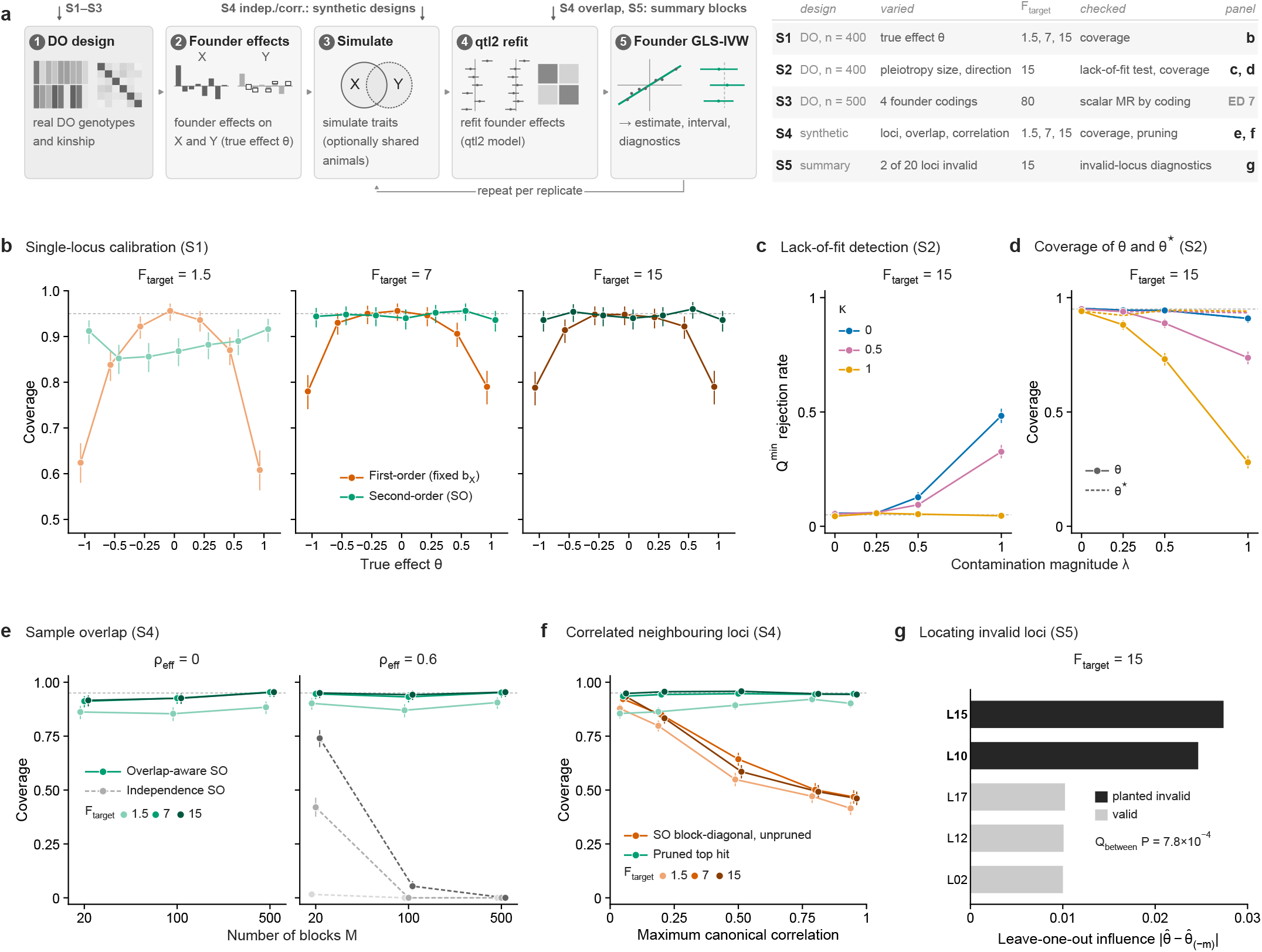
Simulation study of Founder GLS-IVW. No replicate or block was selected on 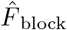. **a**, Design. S1–S3 use real Diversity Outbred (DO) genotypes and kinship; the S4 locus arms use synthetic designs; the S4 overlap arm and S5 use summary-level coefficient blocks (Methods). The table lists the *F* _target_ levels shown; S3 and the S4 independent arm are in Extended Data Figs 7 and 4. **b**, S1 (single locus, *n* = 400, no pleiotropy, 500 replicates per *θ*): coverage of first-order Wald intervals (fixed *b*_X_) and second-order (SO) sets against the true effect *θ*. **c**,**d**, S2 (single locus, *n* = 400, *θ* = 0.25, full sample overlap with residual correlation 0.2, *F* _target_ = 15, 1,000 replicates): pleiotropic contamination of magnitude *λ* and alignment with the exposure effects *κ* (0, orthogonal; 1, fully aligned). **c**, *Q* ^min^ rejection rate (*α* = 0.05). **d**, Coverage of *θ* (solid) and of the projection slope *θ*^⋆^ (dashed, no bars). **e**, S4 sample overlap (seven-dimensional blocks, *θ* = 0.25, 500 replicates): coverage against the number of blocks *M* at effective coefficient-error correlation *ρ*_eff_ = 0 and 0.6 for the overlap-aware SO objective (green, solid) and, at *ρ*_eff_ = 0.6, the independence objective (grey, dashed). **f**, S4 correlated neighbouring loci (*M* = 10, *n* = 3,000, *θ* = 0.25, 1,000 replicates): coverage against the maximum canonical correlation between blocks, for the unpruned SO block-diagonal analysis (orange) and one pre-specified block per cluster (‘pruned top hit’, green). **g**, S5 (*M* = 20 loci at *F* _target_ = 15, *θ* = 0.25, 200 replicates, two planted invalid loci): leave-one-out influences of the five most influential loci in a representative replicate chosen by a fixed rule (Methods); black, planted invalid loci; *P*, between-locus heterogeneity test (*Q* _between_). Shading from light to dark marks *F* _target_ = 1.5, 7 and 15 (b, e and f). Coverage, the proportion of replicates whose exact 95% profile-*Q* set (possibly unbounded; Supplementary Table 1b) contains *θ* (Methods). Bars, Clopper–Pearson 95% CIs. Dashed grey lines, nominal 0.95 coverage or 0.05 rejection. Further results: Supplementary Note 4.

When the founder effects on the outcome were proportional to those on the exposure, the SO estimate was centred on the true effect, and its profile-*Q* intervals covered it at close to the nominal rate (0.94–0.96 at *F*_target_ of 7 and 15; Fig. 3b and Extended Data Fig. 3). Intervals that treat the estimated exposure effects as fixed were too narrow for larger effects, which is why the method propagates their uncertainty.

Combining loci raises two dependencies. When the exposure and outcome are measured in the same animals, their estimated founder effects share errors; ignoring this repeats the same error at every locus, so that at an error correlation of 0.6 and *F*_target_ = 7, coverage fell from 0.42 with 20 loci to zero with 100. With the overlap covariance known, the overlap-aware estimate kept coverage at 0.93–0.95 up to 500 loci at *F*_target_ of 7 and 15 (Fig. 3e and Extended Data Fig. 4). Neighbouring loci can also be correlated: treating correlated blocks as independent reduced coverage to below 0.5 at high correlation, whereas analysing one pre-specified representative block per correlated cluster preserved it at *F*_target_ of 7 and 15 (Fig. 3f). The instrument sets of the real analyses can retain linked loci, so this condition is not guaranteed there (Methods).

At weak strength (*F*_target_ = 1.5), multi-locus SO estimates remained approximately centred when blocks were not selected on their fitted strength, but coverage fell below nominal (0.85–0.91), so intervals from real analyses that combine many weak blocks are approximate and may be too narrow; selection in the same sample is considered in the Discussion. Two-step IVW, weighted median and weighted mode estimates, by contrast, were biased towards zero even when every locus was valid (Extended Data Fig. 5).

The diagnostics were tested by adding a pleiotropic component to the outcome founder effects at one locus (Fig. 3c,d). Its size relative to the causal signal is *λ* (*λ* = 0 for a valid instrument, *λ* = 1 for pleiotropy as large as the causal effect) and its shape is *κ*: at *κ* = 1 it copies the exposure founder pattern, so the outcome pattern stays proportional with a different slope; at *κ* = 0 it is unrelated to the exposure pattern. As *λ* grew, *Q*^min^ rejected more often the further *κ* was from 1. Conversely, the closer *κ* was to 1, the more the estimate moved towards the slope of the contaminated pattern (*θ*^⋆^), so that coverage of *θ* fell while *θ*^⋆^ remained covered (Extended Data Fig. 6). Within a locus, *Q*^min^ therefore detects pleiotropy that changes the shape of the founder pattern, whereas pleiotropy that copies the exposure pattern is indistinguishable from a different causal effect and is absorbed into the estimate. Comparing loci addresses this second case. When two of 20 loci carried such a shifted but proportional effect, the global SO interval lost coverage (weighted median and mode estimates were less affected), but between-locus heterogeneity and leave-one-out analysis identified the invalid loci (Fig. 3g and Extended Data Fig. 5). When the outcome pattern was not proportional, different binary codings of the same locus also targeted different scalar estimands (Extended Data Fig. 7), a choice the founder block does not require.

### A positive control: femoral width and resistance to twisting

As a positive control for the multi-locus analysis, we estimated the effect of femoral width on the femur’s resistance to twisting. Medial–lateral femoral width (ML) is the width of the femoral cross-section, and the polar moment of inertia (pMOI) measures how strongly that cross-section resists torsion. Because pMOI grows with the distance of bone from the centre of the section, a wider femur is expected to have a larger pMOI. Each trait was fitted on its own sample (ML, 779 mice; pMOI, 799; 779 shared), and 55 of 57 candidate loci from the ML genome scan passed the inclusion threshold (Methods).

Founder GLS-IVW estimated that a 1-SD wider femur increases pMOI by 0.728 SD (95% profile-*Q* CI 0.680–0.775; Fig. 4a). The package’s two-step IVW (0.745), weighted median (0.747) and weighted mode (0.705) agreed, as did the scalar SNP-IVW on the same loci (0.770), whereas the covariate-adjusted observational association, in the mice measured for both traits, was larger (0.830). The estimate was stable: omitting one locus at a time moved it by at most 0.0098 (locus UNC8027644), and the Steiger-restricted, *F*_block_ ≥ 2 and alternative-list analyses gave 0.703 to 0.739 (Supplementary Table 9(b)); refits on instrument subsets ranged from 0.696 to 0.805 (Extended Data Fig. 8c). The founder-block diagnostic indicated some lack of fit (*Q*^min^ = 472.57 on 384 degrees of freedom, *P* = 0.0013, *χ*^2^ reference); the per-locus heterogeneity statistic, which we report as an uncalibrated diagnostic, was *Q*_between_ = 64.0 on 54 degrees of freedom (nominal *P* = 0.17). We then fitted the reverse direction. With pMOI as the exposure, 45 loci gave 0.849 ML SD per pMOI SD (0.790–0.910), but this estimate was not robust: restricted to loci acting more strongly on pMOI than on ML (the Steiger condition), it fell to 0.657 (0.586–0.726), outside that interval, and the 45-locus fit showed strong lack of fit (*Q*^min^ *P* = 2.8 *×* 10^−6^; Supplementary Table 9).

**Fig. 4.**
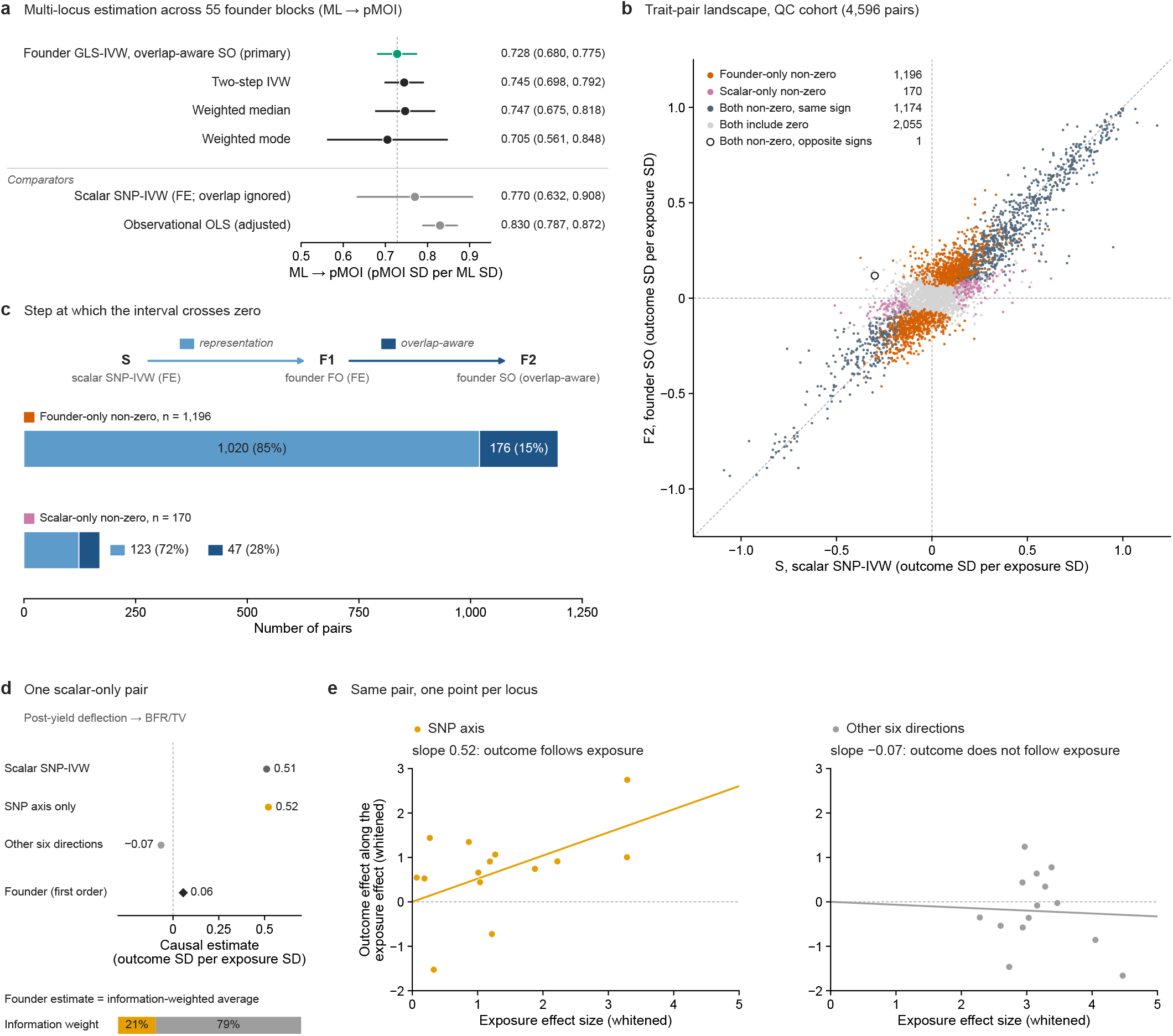
Multi-locus estimation and the trait-pair landscape. **a**, Effect of medial–lateral femoral width (ML) on polar moment of inertia (pMOI, *µ*CT) from 55 founder blocks with 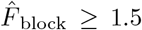; each trait uses its own sample (ML, *n* = 779; pMOI, *n* = 799; 779 shared). Green, overlap-aware second-order (SO) Founder GLS-IVW (primary, dashed line; nominal 95% profile-*Q* interval, approximate). Two-step IVW, weighted median and weighted mode combine per-block estimates (95% Wald intervals; median and mode, parametric-bootstrap SE). Below the line: scalar SNP-IVW (S) at the same 55 loci, and observational least squares (shared mice; adjusted for sex, age and generation). Values, Supplementary Table 9. **b**, Founder SO estimate (F2) against S for the 4,596 ordered pairs of the QC cohort (of 5,402 planned; Extended Data Fig. 9). All tiers of a pair use the same loci; each trait, its own sample. Colour, endpoint interval state: founder-only non-zero, S interval includes zero, F2 excludes it; scalar-only non-zero, the reverse. Intervals: S, fixed-effect (FE) Wald; F2, nominal 95% profile-*Q*. Dashed lines, *y* = *x* and zero lines. Counts without the 290 planned pairs involving Imax or Imin, Supplementary Table 3. **c**, F1, the founder first-order estimate, has an FE Wald interval. Step 1 (representation) replaces the scalar code by the founder block; step 2 (overlap-aware) adds the exposure–outcome cross-covariance and second-order terms. Bars, number of pairs in each discordant cohort (founder-only non-zero, 1,196; scalar-only non-zero, 170) by the step at which the interval crosses zero: step 1 (light blue) or step 2 (dark blue); labels, count (percentage of the cohort). Each pair crosses at exactly one step. **d**, One scalar-only pair (BFR/TV, bone formation rate per total volume): S, SNP-axis and other-six first-order estimates, and F1. On its 14 complete loci, F1 is the information-weighted average of the SNP-axis and other-six estimates (weights 21% and 79%); F1 shown uses all 15 loci. **e**, The pair of **d** per locus, on the SNP axis (left) and other six directions (right): whitened exposure effect size (*x* ) and outcome effect along it (*y* ); lines through the origin, slopes of **d**.

This positive control shows how founder MR separates causation from association: the observational association exceeded the causal estimate, and the reverse direction was not robust, whereas the forward estimate changed little under the Steiger condition and the alternative instrument lists. From many loci of modest strength, Founder GLS-IVW recovered a causal effect of femoral width on pMOI in the expected direction.

### Scalar–founder differences extend across the phenotypic landscape

We next compared scalar and founder MR across 74 phenotypes of the same cohort, spanning bone, adiposity, body size, glucose, muscle and marrow-adipose traits. Of the 74 *×* 73 = 5,402 ordered exposure–outcome pairs, 4,596 passed the estimability and quality-control (QC) checks and form the cohort analysed here (Methods; Extended Data Fig. 9). For each pair, three analyses used the same loci, those meeting the Steiger condition, with each trait fitted on its own sample. S is the scalar SNP-IVW of human summary-data MR: each locus is reduced to its marker dosage, sample overlap is ignored and the standard error is fixed-effect, so S gives the point estimate of a conventional IVW analysis. F1 replaces the marker with the full founder block but keeps first-order estimation with a fixed-effect standard error. F2 is Founder GLS-IVW, the overlap-aware SO estimate with its profile-*Q* interval. The representation step, S→F1, therefore changes only how each locus is coded; the overlap-aware step, F1→F2, adds the second-order terms and the exposure–outcome covariance from shared mice. All three intervals are fixed-effect intervals: none is inflated for between-locus heterogeneity. The S and F2 intervals differed in whether they excluded zero for 1,366 pairs (29.7%; Fig. 4b): for 1,196 only the founder interval excluded zero, and for 170 only the scalar interval did, about seven to one.

Most of these differences arose when the founder block replaced the marker (Fig. 4c): 1,020 of the 1,196 founder-only pairs (85%) and 123 of the 170 scalar-only pairs (72%) changed their conclusion at the representation step, and the rest at the overlap-aware step. The same balance held when the inclusion threshold was raised to *F*_block_ ≥ 2 (1,181 and 168 pairs) or when instruments were selected with a 2-LOD peak drop (1,140 and 159; Supplementary Table 3).

The scalar analysis sees only one direction of each founder block: the contrast between the founders that carry the two alleles of the marker SNP. We therefore split each founder block into this SNP axis and the six remaining directions (Methods). The SNP axis carried only about a fifth of the information for the causal estimate (medians 18–20% across groups of pairs, after correction for estimation noise in the exposure effects), where the information of a direction is the precision of the first-order estimate it gives. In the 123 scalar-only pairs that changed at the representation step, exposure and outcome effects agreed along the SNP axis but not along the other six directions (median concordance index 0.88 and −0.07 in the 85 and 68 pairs where it could be estimated, where 1 means proportional patterns and 0 unrelated ones), whereas in a random sample of 100 pairs significant under both analyses they agreed along both (0.91 and 0.94), a contrast that partly follows from how the groups were defined (Supplementary Note 6). Fig. 4d,e show one such pair, post-yield deflection → bone formation rate per total volume (BFR/TV). Along the SNP axis alone the estimate was 0.52, essentially the scalar estimate; the other six directions, which carry about four-fifths of the information, gave −0.07; and the first-order founder estimate, the information-weighted average of the two, was 0.06. A scalar-only signal therefore arises when exposure and outcome happen to agree along the one direction a SNP can see, while the rest of the founder information does not support it (Supplementary Note 6). For 11 of the 47 pairs that changed at the overlap-aware step, the SNP-axis signal was significant only when sample overlap was ignored.

The landscape thus shows what the founder representation adds to a conventional scalar analysis. Using all founder directions rather than the one a SNP sees, Founder GLS-IVW found many more pairs whose interval excluded zero only under the founder analysis than the reverse, and it identified scalar-only signals that rest on a single direction carrying little of the exposure information, which are potential false positives. Applied across 4,596 trait pairs of one cohort, the same workflow ran as a routine scan, so that causal hypotheses can be screened across a deeply pheno-typed resource, providing supporting evidence and prioritising experiments on biological mechanisms.

## Discussion

Founder GLS-IVW extends MR to the genetic representation used in DO mice: rather than reducing a local QTL to one scalar association, as a direct extension of human summary-data MR would, it retains the founder-effect pattern and uses it for both estimation and diagnosis. At *Qsox1*, opposing founder effects made a strong eQTL nearly invisible to the array marker, whereas the founder block recovered an estimate whose direction agrees with published knockout experiments and whose founder residuals point to the haplotypes behind the traits that the knockout did not reproduce. Across 4,596 trait pairs, the direct scalar extension and Founder GLS-IVW reached different interval conclusions for about three pairs in ten, and for seven in eight of these the founder interval excluded zero where the scalar interval did not. Most of these changes appeared at the representation step, when the founder block replaced the marker; fewer came from the overlap-aware step.

The estimate describes how the outcome pattern scales with the exposure pattern, and the diagnostics identify departures from proportionality. These can reflect horizontal pleiotropy, an exposure effect that varies across haplotype backgrounds, or a nonlinear or buffered exposure–outcome relationship, which the diagnostics do not distinguish. A component that copies the exposure pattern escapes *Q*^min^ and must be sought by comparing loci, and large non-proportional components can shift the estimate (Supplementary Note 4), so the estimate and its residual structure must be interpreted together. Beyond the between-locus diagnostics available to scalar MR, founder effects reveal departures within a locus and identify haplotype backgrounds for closer investigation.

The main limitation is that the intervals of multi-locus analyses are approximate and may be too narrow. In typical human summary-data MR, the associations of each instrument are estimated in very large samples. A DO cohort comprises hundreds of mice, yet a multi-locus analysis estimates seven founder contrasts at each of many loci, so individual blocks are often weak (median *F*_block_ = 2.95 in the ML–pMOI analysis) and their estimated exposure effects carry substantial error. We therefore report the profile-*Q* interval, which propagates this error and need not be symmetric about the estimate, and read it as approximate. In our simulations, which fitted the covariance model that generated the data, multi-locus estimates at weak strength remained approximately centred when blocks were not selected on their fitted strength, but their intervals covered the true effect less often than nominal.

Real analyses can depart further from these conditions. As in any MR analysis, selecting instruments in the same sample used for estimation can introduce a winner’s-curse bias^26^; our simulations indicate that this also affects weak founder blocks, with coverage that worsens as loci are added (Supplementary Note 4), and the loci of our real analyses were selected in the same mice. Linked peaks on the same chromosome are also treated as independent, which in simulation reduced coverage when blocks were strongly correlated. Selecting loci in independent data, or cross-fitting across sample splits ^27^, could separate selection from estimation; adapting these approaches to founder blocks is a natural next step. The scalar interval, which ignores the uncertainty in the exposure associations and any sample overlap, is also approximate. The landscape therefore compares interval conclusions rather than error rates, and the excess of founder-only over scalar-only pairs may partly reflect narrower founder intervals rather than additional signal alone.

Other elements of the method are working approximations. The star-space cross-covariance is a qtl2-aligned working model rather than a fitted bivariate mixed model. Its accuracy was checked at *F*_target_ = 100 and nominal residual correlations no greater than 0.2 (Supplementary Note 4, Section 4.6), narrower conditions than the weak, strongly overlapping blocks in the ML–pMOI analysis. Founder probabilities and heritabilities are also treated as fixed, without propagating their uncertainty.

Several extensions follow. Because a founder block offers seven directions at each locus, a multivariable extension ^28,29^ could separate the effects of exposures that share a locus, provided their founder patterns differ, such as the expression of *Qsox1* and of the colocalising *Ier5*. The approach could also be adapted for the Collaborative Cross ^30^, heterogeneous stock mice and rats ^31,32^ and the *Drosophila* Synthetic Population Resource ^33^, because the contrast construction is not restricted to eight founders and qtl2 supports multiple crossing designs ^8^. In humans, haplotype contrasts might resolve opposing effects obscured in single-SNP associations, extending haplotype and regional-variant methods ^34–36^; this would require haplotype-level effects and covariance rather than standard summary statistics, and local ancestry in admixed cohorts raises its own issues ^37,38^. Finally, mouse resources allow these predictions to be tested experimentally. At *Qsox1*, CAST/EiJ raises expression with almost no effect on pMOI, whereas WSB/EiJ lowers expression and raises pMOI; perturbations on these backgrounds could ask whether the locus acts through *Qsox1* expression alone. By retaining genetic structure that scalar coding can obscure, Founder GLS-IVW can help to prioritise such experiments and to use deeply phenotyped mouse populations to investigate relationships that are difficult to study in humans.

## Methods

### Founder-contrast representation and qtl2-compatible coefficient fitting

At a locus *m* (index suppressed where unambiguous), let *P* ∈ ℝ^*n×*8^ be the R/qtl2 founder-probability or founder-dosage matrix for the *n* mice analysed ^7,8^; regressing on *P* follows the Haley– Knott approach ^39^. Rows sum to a constant (*P* **1**_8_ = *c***1**_*n*_, with *c* = 1 for probabilities and *c* = 2 for dosages), so in a model with an intercept only founder contrasts are identifiable. We therefore use the founder-contrast block *Z* = *PH*, where *H* ∈ ℝ^8*×*7^ is a fixed orthonormal Helmert basis (*H*^T^**1**_8_ = 0, *H*^T^*H* = *I*_7_), and a contrast-coefficient vector *b*_*T*_ ∈ ℝ^7^ for trait *T* maps to sum-to-zero founder effects *a*_*T*_ = *Hb*_*T*_ . *H* is a coordinate system, not a biological assumption: it is fixed and shared by all traits, and the Founder GLS-IVW estimate, its interval, *Q*^min^ and the diagnostics are invariant to any change of contrast basis applied consistently to coefficients and covariances (Supplementary Note 2).

For each trait, coefficients are estimated in the R/qtl2 founder-contrast linear mixed model ^40,41^

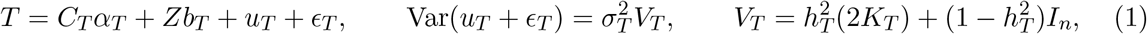

where *C*_*T*_ contains the intercept and covariates, *K*_*T*_ is the R/qtl2 kinship matrix (the factor 2 follows the R/qtl2 convention) and the residual heritability 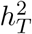 is estimated by restricted maximum likelihood (REML) under the null model without the founder term and then held fixed. With 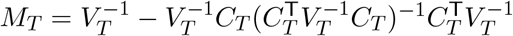,

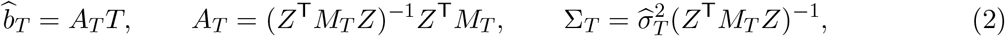

where *A*_*T*_ is the coefficient influence operator and Σ_*T*_ the full 7 *×* 7 coefficient covariance. Because R/qtl2 (`scan1coef`) returns only 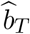 and marginal standard errors, a qtl2-compatible full-covariance fitter reproduces the same model and also returns Σ_*T*_, *A*_*T*_, the residuals and 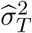; its estimates and standard errors were checked against `scan1coef` at the same fixed 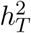 in the *Qsox1* and ML–pMOI analyses (Supplementary Table 4).

### Exposure–outcome cross-covariance under sample overlap

When an exposure *X* and an outcome *Y* are measured on overlapping mice, their coefficient errors are correlated, and the MR objective requires 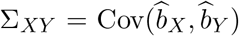, which is zero for independent samples ^17^. Each trait is fitted on its own complete sample, so *n*_*X*_ and *n*_*Y*_ may differ and the two traits may be measured on the same, different or partly overlapping mice; only mice measured for both traits contribute to Σ_*XY*_ . Because R/qtl2 fits each trait separately, we reconstruct Σ_*XY*_ with a qtl2-aligned working model in whitened (star) space rather than a fitted bivariate mixed model.

Each trait is whitened with its own fitted covariance by a symmetric *S*_*T*_ with 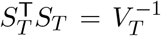, so that 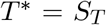 and 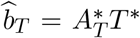 with 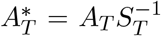. Let *O*_*XY*_ be the *n*_*X*_ *× n*_*Y*_ overlap-matching matrix, with entry 1 where the same mouse appears in both samples and 0 otherwise. Assuming that star-space residuals of overlapping mice share a single cross-correlation *ρ*_∗_,

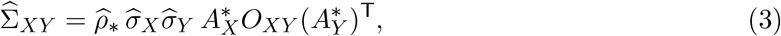

where 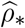 is the correlation of centred star-space residuals across the overlapping mice; when no mouse is shared, 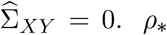 equals a common genetic and environmental correlation only in special cases, such as identical samples with 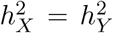 (Supplementary Note 3); in general it is a working scale parameter for the overlap term. Before use, the joint covariance 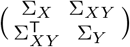 is checked for positive semidefiniteness, equivalently 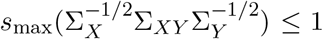 for positive-definite marginals.

### Founder GLS-IVW

For *M* retained loci with summaries 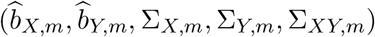 and a candidate scalar effect *θ*, define the founder-contrast residual 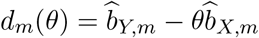 and its covariance

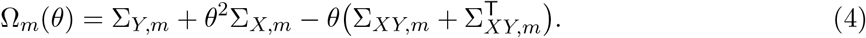

We call this weighting second order (SO) because it propagates uncertainty in both coefficient vectors and includes the overlap cross-term. The Founder GLS-IVW objective and estimator are

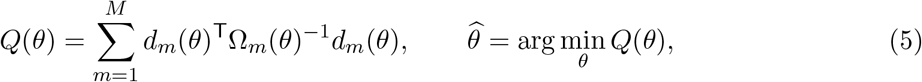

with retained blocks treated as independent across loci, a working assumption (Instrument selection). The same objective is used for one locus (*M* = 1) and for several (global SO), and is minimised over the whole real line (Supplementary Note 1). The founder first-order (FO) estimate, a comparator that treats 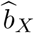 as fixed and ignores Σ_*XY*_, is

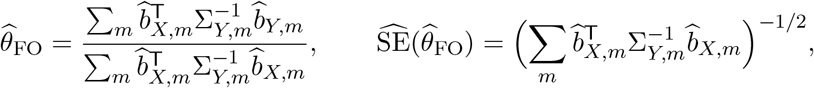

with this fixed-effect standard error and a Wald interval (Supplementary Note 1, Section 1.5). When *b*_*Y,m*_ = *θb*_*X,m*_ for all retained blocks, 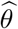 targets *θ*; otherwise it is the single scaling of the exposure founder associations that best fits the outcome founder associations under the SO weighting (simulation S2; Results). The construction parallels summary-data MR with correlated variants ^11,15,42^, and Ω(*θ*) resembles the second-order weights of summary-data MR ^16,43^ and the F-LIML weighting for cis-MR ^36^, a similarity of form rather than an equivalence of estimators.

The primary interval is the nominal 95% profile-*Q* confidence interval,

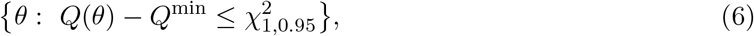

where *Q*^min^ = inf_*θ*_ *Q*(*θ*) over the real line, which equals 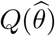 when the minimum is attained at a finite 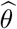 (Supplementary Note 1). It is approximate (Discussion) and need not be symmetric about 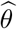. It can also be unbounded: for a single block this occurs when 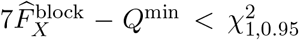, so unbounded sets are expected when the first stage is weak (Supplementary Note 1).

### Diagnostics

#### Founder residuals

At 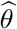, the contrast residual 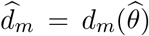 maps to the eight-founder residual 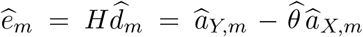, the pattern left after removing the fitted exposure-proportional component. With 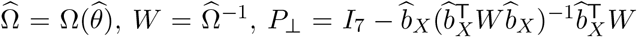 and *v*_*f*_ the contrast with entry 1 for founder *f* and −1*/*7 elsewhere, the one-founder-versus-rest scores

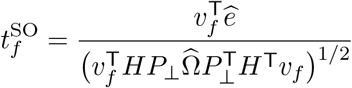

(written *t*_SO_ in Results and figures) are descriptive residual-localisation diagnostics, not causal estimates.

#### Founder-block lack of fit

*Q*^min^, attained at a finite 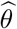 in all reported analyses, measures lack of fit after the best common scaling. For a single locus it asks whether the outcome founder fingerprint is a scalar multiple of the exposure fingerprint, an over-identification question^44,45^, and is referred to an approximate 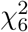 distribution (null calibration, simulation S2). For *M >* 1 it combines within-block non-proportionality and between-locus inconsistency and is referred to an approximate 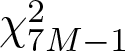 distribution as a nominal diagnostic (Supplementary Note 1).

#### Multi-locus scalar layer

Applying the objective with *M* = 1 to each retained locus gives perlocus SO estimates 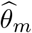, with curvature-based variances (Supplementary Note 1, Section 1.2) and weights 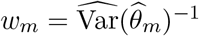, for a forest plot, the two-step IVW estimate and Cochran’s heterogeneity statistic:

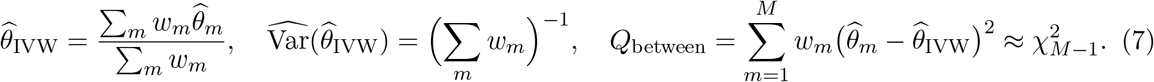

*Q*_between_ targets only between-locus heterogeneity and, with its null calibration not established for weak founder blocks, is reported as an uncalibrated diagnostic. Two-step IVW shares the no-directional-pleiotropy logic of global SO and is not independent evidence. Weighted median ^46^ and weighted mode ^47^ estimates from *{*(*θ*_*m*_, *w*_*m*_)*}* are majority- and plurality-valid sensitivity summaries (intervals, Supplementary Note 5). Leave-one-locus-out refits identify influential loci, and instrument-set refits (excluding one chromosome at a time, or keeping the blocks with the largest first-stage statistics) assess dependence on the instrument set. MR-Egger ^48^ is not used, because a seven-dimensional block has no unique scalar exposure and outcome association.

### Instrument selection

Instruments are local founder blocks at exposure QTL peaks: in the single-locus analysis, the block at the lead marker of the exposure eQTL. For the multi-locus analyses, candidate loci were the autosomal peaks of an upstream genome scan of the exposure (Upstream scan and instrument data), called with the R/qtl2 rule `find peaks`^8^ at a LOD threshold of 4 and a peak drop of 1.5, each represented by its maximum-LOD marker; X-chromosome peaks were excluded (Supplementary Note 5). A locus was removed when its rarest founder allele had an expected count *E* = min_*f*_∑ _*i*_ 2*p*_*if*_ below 5 in the trait’s own complete sample (*p*_*if*_, the probability that a haplotype of mouse *i* derives from founder *f*, half its expected founder dosage), on the exposure’s sample at selection and, for each pair, on the outcome’s sample, where a failing locus was dropped for that pair only. A locus was also removed when more than 5% of the exposure’s sample lacked observed SNP dosage at its marker (Supplementary Note 5). The result was one fixed instrument list per exposure, used by every analysis, with two alternative lists for sensitivity analyses: the peak-drop list (peak drop 2) and the contiguous-run list (Supplementary Note 5). The peak rule reduces, but does not remove, dependence between retained blocks, which equation (5) treats as independent (simulation S4; Discussion).

Exposure strength at a block is summarised by the founder-block first-stage statistic

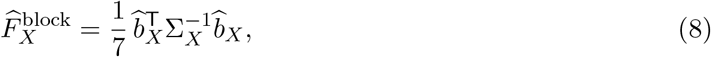

the kinship-adjusted Wald statistic for *b*_*X*_ = 0 per contrast direction (written 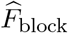 in Results and figures), a descriptive average that reduces to the usual first-stage *F* statistic for a single scalar coefficient^49,50^. It was computed once, from the exposure’s own-sample fit, and used unchanged. In the multi-locus analyses, blocks with 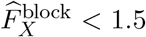 were excluded before stacking (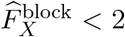 in sensitivity analyses), a lenient inclusion threshold rather than a test of relevance (Supplementary Note 5). Where stated, loci also satisfied the Steiger condition 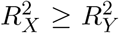 within the pair ^51^, with 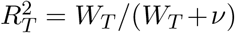 the partial *R*^2^ from the block Wald statistic 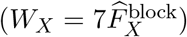 and *ν* the residual degrees of freedom.

Scalar comparators collapse the founder block to one genotype score: an observed marker dosage (0/1/2), a founder-aware variant dosage or, in simulation, *g*_*s*_ = *Ps* for a binary founder grouping *s* ∈ *{*0, 1*}*^8^. One-dimensional kinship-adjusted mixed models give scalar coefficients 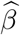 with variances *v*, the first-stage statistic 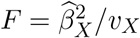 (*χ*^2^ reference) and the single-locus Wald ratio 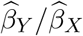. Across loci, the scalar estimate (S in the landscape) is the summary-data SNP inverse-variance-weighted (IVW) estimate of human MR,

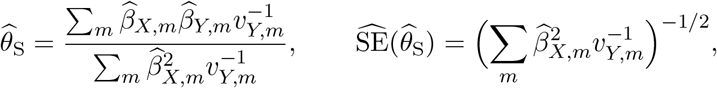

with this fixed-effect standard error and a Wald interval; like a conventional IVW analysis, it ignores the uncertainty in 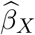 and any sample overlap.

### Simulation design

Parameters, replicate numbers and data-generating equations are given in Supplementary Note 4 and Supplementary Table 1. Instrument strength was set by 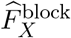, the population founder-block statistic of the design, not the fitted *F* ^block^, and sample overlap by a residual correlation on shared mice, giving an effective coefficient-error correlation *ρ*_eff_ .

#### Single locus (S1–S3)

These kept the genotypes, kinship and covariates of real mice from the *Qsox1* analysis (*n* = 400 at `JAX00272363`; for S3, *n* = 500 at `UNC494697`), simulated founder effects and residuals from the covariance model used in fitting, with outcome founder effects *θb*_*X*_ + *δ*, and refitted each replicate with the qtl2-compatible model. S1 tested calibration under proportionality (*δ* = 0) against first-order intervals, with a synthetic-design check of the overlap-aware and independence (Σ_*XY*_ = 0) objectives and a check over *h*^2^. S2 added a contamination *δ* of relative magnitude *λ* and alignment *κ* with the exposure pattern (*κ* = 0 orthogonal, *κ* = 1 aligned), recording bias, coverage of *θ* and of the projection slope *θ*^⋆^ (which adds to *θ* the part of *δ* aligned with *b*_*X*_) and *Q*^min^ rejection (null calibration at *λ* = 0). S3 collapsed one non-proportional architecture under four binary codings *g*_*s*_ = *P*_*m*_*s*, comparing scalar *F* statistics and Wald ratios with their analytical estimands (Supplementary Note 1, Section 1.7).

#### Multiple loci (S4, S5)

S4 had three arms: independent synthetic blocks; summary-level blocks with known sample-overlap covariance, comparing the independence and overlap-aware objectives for up to 500 blocks; and correlated neighbouring synthetic blocks, comparing an unpruned block-diagonal analysis with one pre-specified representative block (the ‘pruned top hit’) and a full-covariance comparator. S5 used summary-level blocks for *M* = 20 loci, clean or with two high-information invalid loci (shifted but proportional effects, orthogonal contamination or both), and evaluated the multi-locus estimators and diagnostics. The replicate shown in Fig. 3g was, among replicates in which *Q*_between_ rejected at *P <* 0.05 and the top leave-one-out locus was a planted invalid locus, the one whose global-estimate error was closest to the scenario median.

#### Selection, coverage and performance

Main results use all replicates and blocks; at *F*_target_ = 1.5 the multi-locus analyses were repeated after dropping blocks with 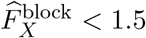. Coverage is exact membership of the profile-*Q* set, which can be unbounded or disconnected: *θ* is covered when 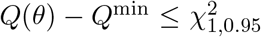, with *Q*^min^ including the limit as |*θ*| → ∞. We report relative error, mean relative bias, coverage and *Q* rejection rates, with Monte Carlo or binomial 95% intervals ^52^.

### Upstream scan and instrument data

All real-data analyses used the Farber laboratory DO bone cohort ^24^. For the multi-locus analyses, we used that laboratory’s R/qtl2 genome scan of the cohort’s traits, supplied with the cohort data, only to select instruments: its peaks defined the candidate loci for every exposure except body weight and body length (Instrument selection). Because that scan adjusted for body weight and body length, we rescanned these two traits without those covariates; all downstream estimates, including instrument strength, founder support and every MR effect, come from our own mixed models (Supplementary Note 5). Within our pipeline, R/qtl2^8^ provides the genome scans (`scan1`), the peak rule (`find_peaks`) and the founder-coefficient model (`scan1coef`) that the full-covariance fitter reproduces, and lme4qtl^53^ provides the variance components of the scalar SNP fits.

### Real-data analyses

#### Data and ethic

The *Qsox1* analysis used the published R/qtl2 cross files of the cohort (Zenodo record 4265417): 619 phenotyped mice, 192 with cortical-bone RNA-seq. The ML–pMOI and landscape analyses used additional Farber laboratory genotype and phenotype data, including animals not in those files. Leave-one-chromosome-out (LOCO) kinship matrices came from the cohort’s genotype data, subset to the mice analysed for each trait. This is a secondary analysis of existing data ^24^; no new animal experiments were performed.

#### Traits and processing

Bone-trait names and abbreviations ^54^ are listed in Supplementary Note 5. Each trait was analysed on its own complete sample: the mice with a finite value for that trait and complete covariates that were also present in the kinship and founder-probability data. Phenotypes and expression were z-standardised (in the *Qsox1* analysis on all mice with an observed value, otherwise on each trait’s own complete sample), so that estimates are in outcome standard deviations (SD) per exposure SD; the four marrow adipose tissue (MAT) volumes, zeros included, were instead rank inverse-normal transformed on each trait’s own complete sample (Supplementary Note 5). In the ML–pMOI and landscape analyses, every fit adjusted for sex, age at sacrifice and generation as a categorical factor (indicators for generations 24–35, with 23 as reference), with no body-size covariate. Two features of the data were left unchanged: in one generation the maximum and minimum moments of inertia (Imax and Imin) do not sum to pMOI as they should, so landscape counts are also given excluding the 290 pairs that involve Imax or Imin; and 20 mice with pMOI but no ML were kept, so the ML and pMOI samples overlap only partly.

#### Single-locus analysis at Qsox1

The exposure was *Qsox1* expression in 192 mice, adjusted for sex and 48 probabilistic estimation of expression residuals (PEER) factors ^55^ (processing, Supplementary Note 5). The outcomes were eight femoral traits in 611 mice (591 for medial–lateral femoral width, ML), 187 (169 for ML) of them with expression data, adjusted for sex, age at sacrifice, body weight and generation indicators (generations 24–33, with 23 as reference). Adjusting the outcomes for body weight and the exposure for PEER factors were deliberate choices for this single-locus design; the multi-locus analyses use neither. All fits used the chromosome 1 LOCO kinship matrix ^56^, with *h*^2^ estimated by REML for each trait and held fixed (Supplementary Table 2(a)). The instrument was the founder block at the lead *Qsox1* eQTL marker `JAX00272363` (Supplementary Note 5; Extended Data Fig. 1). Founder GLS-IVW with *M* = 1 gave 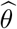, the profile-*Q* interval, *Q*^min^ and 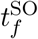. Scalar comparators were the GigaMUGA^25^ dosage at `JAX00272363` and the founder-aware scan1snps dosage at `rs232318063`.

#### Multi-locus analysis of ML and pMOI

The exposure was ML and the outcome the polar moment of inertia (pMOI), fitted on 779 and 799 mice; Σ_*XY,m*_ was built from the 779 measured for both. Each block used its chromosome’s LOCO kinship matrix, with *h*^2^ estimated by REML for each trait and chromosome and held fixed (Supplementary Table 2(b)). The primary analysis used the 55 of the 57 ML loci on the fixed list with 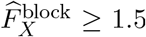, without the Steiger condition; all 55 joint covariances were positive semidefinite and none was adjusted (Supplementary Table 7(b)). We report the global SO estimate with its profile-*Q* interval, the diagnostics above (refits, Supplementary Note 5) and a fit with Σ_*XY*_ = 0 and a Wald interval. Sensitivity analyses used 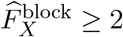 (49 loci), the Steiger condition and the peak-drop and contiguous-run lists; the reverse direction, pMOI to ML, used the pMOI instruments (45 loci with 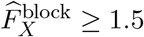). Comparators were the scalar SNP-IVW at the 55 loci and an ordinary least-squares regression of pMOI on ML in the 779 mice measured for both, with the covariates of the MR fits. All are reported in Supplementary Table 9.

#### Landscape of trait pairs

The landscape used 74 traits (Supplementary Note 5): the 82 traits of the cohort’s genome scan, less eight derived binary and non-zero indicators that duplicate the four MAT volumes. A trait served as an exposure if it had at least 10 primary instruments with 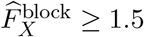; all 74 did, body weight and body length after their own scan. Every trait also served as an outcome, giving 74 *×* 73 = 5,402 planned ordered pairs, each fitted as above.

For each pair, three analyses used the same loci, the set *L*: the exposure’s primary instruments with 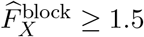, less any locus dropped for that pair by outcome-side founder support, restricted to loci satisfying the Steiger condition in the pair. S is the scalar SNP-IVW (SNP effects, Supplementary Note 5); F1 is 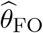 with its Wald interval; F2 is the global SO estimate with its profile-*Q* interval. All three are fixed-effect analyses: no interval is inflated for between-locus heterogeneity. The representation step, S→F1, replaces one observed SNP dosage per locus by the seven founder contrasts, with the same loci, samples, covariates and kinship; both analyses are first order and ignore sample overlap. The overlap-aware step, F1→F2, replaces the first-order estimator and Wald interval by the second-order estimator, which propagates the uncertainty in 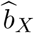 and includes Σ_*XY*_, and its profile-*Q* interval. All founder analyses of a pair came from one fit per exposure (Supplementary Note 5).

An interval excludes zero when both bounds lie on the same side of zero. Each pair in the QC cohort was assigned one of five endpoint states by whether its S and F2 intervals exclude zero (only F2; only S; neither; both, with the same sign; both, with opposite signs), and likewise for (S, F1) and (F1, F2). At each step, a gain is a change from including to excluding zero and a loss the reverse; a pair can gain at one step and lose at the other. Sensitivity analyses repeated the classification on the primary QC cohort with 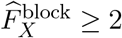 and with the peak-drop and contiguous-run lists (Supplementary Table 3; Supplementary Note 5).

### From planned pairs to the landscape QC cohort

Of the 74 *×* 73 = 5,402 planned ordered pairs, 4,596 form the quality-control (QC) cohort analysed in Results; the other 806 were removed by a scalar-estimability check and by QC. No pair was lost in the founder fits: none returned an error or had an empty instrument set, and every pair had successful fits on *L* and without a threshold. However, 132 pairs were not estimable in the scalar analysis, because at least one locus of *L* had no SNP effect for the exposure or the outcome (its marker had a minor allele frequency below 0.01 in that trait’s own sample). These pairs were removed whole rather than by dropping the locus, so that S, F1 and F2 keep the same loci, leaving 5,270 classifiable pairs. QC used the global SO estimate at inclusion thresholds of 0, 1.5 and 2 without the Steiger condition, and at 1.5 and 2 with it. A pair was not reportable (class A) unless its fits at thresholds 1.5 and 2 without the Steiger condition succeeded with at least 10 and 5 loci, respectively; weak-locus sensitive (class B) if the estimate at threshold 1.5 or 2 changed sign or moved by more than 0.25 from the estimate without a threshold; and Steiger sensitive (class C) if, at either threshold, adding the Steiger condition made the fit fail, changed the sign of the estimate or moved it by more than 0.25. Each pair was counted once, in the first class it met in the order A, B, C: none was in class A, 402 in class B and 272 in class C. All 4,596 remaining pairs gave a finite F2 estimate inside finite, ordered profile bounds and form the QC cohort (4,349 of 5,112 pairs when the 290 pairs involving Imax or Imin are excluded). Because QC requires stability across thresholds, the 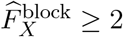 sensitivity analysis is not independent of the QC cohort.

### Basis-rotation decomposition

The follow-up covered the 170 scalar-only pairs, the pair with opposite-sign estimates and random samples of 100 pairs significant under both S and F2 and of 100 founder-only pairs. At each locus whose marker has a complete founder strain distribution, the founder coefficients of both traits were whitened with the first-order outcome precision 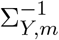 and rotated so that the first axis, the SNP axis, lies along the marker’s strain-distribution contrast and the other six span the founder directions orthogonal to it in this metric; for the outcome, the first coordinate is the marker regression within the founder model, which the S analysis approximates. The rotation leaves the first- and second-order estimates unchanged, and on these loci the first-order estimate, unlike the second-order estimate, is an information-weighted average of the estimates from the SNP axis and from the other six directions. The noise-corrected SNP-axis share of first-order information is 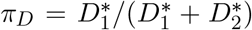, where 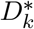 is the squared whitened exposure effect in part *k*, summed over loci, less its expected noise. Agreement between exposure and outcome effects within each part was summarised by the concordance index *r*_*k*_, a cosine between the two founder-effect patterns corrected for estimation noise and sample overlap (1, proportional; 0, unrelated; the correction allows values above 1). Overlap amplification (OA) marks pairs whose SNP-axis interval excludes zero when overlap is ignored but whose overlap-aware profile set contains zero. Rules were fixed in advance; two deviations and one added flag are stated in Supplementary Note 6 (values, Supplementary Table 10).

### Software

All founder-based analyses (Founder GLS-IVW and its diagnostics, including the two-step IVW, weighted-median and weighted-mode estimators) and simulation S5 used qtl2mr 0.7.0. Observational estimates were ordinary least-squares regressions, simulations S1–S4 were generated and analysed with the simulation engine (Supplementary Note 4), and the roles of R/qtl2 and lme4qtl are described under Upstream scan and instrument data. The single-locus scalar Wald ratios at *Qsox1* were computed from R/qtl2 associations. Validation of the full-covariance fitter, invariance to the contrast basis and validity of the joint covariance in the real-data analyses, and the accuracy of the star-space reconstruction in simulation, are reported in Supplementary Tables 4–7 (Supplementary Notes 2–4).

### Statistics

All confidence intervals are nominal 95% and two-sided, and tests of *θ* = 0 are two-sided. The primary Founder GLS-IVW interval is the profile-*Q* interval (equation (6)). Heterogeneity statistics use the upper tail of the approximate *χ*^2^ references given under Diagnostics, and all *P* values are nominal. *Q*^min^ *P* values across the eight correlated *Qsox1* bone phenotypes are diagnostic references, not independent multiple-testing claims, and founder residual scores with 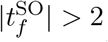 are a descriptive guide, not a corrected threshold. Landscape summaries are descriptive: no paired estimator-difference tests were performed, and no analysis is treated as a truth benchmark.

### Use of large language models

OpenAI Codex (GPT-5- and GPT-6.1-series models) and Claude Code (Anthropic; Claude Opus 5.5) were used by X.D. for code development and review, running and checking analyses on a high-performance computing cluster under the author’s direction, independent review of results, drafting and editing text, and figure and L^A^TEX layout. X.D. directed this work, reviewed all AI-assisted material and checked the software, analyses, numerical results, citations and figures. The authors take responsibility for the final manuscript.

## Data availability

The published R/qtl2 cross files of the Farber laboratory DO bone cohort used in the *Qsox1* analysis are available from Zenodo (https://doi.org/10.5281/zenodo.4265417), and the corresponding expression data from the Gene Expression Omnibus (GSE152708 and GSE152806)^24^. Other geno-type and phenotype data from the Farber laboratory DO bone cohort analysed in this study are held by the Farber laboratory; further releases of these data will be made available by that laboratory.

## Code availability

Founder GLS-IVW is implemented in the R package qtl2mr, which is publicly available (version 0.7.0; https://github.com/XDai-bris/qtl2mr-uvmr). The scripts that reproduce the analyses and figures are available to editors and reviewers on request and will be made public on publication.

## Acknowledgements

We thank Saikat Bandyopadhyay for transferring the DO bone cohort data and for help with their interpretation, and Laurence Howe for comments on the manuscript. This work was carried out using the computational facilities of the Advanced Computing Research Centre, University of Bristol (https://www.bristol.ac.uk/acrc/).

## Author contributions

X.D. conceived the study, developed the method and software, performed the simulations and analyses, and wrote the manuscript. G.H. and G.D.S. supervised the work, contributed to the development of the method and to the interpretation of the results, and commented on the manuscript. G.C. contributed to the founder-haplotype framework and to the interpretation of the results, commented on the manuscript and contributed to its writing. L.D.M., G.M.C. and C.R.F. generated the DO bone cohort data and contributed to their interpretation. All authors reviewed and approved the final manuscript.

## Extended Data figure legends

**Extended Data Fig. 1.**
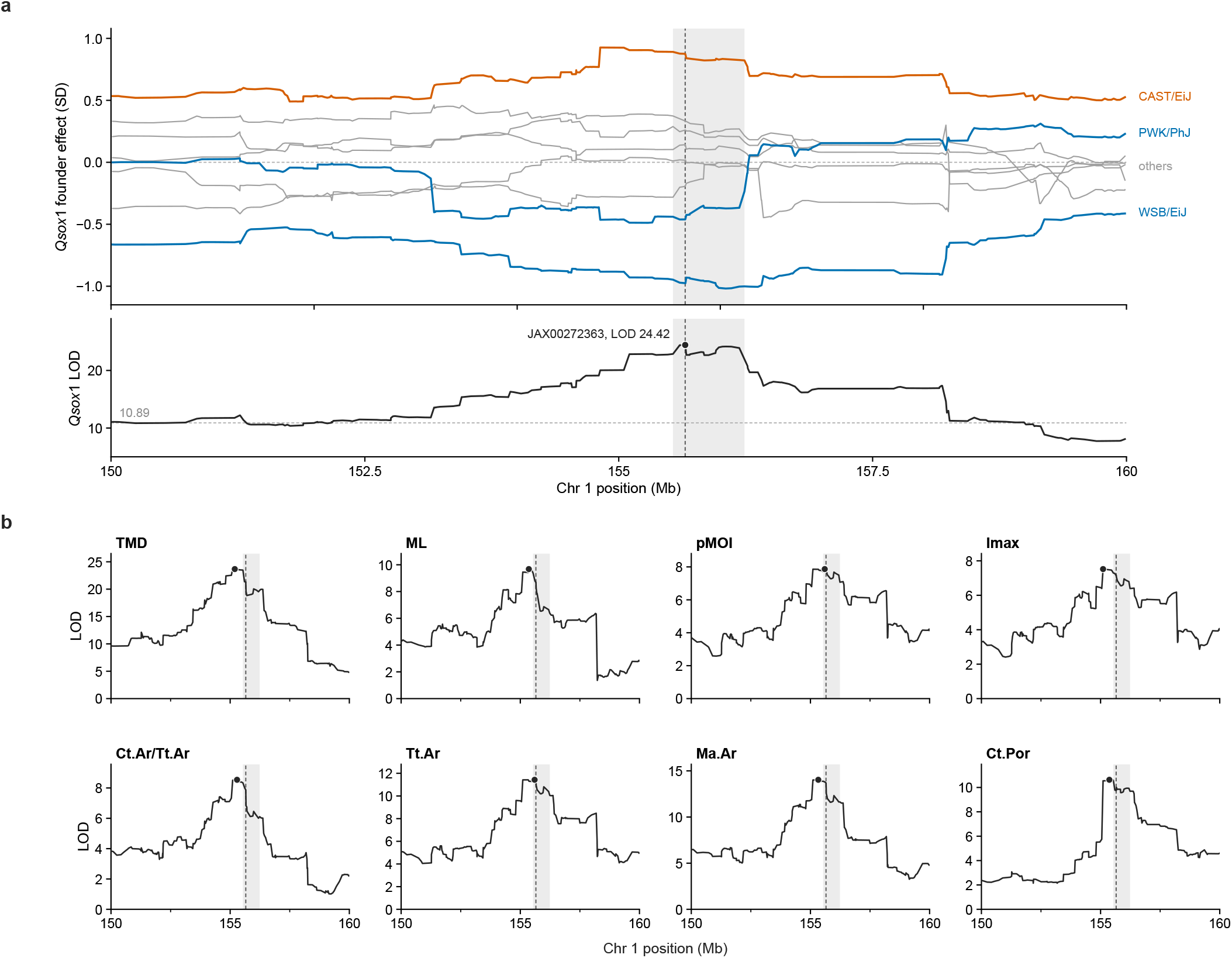
*Qsox1* founder eQTL and bone-trait QTL on chromosome 1. **a**, Top, founder effects on standardised *Qsox1* expression (expression SD) along chromosome 1 from 150 to 160 Mb, estimated as best linear unbiased predictors (R/qtl2 scan1blup). These shrunken estimates are smaller than the coefficients in Fig. 2a. CAST/EiJ is shown in vermillion, PWK/PhJ and WSB/EiJ in blue and the other founders in grey. Bottom, founder-probability eQTL LOD scores for *Qsox1* (*n* = 192 mice with cortical-bone RNA-seq). The point marks the lead marker JAX00272363 (155.654 Mb; LOD 24.42), and the horizontal dashed line marks the autosomal eQTL permutation threshold of the original study (LOD 10.89) ^24^. **b**, Founder-probability QTL LOD scans of the eight bone traits analysed at this locus (published cross; up to 619 mice). Points mark each trait’s lead marker. In **a** and **b**, the grey band is the *Qsox1* eQTL support interval reported by the original study (155.53– 156.24 Mb), and the dashed vertical line marks the eQTL lead marker. Overlapping peaks motivate the founder-block analysis but do not test colocalisation or mediation. TMD, cortical tissue mineral density; ML, medial–lateral femoral width; pMOI, polar moment of inertia; Imax, maximum moment of inertia; Ct.Ar/Tt.Ar, cortical area fraction; Tt.Ar, total area; Ma.Ar, medullary area; Ct.Por, cortical porosity.

**Extended Data Fig. 2.**
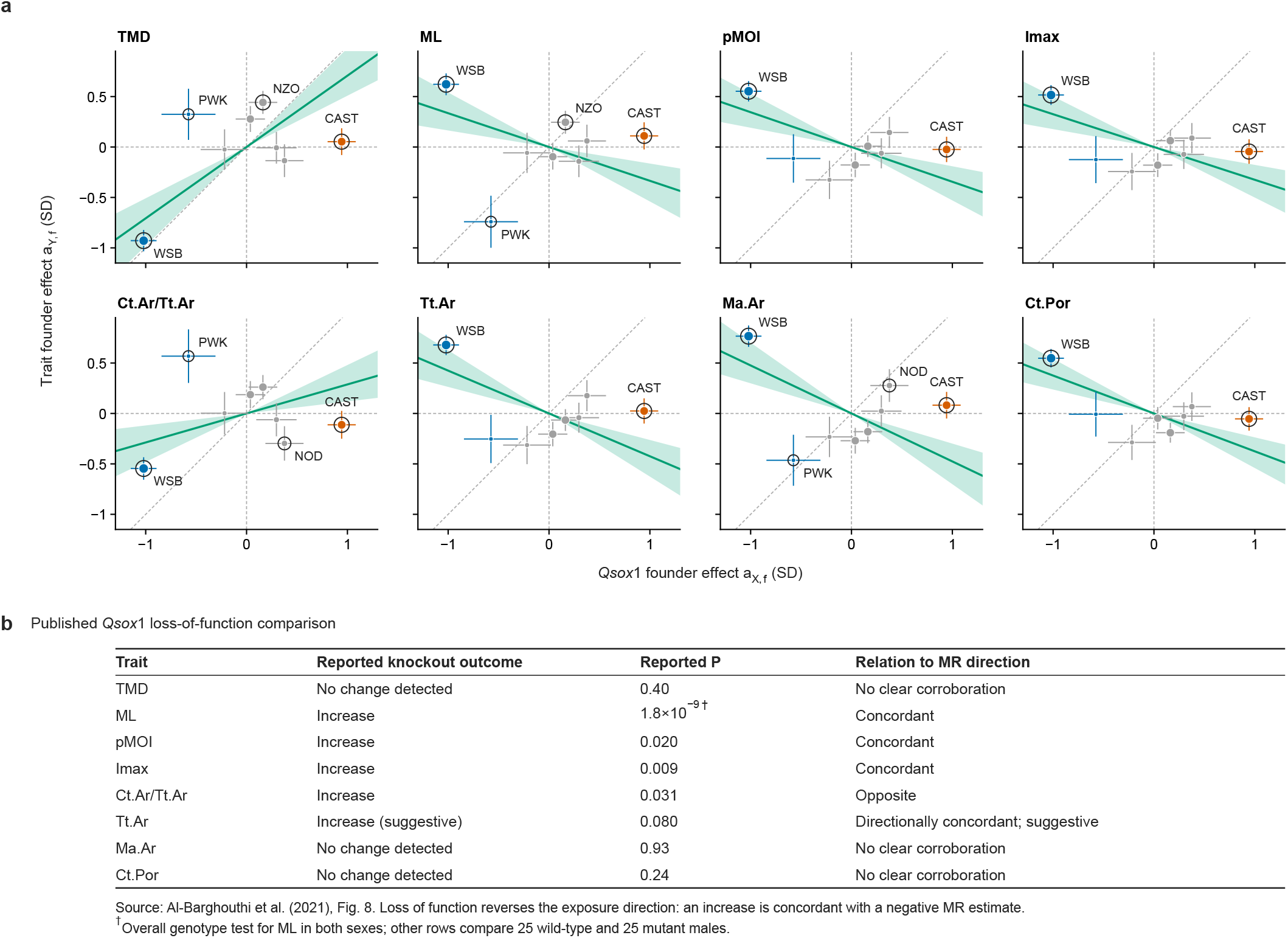
*Qsox1* and eight bone traits in founder space, with the published knockout outcomes. **a**, Mean-centred founder effects on each bone trait (*a*_Y,f_, trait SD) plotted against founder effects on *Qsox1* expression (*a*_X,f_, expression SD), in the encoding of Fig. 2d. The exposure fit is the same in every panel. Crosshairs show *±*1 marginal s.e., and point area is proportional to 1/Var(*a* _X,f_). The green line is the overlap-aware second-order (SO) Founder GLS-IVW estimate 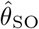 through the origin, and the green tint is its nominal profile-*Q* 95% CI. Dashed lines mark *x* = 0, *y* = 0 and *y* = *x*. Rings mark founders whose one-founder-versus-rest residual has |*t* _SO_| *>* 2, a descriptive guide rather than a formal threshold. CAST/EiJ is shown in vermillion, PWK/PhJ and WSB/EiJ in blue and the other founders in grey. All panels share axes and have equal aspect. Founder labels are abbreviated (CAST, CAST/EiJ; NOD, NOD/ShiLtJ; NZO, NZO/HlLtJ; PWK, PWK/PhJ; WSB, WSB/EiJ). **b**, Comparison with published *Qsox1* loss-of-function experiments ^24^. Five CRISPR mutant lines lacked serum QSOX1 activity. *µ*CT traits were measured in 25 wild-type and 25 mutant male F2 mice on B6/SJL-derived backgrounds; ML was measured in a larger sample of both sexes. Reported *P* values are reproduced without reanalysis: the ML value is the overall genotype test in the sex-combined model, whereas the others are contrast *P* values comparing wild-type and mutant mice, adjusted for multiple comparisons in the original study. An increase after loss of function is directionally concordant with a negative expression–trait estimate. A non-significant experimental result is labelled as no detected change, not evidence of no effect. The comparison concerns direction, not the magnitude of the MR estimate. Exposure *n* = 192. Outcome *n* = 611 for all traits except ML (591); 187 of these mice overlap with the exposure sample (169 for ML). Estimates, *Q* ^min^ and the largest founder residual for each trait are given in Fig. 2e. Trait abbreviations are as in Extended Data Fig. 1.

**Extended Data Fig. 3.**
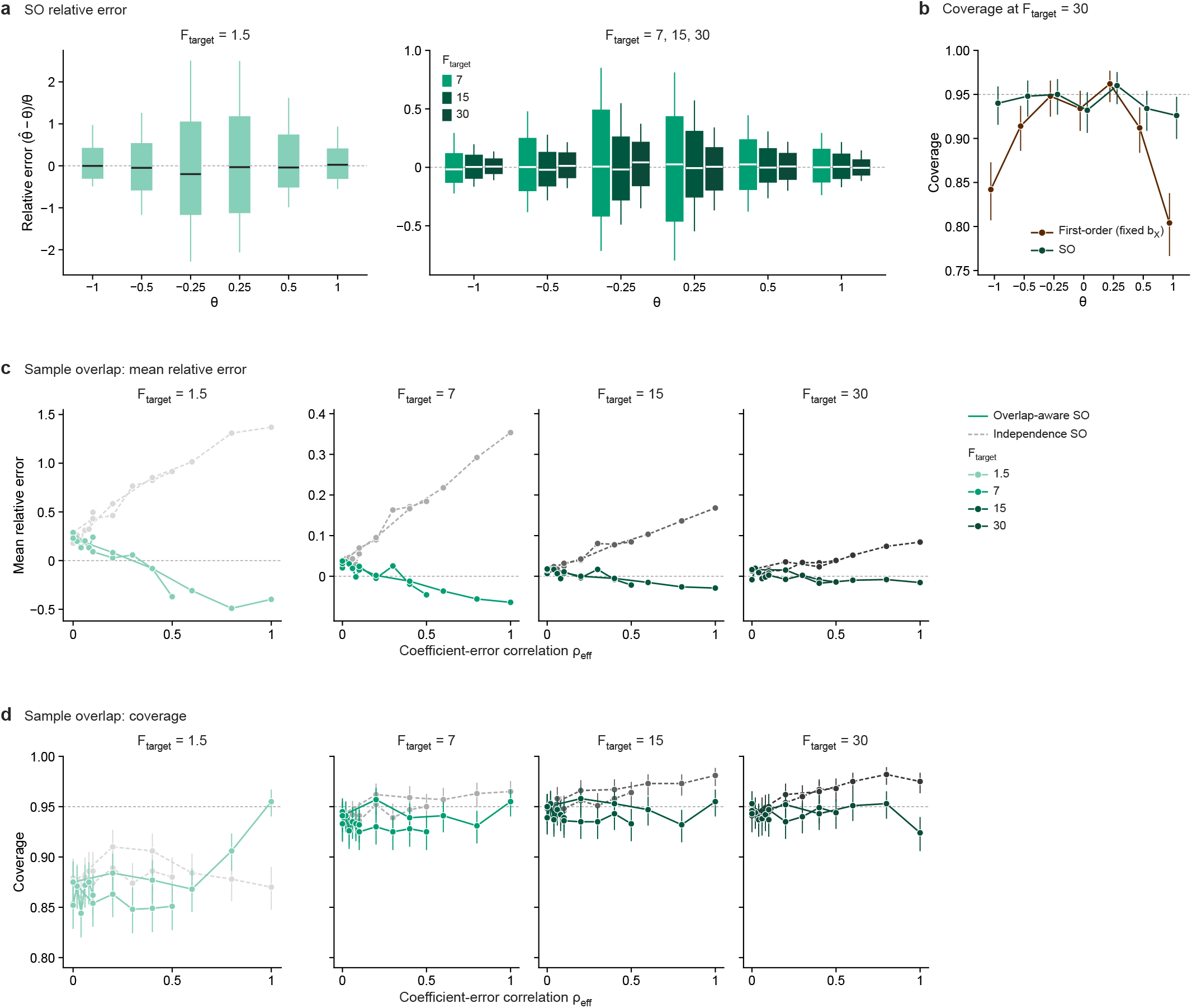
Single-locus simulations (S1): estimates, coverage at *F* _target_ = 30 and sample overlap. All replicates are shown; shade gives *F* _target_, from light to dark 1.5, 7, 15 and 30. **a**, Relative error 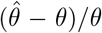 of the second-order (SO) estimate (Σ_XY_ = 0, its true value under independent residuals) by *θ* (*θ* ≠ 0; 500 replicates each); *F* _target_ = 1.5 has its own scale. Boxes, interquartile range; whiskers, 10th–90th percentiles; lines, medians. **b**, Coverage at *F* _target_ = 30 of first-order Wald intervals (fixed *b*_X_) and SO profile-*Q* sets. **c**,**d**, Single-locus overlap check in a synthetic design (400 exposure and 400 outcome mice, simulated kinship and founder probabilities; *θ* = 0.25, 1,000 replicates per setting): mean relative error (**c**) and coverage (**d**) of the overlap-aware (green, solid lines) and independence (grey, dashed lines) SO objectives against the effective coefficient-error correlation *ρ*_eff_, the product of the overlap fraction (0.1, 0.5 or 1) and the residual correlation (0–1). Lines join settings with the same overlap fraction. Coverage is exact profile-set coverage (Fig. 3); at *F* _target_ = 1.5 the SO set is unbounded in 11–15% of S1 replicates and 3–15% of overlap-check replicates, and in none at *F* _target_ ≥ 7. Bars, Clopper–Pearson 95% CIs. Dashed grey lines, zero error or 0.95 coverage. Detailed numbers are in Supplementary Note 4.

**Extended Data Fig. 4.**
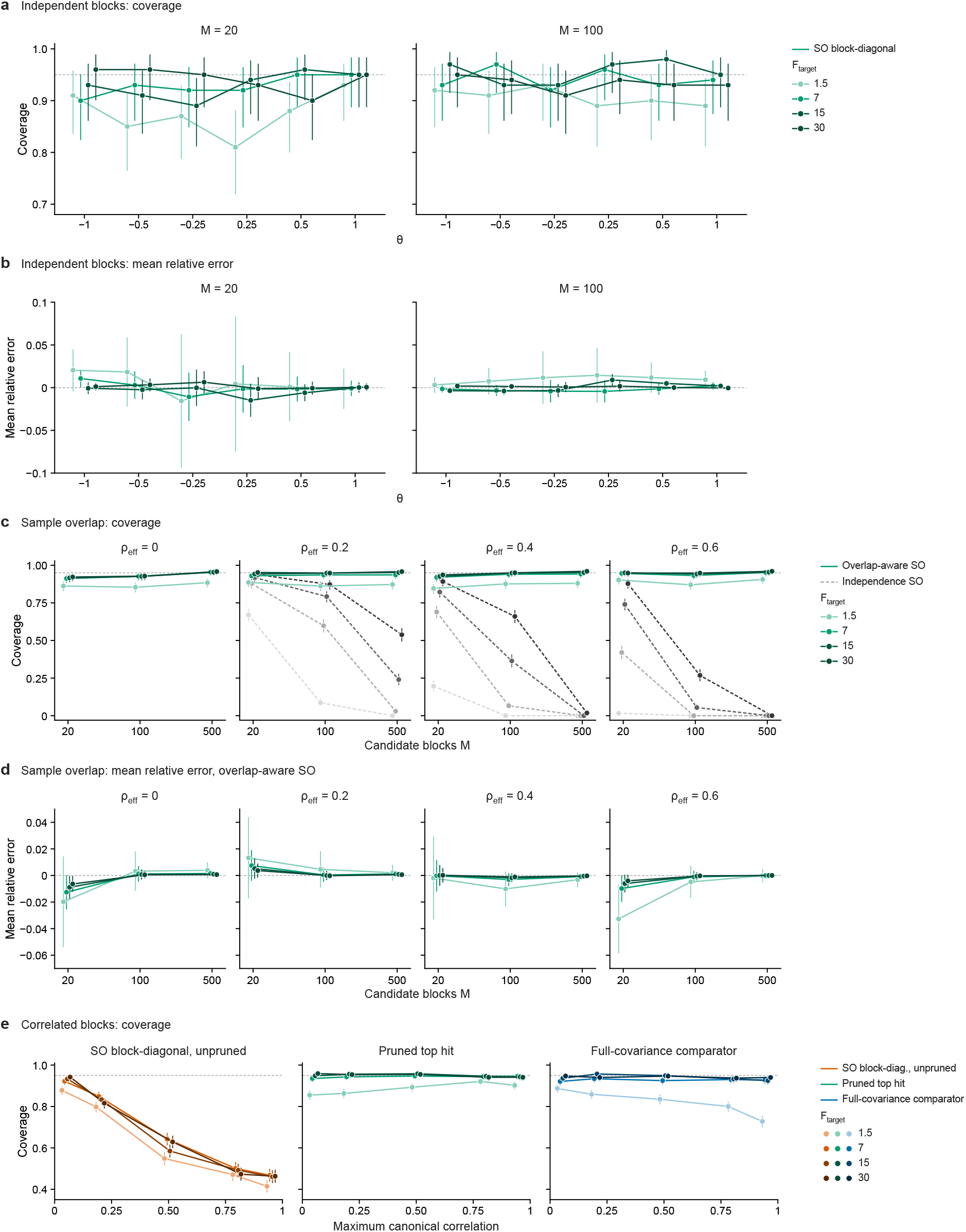
Multi-locus simulations (S4): independent blocks, sample overlap and correlated blocks. All candidate blocks are stacked; within each hue, shade gives *F* _target_, from light to dark 1.5, 7, 15 and 30. **a**,**b**, Independent-block arm (synthetic *n* = 900 designs refitted by GLS; *M* = 20 or 100 blocks; 100 replicates per *θ*): coverage (**a**) and mean relative error (**b**; bars, 95% CI of the mean) of the SO block-diagonal analysis. **c**,**d**, Sample-overlap arm (coefficient-summary blocks, *θ* = 0.25, 500 replicates): coverage of the overlap-aware (green, solid lines) and independence (grey, dashed lines) objectives (**c**) and mean relative error of the overlap-aware objective (**d**; bars as in **b**) against the number of candidate blocks *M*, at effective coefficient-error correlation *ρ*_eff_ = 0–0.6. At *ρ*_eff_ = 0 the two objectives coincide. **e**, Correlated-block arm (*M* = 10, *n* = 3,000, *θ* = 0.25, 1,000 replicates): coverage against the observed maximum canonical correlation for the unpruned SO block-diagonal analysis (orange), the pruned top hit (green) and the full-covariance comparator (blue). Coverage is exact profile-set coverage (Fig. 3); at *F* _target_ = 1.5 the pruned-top-hit set is unbounded in up to 15% of replicates and every other set in at most 1%. Bars in **a, c** and **e**, Clopper–Pearson 95% CIs. Dashed grey lines, 0.95 coverage or zero error. Detailed numbers are in Supplementary Note 4.

**Extended Data Fig. 5.**
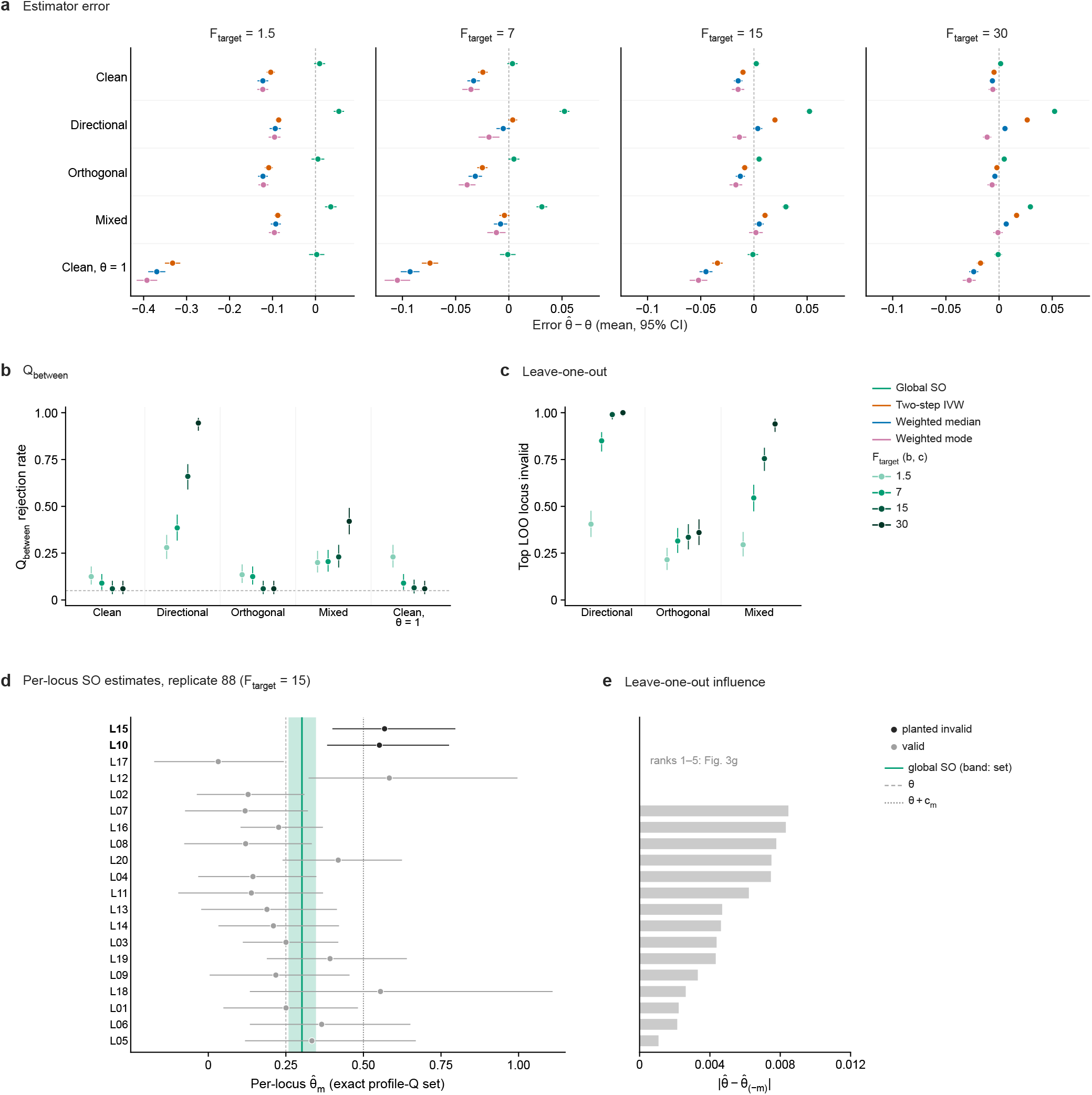
Cross-locus diagnostics (S5) across scenarios and *F* _target_ levels. Each replicate stacks all *M* = 20 candidate blocks, every locus at the same *F* _target_, with *θ* = 0.25 (200 replicates per scenario). Scenarios: Clean, all loci proportional; Directional, the two highest-information loci have *b*_Y_ = (*θ* + *c*_m_)*b*_X_ with *c*_m_ = *θ*; Orthogonal, the same two loci carry a component orthogonal to *b*_X_; Mixed, *c*_m_ = *θ*/2 plus an orthogonal component; Clean, *θ* = 1, as Clean with *θ* = 1. **a**, Error 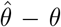 (mean and 95% CI) of the global SO, two-step IVW, weighted-median and weighted-mode estimates. **b**, *Q* _between_ rejection rate (*α* = 0.05). **c**, Rate at which the locus with the largest leave-one-out influence is a planted invalid locus. In **b** and **c**, shade gives *F* _target_, from light to dark 1.5, 7, 15 and 30; bars, Clopper–Pearson 95% CIs; dashed grey line, 0.05. **d**,**e**, Representative replicate 88 of the directional scenario at *F* _target_ = 15 (Fig. 3g), loci ordered by leave-one-out influence: per-locus SO estimates with exact profile-*Q* sets (**d**; black, planted invalid loci; green line and band, global SO estimate 0.302 and its set 0.258–0.347; dashed, *θ*; dotted, *θ* + *c*_m_) and influence 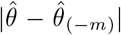 for ranks 6–20 (**e**; ranks 1–5 in Fig. 3g). Detailed numbers are in Supplementary Note 4.

**Extended Data Fig. 6.**
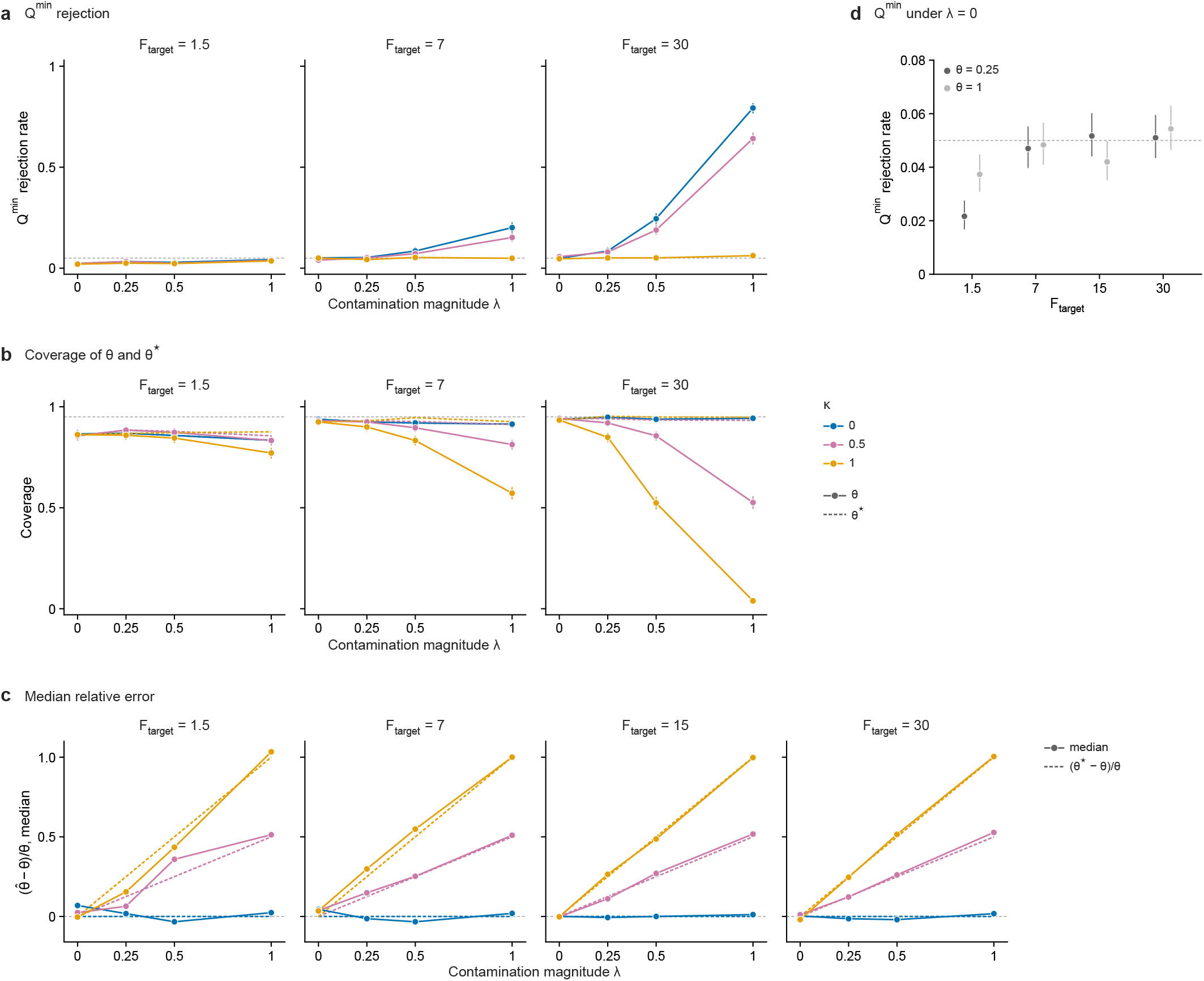
Founder-block contamination (S2) at further *F* _target_ levels. Design as in Fig. 3c,d (JAX00272363, *n* = 400, *θ* = 0.25, full overlap with residual correlation 0.2, 1,000 replicates per setting, all shown), with contamination magnitude *λ* and exposure alignment *κ* (hues). **a**, *Q* ^min^ rejection rate (*α* = 0.05, approximate *χ*^2^_6_ reference) at *F* _target_ = 1.5, 7 and 30 (15 is in Fig. 3). **b**, Coverage of *θ* (solid) and of the projection slope *θ*^⋆^ (dashed; Methods). **c**, Median relative error 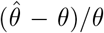, also at *F* _target_ = 15; dashed lines, the shift of the projection slope, (*θ*^⋆^ − *θ*)/*θ*. The median is shown because the mean is heavy-tailed at *F* _target_ = 1.5. **d**, *Q* ^min^ rejection under *λ* = 0 (*κ* pooled, 3,000 replicates) by *F* _target_, for *θ* = 0.25 (dark grey) and *θ* = 1 (light grey). Coverage is exact profile-set coverage (Fig. 3); at *F* _target_ = 1.5 the set is unbounded in 11–16% of replicates per setting, and in at most 0.1% at *F* _target_ ≥ 7. Bars, Clopper–Pearson 95% CIs. Dashed grey lines, 0.05 rejection or 0.95 coverage. Detailed numbers are in Supplementary Note 4.

**Extended Data Fig. 7.**
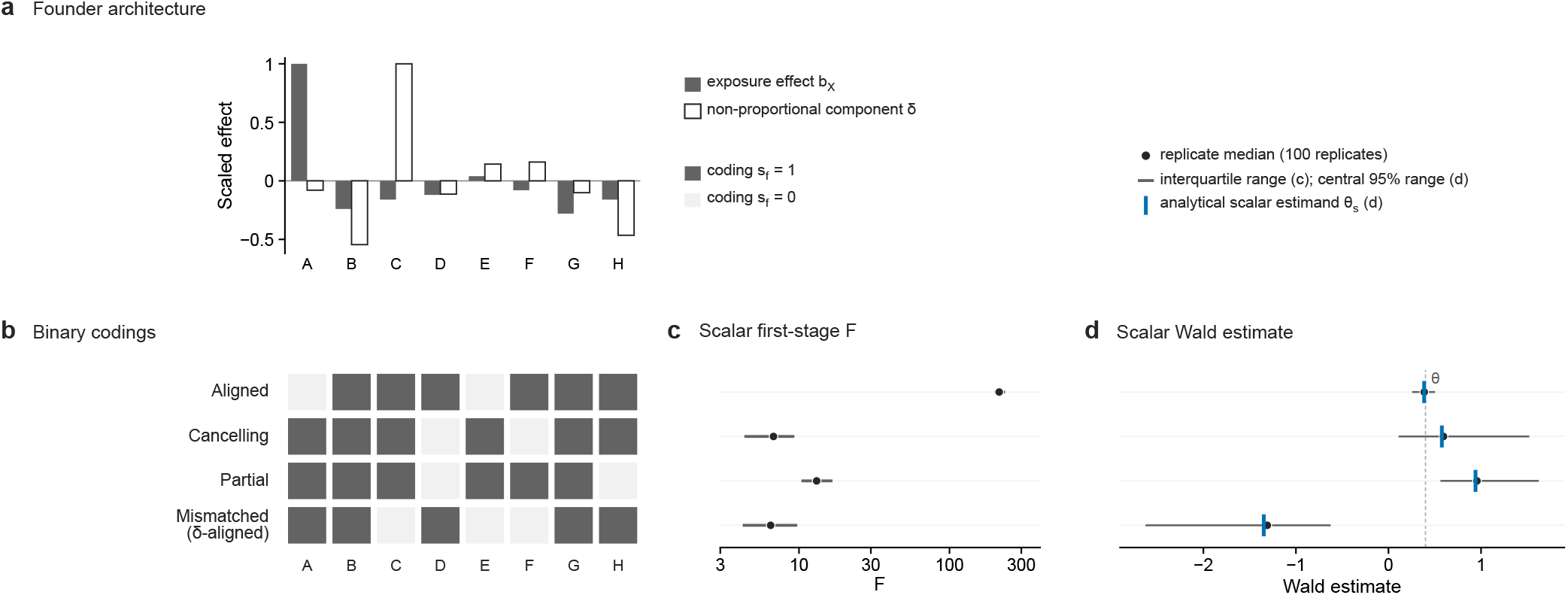
Scalar-coding simulation (S3): design and scalar estimates. S3 uses the empirical Diversity Outbred design at UNC494697 (*n* = 500), exposure founder effects scaled to *F* _target_ = 80 and a non-proportional outcome architecture *b*_Y_ = *θb*_X_ + *δ* with *θ* = 0.4, where *δ* is orthogonal to *b*_X_ in founder-contrast space (correlation 0.0003). **a**, Founder effects *b*_X_ (filled) and *δ* (open), each scaled to a maximum absolute value of 1 (unscaled maxima: *b*_X_ 3.58 at founder A; *δ* 1.09 at founder C). Founders: A, A/J; B, C57BL/6J; C, 129S1/SvImJ; D, NOD/ShiLtJ; E, NZO/HlLtJ; F, CAST/EiJ; G, PWK/PhJ; H, WSB/EiJ. **b**, The four binary codings *s* that collapse the founders into a scalar genotype *g*_s_ = *P* _m_*s* (filled, *s*_f_ = 1). Complementing a coding changes neither *F* nor the Wald ratio. **c**, Scalar first-stage *F* over 100 simulated replicates: medians (points) and interquartile ranges (bars). **d**, Scalar Wald estimates: medians (points), central 95% ranges of the replicate estimates (bars; not per-replicate confidence intervals) and the analytical scalar estimand *θ*_s_ under the non-proportional architecture (blue ticks). Dashed line, *θ* = 0.4, the estimand of every coding under proportionality. Detailed numbers are in Supplementary Note 4.

**Extended Data Fig. 8.**
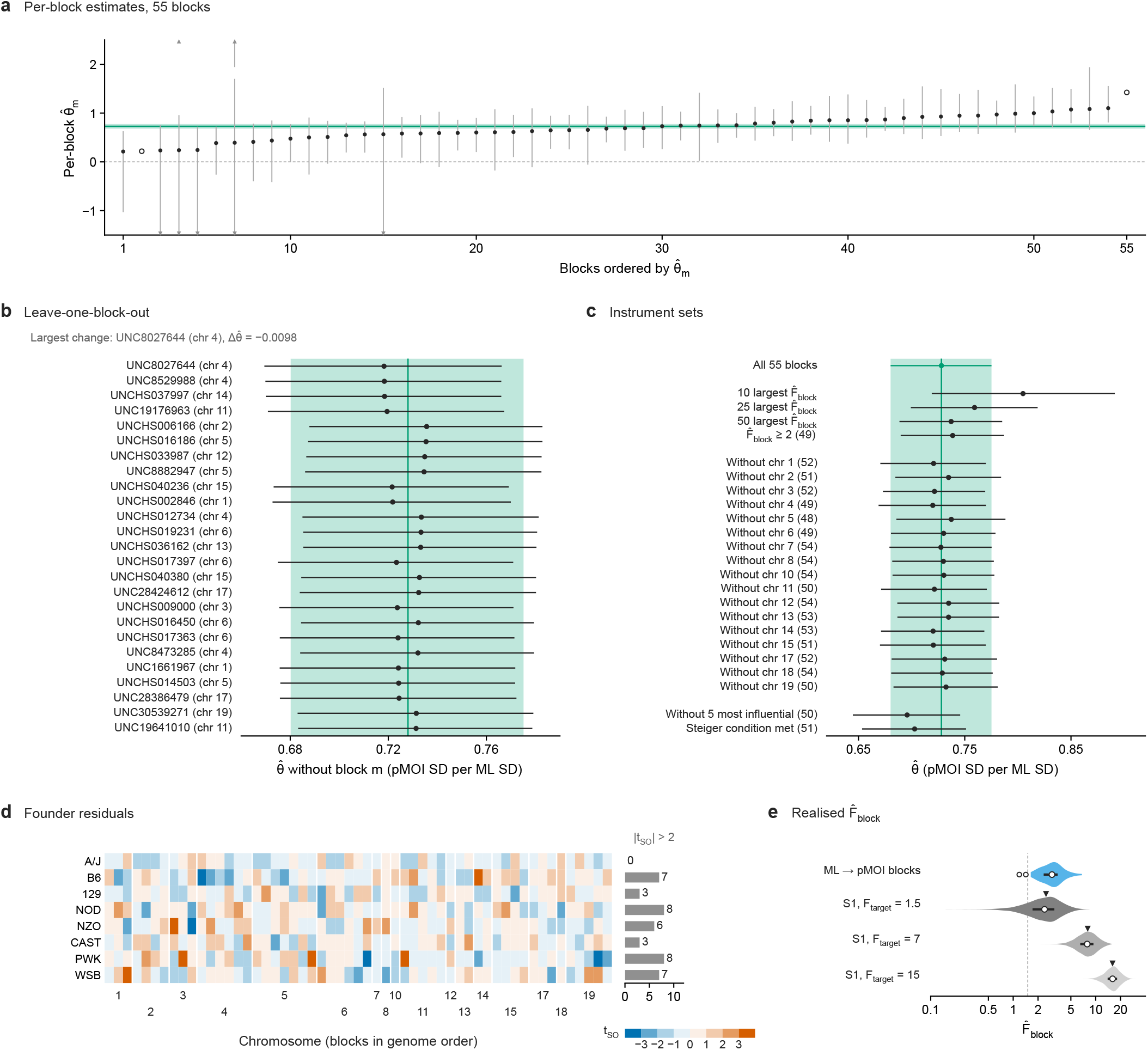
Diagnostics for the ML → pMOI multi-locus analysis. Panels a–d use the 55 founder blocks with 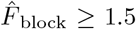 (2 of 57 candidate blocks excluded); each trait uses its own sample (ML, *n* = 779; pMOI, *n* = 799; 779 shared). Estimates are in pMOI SD per ML SD. The green line and tint show the overlap-aware SO estimate over all 55 blocks and its nominal 95% profile-*Q* interval (approximate; Discussion): 0.728 (0.680 to 0.775). Overall estimate and the 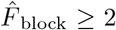 and Steiger sets, Supplementary Table 9. **a**, Per-block SO estimates with per-block nominal 95% profile-*Q* confidence sets, ordered by estimate. For 51 blocks the set is one bounded interval; for 2 it is the union of two unbounded rays, drawn as two segments or axis triangles; for 2 it is the whole real line (open points). Triangles mark a set that continues beyond the axis. **b**, Leave-one-block-out estimates with nominal 95% profile-*Q* intervals for the 25 blocks whose omission moves 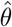 most. The largest change is 0.0098 (UNC8027644, chromosome 4). **c**, Estimates from instrument subsets: the blocks with the 10, 25 and 50 largest 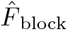; blocks with 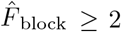; each chromosome left out; the 5 most influential blocks (**b**) left out; and blocks meeting the Steiger condition. Numbers of blocks in parentheses. With only the 10 blocks of largest 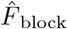, the estimate is 0.805, outside the primary interval (0.680 to 0.775), consistent with the intervals being approximate and possibly too narrow. **d**, One-founder-versus-rest residual scores *t* _SO_ at 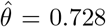 by block (columns, genome order) and founder (rows), in colour steps of 1; |*t* _SO_| *>* 3 takes the darkest step. Bars count blocks with |*t* _SO_ | *>* 2. Founder labels: B6, C57BL/6J; 129, 129S1/SvImJ; NOD, NOD/ShiLtJ; NZO, NZO/HlLtJ; CAST, CAST/EiJ; PWK, PWK/PhJ; WSB, WSB/EiJ. **e**, Realised 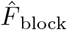 for the 55 analysed blocks (open points, the 2 excluded blocks) and for simulation S1 replicates at *F* _target_ = 1.5, 7 and 15 (3,500 each). Violins, smoothed densities (log scale); bars, interquartile ranges; white points, medians. Triangles mark *F* _target_ + 1, the approximate expected 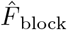; the dashed line marks the inclusion threshold 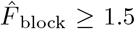.

**Extended Data Fig. 9.**
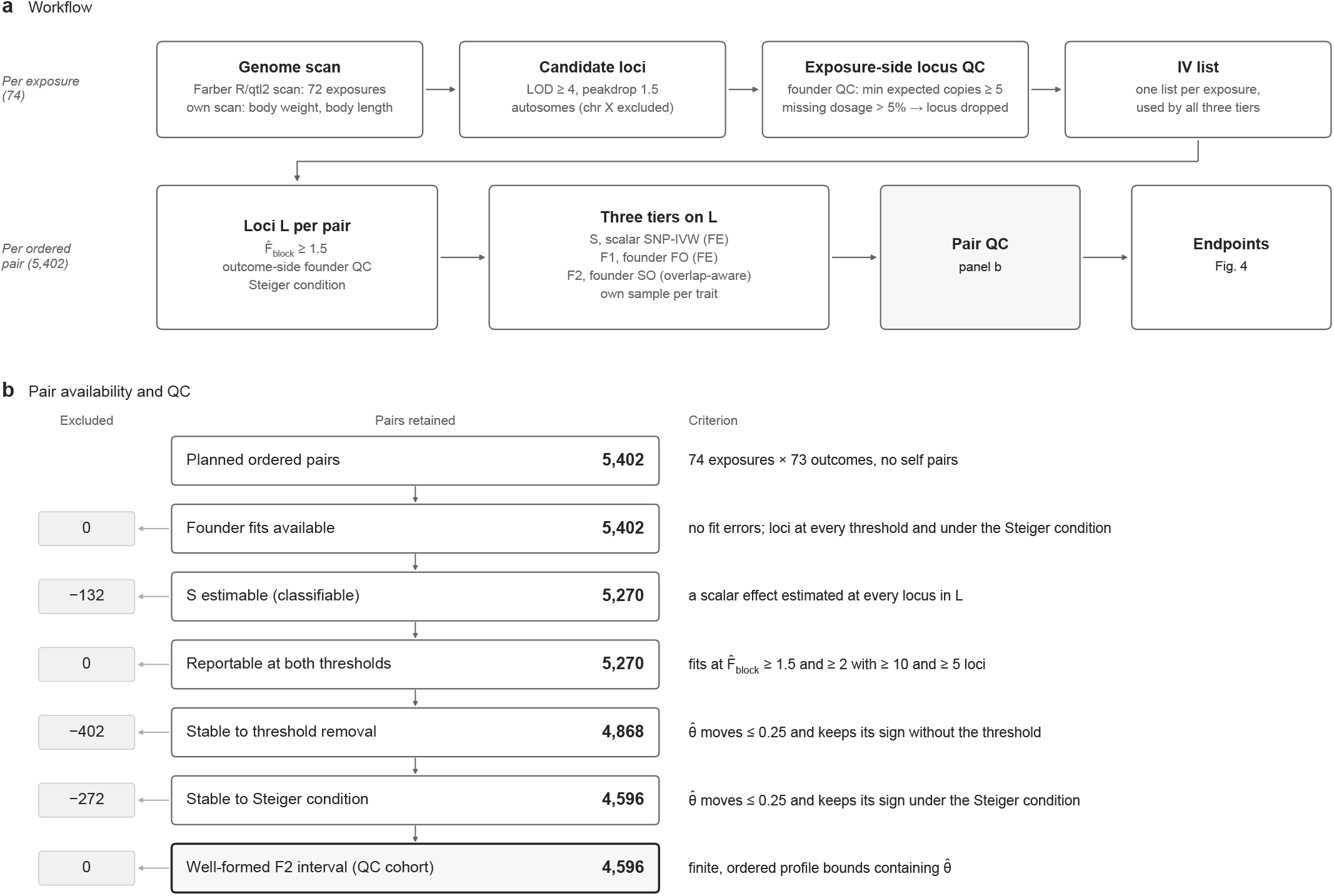
Instrument selection, pair availability and quality control of the trait-pair landscape. **a**, Workflow. Candidate loci for the 72 exposures of the main set are taken from an existing R/qtl2 genome scan (Farber laboratory); body weight and body length use our own scan (Methods). Candidate loci are autosomal peaks with LOD ≥ 4, separated by a LOD drop of at least 1.5 (peakdrop). A locus is dropped when, in the exposure’s sample, the rarest founder has fewer than 5 expected allele copies (founder QC) or more than 5% of the exposure’s sample lacks observed dosage; this gives one instrument list per exposure. For each ordered pair, the loci L are the listed loci with 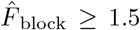 that also pass founder QC in the outcome’s sample and meet the per-locus Steiger condition. The three tiers (S, scalar SNP-IVW (FE); F1, founder FO (FE); F2, founder SO (overlap-aware)) are fitted on L, each trait on its own sample. **b**, Pair availability and QC. Of the 74 *×* 73 = 5,402 planned ordered pairs, all had founder fits with loci, and 132 were excluded because a scalar effect could not be estimated at a locus in L, leaving 5,270 classifiable pairs. QC steps were applied in the order shown, and each pair is counted at the first step it fails: reportable fits under both the inclusion threshold 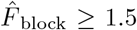 and the alternative 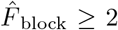, with at least 10 and 5 loci, respectively, counted before the Steiger condition is applied; estimates within 0.25 of, and with the same sign as, the estimate without any threshold; the same stability with and without the Steiger condition; and a finite F2 estimate inside finite, ordered profile bounds. The resulting QC cohort of 4,596 pairs is analysed in Fig. 4. Because QC requires stability across thresholds, the 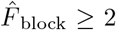 sensitivity analysis (Supplementary Table 3) is not independent of this denominator.

