## Supplementary Information for "Extending Mendelian randomisation to founder-haplotype data in Diversity Outbred mice"

Supplementary Tables 1–10 are supplied in a separate workbook. Supplementary Fig. 1 is in Supplementary Note 6.

- Supplementary Note 1: Founder GLS-IVW.
- Supplementary Note 2: Founder contrasts and the fitter.
- Supplementary Note 3: Cross-covariance under sample overlap.
- Supplementary Note 4: Simulations.
- Supplementary Note 5: Additional methods.
- Supplementary Note 6: Basis-rotation decomposition.

### Supplementary Note 1. Founder GLS-IVW

For retained blocks $m = 1, \ldots, M$, the founder-contrast coefficient estimates $\widehat{b}_{X,m}$ and $\widehat{b}_{Y,m}$ are taken with their covariance matrices $\Sigma_{X,m}$, $\Sigma_{Y,m}$ and $\Sigma_{XY,m} = \mathrm{Cov}(\widehat{b}_{X,m}, \widehat{b}_{Y,m})$, treated as known. The working model is

$$\widehat{b}_{T,m} = b_{T,m} + \xi_{T,m}, \qquad \begin{pmatrix} \xi_{X,m} \\ \xi_{Y,m} \end{pmatrix} \sim N\left\{0,\ C_m = \begin{pmatrix} \Sigma_{X,m} & \Sigma_{XY,m} \\ \Sigma_{XY,m}^{\mathsf{T}} & \Sigma_{Y,m} \end{pmatrix}\right\}, \tag{S1}$$

independently across retained blocks, and $\Gamma_m = \Sigma_{XY,m} + \Sigma_{XY,m}^{\mathsf{T}}$. The block index is dropped where a statement holds block by block.

#### 1.1 The second-order objective

For a candidate effect $\theta$, the founder-contrast residual $d_m(\theta) = \widehat{b}_{Y,m} - \theta\widehat{b}_{X,m}$ has covariance

$$\Omega_m(\theta) = \mathrm{Var}\{d_m(\theta)\} = \Sigma_{Y,m} + \theta^2\Sigma_{X,m} - \theta\Gamma_m \tag{S2}$$

under (S1). This is the exact covariance for every $\theta$, not a linearisation, because $d_m(\theta)$ is linear in the two estimates. The objective is $Q(\theta) = \sum_m d_m(\theta)^{\mathsf{T}}\Omega_m(\theta)^{-1}d_m(\theta)$ (Methods). Under the common-effect model $b_{Y,m} = \theta_0 b_{X,m}$, $d_m(\theta_0) = \xi_{Y,m} - \theta_0\xi_{X,m} \sim N\{0, \Omega_m(\theta_0)\}$, so

$$Q(\theta_0) \sim \chi^2_{7M} \tag{S3}$$

exactly when the covariances are known. $Q(\theta_0)$ is the Wald statistic for the $7M$ restrictions $b_{Y,m} - \theta_0 b_{X,m} = 0$, the founder-block analogue of the Anderson–Rubin statistic for $\theta = \theta_0$ [1].

#### 1.2 Derivatives and local curvature

With $\Omega_m'(\theta) = 2\theta\Sigma_{X,m} - \Gamma_m$, $\Omega_m'' = 2\Sigma_{X,m}$ and $u_m(\theta) = \Omega_m(\theta)^{-1}d_m(\theta)$,

$$Q'(\theta) = -\sum_m \big\{2\,\widehat{b}_{X,m}^{\mathsf{T}}u_m + u_m^{\mathsf{T}}\Omega_m' u_m\big\}, \tag{S4}$$

$$Q''(\theta) = \sum_m \big\{2\,\widehat{b}_{X,m}^{\mathsf{T}}\Omega_m^{-1}\widehat{b}_{X,m} + 4\,\widehat{b}_{X,m}^{\mathsf{T}}\Omega_m^{-1}\Omega_m' u_m + 2\,u_m^{\mathsf{T}}\Omega_m'\Omega_m^{-1}\Omega_m' u_m - 2\,u_m^{\mathsf{T}}\Sigma_{X,m}u_m\big\}. \tag{S5}$$

At the estimate, $Q'(\widehat{\theta}) = 0$ gives $\sum_m \widehat{b}_{X,m}^{\mathsf{T}}u_m = -\frac{1}{2}\sum_m u_m^{\mathsf{T}}\Omega_m' u_m$: the fitted residuals need not be orthogonal to $\widehat{b}_X$ in the $\Omega^{-1}$ metric because the weight depends on $\theta$, so they differ from fixed-weight projection residuals. The local-curvature standard error is

$$\widehat{\mathrm{SE}}_{\mathrm{curv}} = \big\{Q''(\widehat{\theta})/2\big\}^{-1/2}, \tag{S6}$$

the standard error implied by a quadratic approximation of $Q$ at its minimum; when $Q$ is locally quadratic, $\widehat{\theta} \pm 1.96\,\widehat{\mathrm{SE}}_{\mathrm{curv}}$ coincides with the profile-$Q$ interval. Curvature standard errors give the per-locus weights $w_m = \widehat{\mathrm{SE}}_{\mathrm{curv},m}^{-2}$ of Methods.

#### 1.3 Numerical minimisation

The search in qtl2mr covers the whole real line. It writes $\theta = c\tan\alpha$, with a fixed data-dependent scale $c > 0$, so that $\alpha \in [-\pi/2, \pi/2]$ covers $\theta \in \mathbb{R}$ and its limits $\pm\infty$, and evaluates $Q$ on 257 equally spaced values of $\alpha$, supplemented by the first-order estimate $\widehat{\theta}_{\mathrm{FO}}$ (Section 1.5) and $\widehat{\theta}_{\mathrm{FO}} \pm 1$ and $\pm 4$ first-order standard errors. Because $Q$ is a rational function of $\theta$ (Section 1.4), it can have several local minima when the first stage is weak, so every local minimum on this grid is refined, by Brent's method between the neighbouring grid points followed by Newton steps with analytic derivatives. The smallest value found is $Q^{\min}$ (Section 1.4); when it is not distinguishable from the limit $Q_\infty$ as $|\theta| \to \infty$, no finite estimate is reported. A grid search with local refinement does not guarantee the global minimum.

#### 1.4 The profile-$Q$ set

Dividing numerator and denominator by $\theta^2$,

$$Q(\theta) = \sum_m \Big(\frac{\widehat{b}_{Y,m}}{\theta} - \widehat{b}_{X,m}\Big)^{\mathsf{T}} \Big(\frac{\Sigma_{Y,m}}{\theta^2} + \Sigma_{X,m} - \frac{\Gamma_m}{\theta}\Big)^{-1} \Big(\frac{\widehat{b}_{Y,m}}{\theta} - \widehat{b}_{X,m}\Big)$$
$$\longrightarrow\; Q_\infty = \sum_m \widehat{b}_{X,m}^{\mathsf{T}} \Sigma_{X,m}^{-1} \widehat{b}_{X,m} = 7 \sum_m \widehat{F}_{X,m}^{\mathrm{block}} \qquad \text{(S7)}$$

as $\theta \to \pm\infty$. Let $Q^{\min} = \inf_\theta Q(\theta)$, the infimum over the real line, which is the limit $Q_\infty$ when $Q$ approaches its smallest value only as $|\theta| \to \infty$; when the minimum is attained at a finite $\widehat{\theta}$, $Q^{\min} = Q(\widehat{\theta})$. With $c_{0.95} = \chi^2_{1,0.95} = 3.841$, the nominal 95% profile-$Q$ set is $\mathcal{C} = \{\theta : Q(\theta) - Q^{\min} \le c_{0.95}\}$. $\mathcal{C}$ need not be a single bounded interval: it can be unbounded or a union of disjoint intervals.

*Unboundedness.* If $Q_\infty < Q^{\min} + c_{0.95}$, $\mathcal{C}$ contains both sufficiently distant tails of the real line; if $Q_\infty > Q^{\min} + c_{0.95}$, it contains neither. At equality, tail inclusion depends on how $Q(\theta)$ approaches its limit. For a single block the strict sufficient condition is $7\widehat{F}_X^{\mathrm{block}} - Q^{\min} < c_{0.95}$ (Methods). Because $Q^{\min}$ is approximately $\chi^2_6$ when the model holds (Section 1.6), unbounded sets are common when $\widehat{F}_X^{\mathrm{block}}$ is near or below $(c_{0.95}+6)/7 \approx 1.4$. Their frequencies are in Supplementary Note 4 and Supplementary Table 1b.

*Proportional covariances: closed form.* Suppose $\Sigma_{X,m} = s_X \Psi_m$, $\Sigma_{Y,m} = s_Y \Psi_m$ and $\Sigma_{XY,m} = s_{XY}\Psi_m$ with scalars common to all blocks. With $A_{YY} = \sum_m \widehat{b}_{Y,m}^{\mathsf{T}} \Psi_m^{-1} \widehat{b}_{Y,m}$, $A_{XY} = \sum_m \widehat{b}_{X,m}^{\mathsf{T}} \Psi_m^{-1} \widehat{b}_{Y,m}$ and $A_{XX} = \sum_m \widehat{b}_{X,m}^{\mathsf{T}} \Psi_m^{-1} \widehat{b}_{X,m}$,

$$Q(\theta) = \frac{A_{YY} - 2A_{XY}\theta + A_{XX}\theta^2}{s_Y - 2s_{XY}\theta + s_X\theta^2} = \frac{v^{\mathsf{T}} N v}{v^{\mathsf{T}} D v},$$
$$v = \begin{pmatrix} 1 \\ -\theta \end{pmatrix}, \qquad N = \begin{pmatrix} A_{YY} & A_{XY} \\ A_{XY} & A_{XX} \end{pmatrix}, \qquad D = \begin{pmatrix} s_Y & s_{XY} \\ s_{XY} & s_X \end{pmatrix}. \qquad \text{(S8)}$$

As $\theta$ runs over the real line and $\theta \to \pm\infty$ (direction $v \propto (0,1)^{\mathsf{T}}$), $v$ covers every direction in $\mathbb{R}^2$, so $Q^{\min}$ is the smaller root of $\det(N - \lambda D) = 0$. With $k = Q^{\min} + c_{0.95}$, $\mathcal{C} = \{\theta : q_2\theta^2 - 2q_1\theta + q_0 \le 0\}$ with $q_2 = A_{XX} - ks_X$, $q_1 = A_{XY} - ks_{XY}$ and $q_0 = A_{YY} - ks_Y$: a bounded interval if $q_2 > 0$, the union of two rays or the whole line if $q_2 < 0$, a ray if $q_2 = 0$ and $q_1 \ne 0$, and the whole line if $q_2 = q_1 = 0$. These are the shapes of confidence sets for a ratio of regression coefficients, and of Anderson–Rubin sets[1].

*General case.* When $\Sigma_{XY,m} = 0$, write $\Sigma_{Y,m} = R_m^{\mathsf{T}} R_m$ and diagonalise

$$R_m^{-\mathsf{T}} \Sigma_{X,m} R_m^{-1} = V_m \operatorname{diag}(\lambda_{m1}, \ldots, \lambda_{m7}) V_m^{\mathsf{T}}, \qquad \tilde{u}_m = V_m^{\mathsf{T}} R_m^{-\mathsf{T}} \widehat{b}_{Y,m}, \qquad \tilde{v}_m = V_m^{\mathsf{T}} R_m^{-\mathsf{T}} \widehat{b}_{X,m}.$$

Then

$$Q(\theta) = \sum_{m=1}^{M} \sum_{j=1}^{7} \frac{(\tilde{u}_{mj} - \theta \tilde{v}_{mj})^2}{1 + \lambda_{mj}\theta^2}, \tag{S9}$$

a sum of $7M$ rational terms, so $\mathcal{C}$ can have several components.

*Computation.* qtl2mr solves for every crossing of $Q^{\min} + c_{0.95}$ between neighbouring points of its whole-line search (Section 1.3), classifies $\mathcal{C}$ as bounded, disconnected, unbounded, the whole line or unresolved, and reports finite bounds only for a single bounded component.

#### 1.5 First-order weighting and the independence objective

The first-order (FO) estimator treats $\widehat{b}_X$ as fixed:

$$\widehat{\theta}_{\mathrm{FO}} = \frac{\sum_m \widehat{b}_{X,m}^{\mathsf{T}} \Sigma_{Y,m}^{-1} \widehat{b}_{Y,m}}{\sum_m \widehat{b}_{X,m}^{\mathsf{T}} \Sigma_{Y,m}^{-1} \widehat{b}_{X,m}}, \qquad \widehat{\mathrm{SE}}_{\mathrm{FO}} = \Big(\sum_m \widehat{b}_{X,m}^{\mathsf{T}} \Sigma_{Y,m}^{-1} \widehat{b}_{X,m}\Big)^{-1/2}. \tag{S10}$$

It minimises $Q_{\mathrm{FO}}(\theta) = \sum_m d_m(\theta)^{\mathsf{T}} \Sigma_{Y,m}^{-1} d_m(\theta)$, which is quadratic in $\theta$, so its profile set is the first-order Wald interval $\widehat{\theta}_{\mathrm{FO}} \pm c_{0.95}^{1/2} \widehat{\mathrm{SE}}_{\mathrm{FO}}$, the comparator in the simulations (Fig. 3b).

*Estimating equations.* Under (S1) and $b_{Y,m} = \theta_0 b_{X,m}$, $\mathbb{E}\{d_m(\theta_0)\xi_{X,m}^{\mathsf{T}}\} = \Sigma_{XY,m}^{\mathsf{T}} - \theta_0 \Sigma_{X,m}$ and $\mathrm{Var}\{d_m(\theta_0)\} = \Omega_m(\theta_0)$, so at $\theta_0$

$$\begin{aligned} \mathbb{E}\{\widehat{b}_{X,m}^{\mathsf{T}} \Omega_m^{-1} d_m\} &= \mathrm{tr}\{\Omega_m^{-1}(\Sigma_{XY,m}^{\mathsf{T}} - \theta_0 \Sigma_{X,m})\}, \\ \mathbb{E}(u_m^{\mathsf{T}} \Omega_m' u_m) &= \mathrm{tr}(\Omega_m^{-1}\Omega_m') = 2\theta_0\, \mathrm{tr}(\Omega_m^{-1}\Sigma_{X,m}) - 2\,\mathrm{tr}(\Omega_m^{-1}\Sigma_{XY,m}). \end{aligned}$$

Substituting in (S4),

$$\mathbb{E}\{Q'(\theta_0)\} = 0 \tag{S11}$$

for any first-stage strength and any degree of sample overlap, provided the blocks are not selected on the same data (Supplementary Note 4, Section 4.4): the second-order estimating equation is unbiased at the true value. An unbiased estimating equation does not by itself make $\widehat{\theta}$ centred in finite samples.

For the FO objective,

$$\mathbb{E}\{Q_{\mathrm{FO}}'(\theta_0)\} = -2 \sum_m \mathrm{tr}\big\{\Sigma_{Y,m}^{-1}(\Sigma_{XY,m}^{\mathsf{T}} - \theta_0 \Sigma_{X,m})\big\}, \tag{S12}$$

which is generally non-zero. A ratio-of-expectations approximation gives

$$\mathbb{E}(\widehat{\theta}_{\mathrm{FO}}) - \theta_0 \approx \frac{\sum_m \mathrm{tr}\{\Sigma_{Y,m}^{-1}(\Sigma_{XY,m}^{\mathsf{T}} - \theta_0 \Sigma_{X,m})\}}{\sum_m \{b_{X,m}^{\mathsf{T}} \Sigma_{Y,m}^{-1} b_{X,m} + \mathrm{tr}(\Sigma_{Y,m}^{-1}\Sigma_{X,m})\}}. \tag{S13}$$

The $\theta_0\, \mathrm{tr}(\Sigma_Y^{-1}\Sigma_X)$ term is attenuation towards zero from treating the noisy $\widehat{b}_X$ as fixed (for $\Sigma_X = \Sigma_Y$ and $\Sigma_{XY} = 0$, $\mathbb{E}\widehat{\theta}_{\mathrm{FO}} \approx \theta_0 F/(F+1)$ with $F = b_X^{\mathsf{T}} \Sigma_X^{-1} b_X / 7$, the population block statistic, $F_{\mathrm{target}}$ in the simulations); the $\mathrm{tr}(\Sigma_Y^{-1}\Sigma_{XY}^{\mathsf{T}})$ term is a sample-overlap bias towards the slope of the correlated estimation errors. Numerator and denominator both grow linearly in $M$, so neither bias shrinks as blocks are added.

*FO interval width.* Under a strong-first-stage linear approximation,

$$\widehat{\theta}_{\mathrm{FO}} - \theta_0 \approx \frac{\sum_m b_{X,m}^{\mathsf{T}} \Sigma_{Y,m}^{-1} d_m(\theta_0)}{\sum_m b_{X,m}^{\mathsf{T}} \Sigma_{Y,m}^{-1} b_{X,m}},$$

whose variance relative to the nominal FO variance is approximately

$$\frac{\sum_m b_{X,m}^\top \Sigma_{Y,m}^{-1} \Omega_m(\theta_0) \Sigma_{Y,m}^{-1} b_{X,m}}{\sum_m b_{X,m}^\top \Sigma_{Y,m}^{-1} b_{X,m}}.$$

If $\Sigma_X = g\Sigma_Y$ and $\Gamma_m = 2f\Sigma_{Y,m}$, this factor is $1 + g\theta_0^2 - 2f\theta_0$, and, ignoring bias, nominal 95% FO coverage is about $2\Phi\{1.96(1 + g\theta_0^2 - 2f\theta_0)^{-1/2}\} - 1$: 0.95 at $\theta_0 = 0$ and 0.83 at $|\theta_0| = 1$ for $g = 1$ and $f = 0$. Empirical coverage is in Supplementary Note 4, Section 4.1.

*Independence objective.* Setting $\Sigma_{XY,m} = 0$ in $\Omega_m$ when the samples overlap gives $\Omega_m^{\text{ind}}(\theta) = \Sigma_{Y,m} + \theta^2 \Sigma_{X,m}$ and

$$\mathbb{E}\{Q'_{\text{ind}}(\theta_0)\} = -2 \sum_m \Big[ \operatorname{tr}\big\{(\Omega_m^{\text{ind}})^{-1} \Sigma_{XY,m}^\top\big\} - \theta_0^2 \operatorname{tr}\big\{(\Omega_m^{\text{ind}})^{-1} \Sigma_{X,m} (\Omega_m^{\text{ind}})^{-1} \Gamma_m\big\}\Big], \tag{S14}$$

which is non-zero in general when $\Sigma_{XY,m} \neq 0$ (with $\Sigma_X = g\Sigma_Y$ and $\Sigma_{XY} = f\Sigma_Y$ it equals $-14Mf(1 - g\theta_0^2)/(1 + g\theta_0^2)^2$, which vanishes only at $\theta_0^2 = 1/g$). When the overlap structure is shared, every block contributes a term of the same sign, so the bias of the independence estimate is of order one in $M$ while its standard error falls as $M^{-1/2}$: its coverage falls towards zero as blocks are added (S4 sample-overlap arm, Supplementary Table 1f(iii)), whereas for a single block the bias is not compounded. This is the founder-block form of the bias from participant overlap in summary-data MR[2].

#### 1.6 Founder-block lack of fit

*Single locus.* If $b_X$ were known, $\min_\theta (\widehat{b}_Y - \theta b_X)^\top \Sigma_Y^{-1} (\widehat{b}_Y - \theta b_X)$ would be the residual sum of squares of a GLS regression of seven observations on one regressor with known covariance, and would be exactly $\chi_6^2$ under $b_Y = \theta b_X$. For the second-order objective, $Q^{\min} \leq Q(\theta_0) \sim \chi_7^2$ always (S3), and with a strong first stage $Q(\theta_0) - Q^{\min}$ is approximately $\chi_1^2$, so $Q^{\min}$ is approximately $\chi_6^2$[3,4]. Its null calibration is reported in Supplementary Note 4, Sections 4.1 and 4.2.

*Multiple loci.* $Q^{\min}$ combines non-proportionality within blocks and inconsistency between loci, and the $\theta$-dependent metric, the overlap term and the selection of loci make the fixed-metric $\chi_{7M-1}^2$ reference approximate. Multi-locus $Q^{\min}$ $P$ values referred to it (Methods) are therefore not formal tests.

#### 1.7 Scalar estimand under binary recoding (simulation S3)

For a binary founder coding $s \in \{0,1\}^8$, the scalar genotype is $g_s = P_m s$. With $M_T$ the residual-making matrix for trait $T$ (Methods), the scalar coefficient, its variance and the scalar first-stage statistic of the one-dimensional qtl2-compatible mixed model are

$$\widehat{\beta}_{g_sT} = (g_s^\top M_T g_s)^{-1} g_s^\top M_T T, \qquad \operatorname{Var}(\widehat{\beta}_{g_sT}) = \widehat{\sigma}_T^2 (g_s^\top M_T g_s)^{-1}, \qquad F_s = \widehat{\beta}_{g_sX}^2 / \operatorname{Var}(\widehat{\beta}_{g_sX}), \tag{S15}$$

and the population projection of the founder signal $Z_m b_T$ on $g_s$ is $\beta^\star_{g_sT} = (g_s^\top M_T g_s)^{-1} g_s^\top M_T Z_m b_T$. When both traits are fitted with the same working covariance ($M_X = M_Y = M$, as in S3) and $b_Y = \theta b_X + \delta$, linearity gives

$$\theta_s = \frac{\beta^\star_{g_sY}}{\beta^\star_{g_sX}} = \theta + \frac{\beta^\star_{g_s\delta}}{\beta^\star_{g_sX}}, \tag{S16}$$

where $\beta^\star_{g_s\delta}$ is the projection of the non-proportional component $Z_m\delta$ on $g_s$. Under proportionality ($\delta = 0$) every coding with a non-zero first stage has $\theta_s = \theta$; otherwise different codings of the same founder architecture target different scalar estimands. With different working covariances for the two traits, $\theta_s$ also depends on $M_X$ and $M_Y$ under proportionality.

### Supplementary Note 2. Founder contrasts and the fitter

#### 2.1 Founder contrasts and invariance to the contrast basis

Because each row of the founder-probability matrix $P$ sums to the same constant $c$, $P\mathbf{1}_8 = c\mathbf{1}_n$, write the orthogonal $8 \times 8$ matrix $[\mu, H]$ with $\mu = \mathbf{1}_8/\sqrt{8}$ and $H^\top \mathbf{1}_8 = 0$, $H^\top H = I_7$. For any founder-effect vector $a \in \mathbb{R}^8$, $Pa = P(\mu\mu^\top + HH^\top)a = (c/\sqrt{8})(\mu^\top a)\mathbf{1}_n + Z(H^\top a)$ with $Z = PH$: the first term is absorbed by the intercept, and only $b = H^\top a$ is identifiable. For sum-to-zero founder effects, $a = Hb$. Founders are coded A–H in the standard Diversity Outbred order (A/J, C57BL/6J, 129S1/SvImJ, NOD/ShiLtJ, NZO/HlLtJ, CAST/EiJ, PWK/PhJ, WSB/EiJ). The software builds $H$ from the column-normalised Helmert contrasts of R (`stats::contr.helmert`): for $k = 1, \ldots, 7$, $H_{ik} = -\{k(k+1)\}^{-1/2}$ for $i \le k$, $H_{k+1,k} = k\{k(k+1)\}^{-1/2}$ and $H_{ik} = 0$ for $i > k+1$.

Let $H^\star = HR$ for an orthogonal $7 \times 7$ matrix $R$, applied to every exposure and outcome. Then $Z^\star = ZR$, and the fitted model gives $\widehat{b}^\star_T = R^\top \widehat{b}_T$, $A^\star_T = R^\top A_T$ for both the original-space and the star-space influence operators, and $\Sigma^\star_T = R^\top \Sigma_T R$. Every cross-covariance of the form $A_X(\cdot)A_Y^\top$, including the star-space $\widehat{\Sigma}_{XY}$ of Supplementary Note 3, becomes $R^\top \Sigma_{XY} R$. Hence $d^\star_m(\theta) = R^\top d_m(\theta)$ and $\Omega^\star_m(\theta) = R^\top \Omega_m(\theta) R$, so

$$Q^\star(\theta) = d_m(\theta)^\top R\,\{R^\top \Omega_m(\theta) R\}^{-1} R^\top d_m(\theta) = Q(\theta) \quad \text{for every } \theta, \tag{S17}$$

and $\widehat{\theta}$, the profile-$Q$ set and $Q^{\min}$ are unchanged, as are $\widehat{F}^{\text{block}}_X = \widehat{b}_X^\top \Sigma_X^{-1} \widehat{b}_X/7$, the founder-space residual $\widehat{e} = H^\star \widehat{d}^\star = HRR^\top \widehat{d} = H\widehat{d}$ and the founder residual scores $t^{\text{SO}}_f$ (because $P^\star_\perp = R^\top P_\perp R$, the residual covariance in founder coordinates is unchanged). The per-locus estimates, two-step IVW, weighted median, weighted mode, $Q_{\text{between}}$ and leave-one-out summaries are functions of these quantities and are also invariant (numerical check, Supplementary Table 5). $R = -I_7$ shows that the sign convention of $H$ is immaterial. More generally, for any invertible $7 \times 7$ matrix $T$ and $H^\star = HT$ (for example a non-orthonormal or a centred treatment-coded basis), $\widehat{b}^\star_T = T^{-1}\widehat{b}_T$, $\Sigma^\star_T = T^{-1}\Sigma_T T^{-\top}$ and $\widehat{\Sigma}^\star_{XY} = T^{-1}\widehat{\Sigma}_{XY} T^{-\top}$, while the residuals and $\widehat{\rho}_*$ are unchanged; hence $d^{\star\top}_m \Omega^{\star-1}_m d^\star_m = d_m^\top \Omega_m^{-1} d_m$, and $\widehat{\theta}$, the profile-$Q$ set and $Q^{\min}$ are unchanged. The fitter accepts only orthonormal $H$; quantities defined through $H$ itself are not invariant.

#### 2.2 The full-covariance fitter

For each trait $T$ and block, the fitter reproduces the R/qtl2 founder-contrast linear mixed model (Methods, equation 1). The residual heritability $h^2_T$ is estimated once, by qtl2mr's own profile restricted maximum likelihood under the null model with the same samples, covariates and kinship matrix as the coefficient fit, over $h^2_T \in [0, 1]$, and then held fixed; the residual covariance is proportional to $h^2_T(2K) + (1 - h^2_T)I$, and eigenvalues of $2K$ that are negative within rounding are set to zero. The multi-locus analyses use one value per trait and chromosome because the leave-one-chromosome-out (LOCO) kinship differs by chromosome (Supplementary Table 2). With the eigen-decomposition $2K_T = U\Lambda U^\top$ (as in `qtl2::decomp_kinship`) and weights $w_i = (h^2_T \lambda_i + 1 - h^2_T)^{-1/2}$, the rotated data are

$$\widetilde{T} = WU^\top T, \qquad \widetilde{X} = WU^\top [Z, C_T], \qquad W = \operatorname{diag}(w_1, \ldots, w_n), \tag{S18}$$

the transformation used by the R/qtl2 mixed-model scan, under which the residuals are independent with variance $\sigma^2_T$. A column-pivoted QR decomposition of $\widetilde{X}$ gives the numerical rank, the coefficient estimates, the residual degrees of freedom $\nu = n - \operatorname{rank}(\widetilde{X})$, $\widehat{\sigma}^2_T = \|\widetilde{T} - \widetilde{X}\widehat{\beta}\|^2/\nu$ and the full coefficient covariance $\widehat{\sigma}^2_T(\widetilde{X}^\top \widetilde{X})^{-1}$, whose $7 \times 7$ founder-contrast block is $\Sigma_T$ of Methods equation 2 (by the Frisch–Waugh–Lovell theorem, $\Sigma_T = \widehat{\sigma}_T^2(Z^\mathsf{T} M_T Z)^{-1}$). Writing $\widetilde{A}_T$ for the founder-contrast rows of the least-squares map from $\widetilde{T}$ to the coefficients, the influence operators are

$$A_T = \widetilde{A}_T W U^\mathsf{T} \quad (\widehat{b}_T = A_T T), \qquad A_T^* = \widetilde{A}_T U^\mathsf{T} \quad (\widehat{b}_T = A_T^* T^*), \qquad S_T = UWU^\mathsf{T}, \tag{S19}$$

where $S_T$ is the symmetric whitening matrix of Methods ($S_T^\mathsf{T} S_T = V_T^{-1}$) and $T^* = S_T T$. Because $A_T V_T A_T^\mathsf{T} = (Z^\mathsf{T} M_T Z)^{-1}$ (using $M_T V_T M_T = M_T$), $\Sigma_T = \widehat{\sigma}_T^2 A_T^* A_T^{*\mathsf{T}}$, a fact used in Supplementary Note 3.

Standard `qtl2::scan1coef` with `se=TRUE` returns $\widehat{b}_T$ and the marginal standard errors $D_T = \mathrm{diag}\{\mathrm{SE}_{T,1}, \ldots, \mathrm{SE}_{T,7}\}$. With $R_T$ the correlation matrix of $(Z^\mathsf{T} M_T Z)^{-1}$,

$$\Sigma_T = D_T R_T D_T \tag{S20}$$

is an identity when the samples, covariates, kinship matrix, $h_T^2$, contrast order and rank handling agree, and provides an algebraic reconstruction of the full block from the standard output (Supplementary Table 4).

### Supplementary Note 3. Cross-covariance under sample overlap

#### 3.1 Construction

Each trait is fitted separately on its own analysis sample (Supplementary Note 2), with its own heritability: the exposure with $h_X^2$ and whitening matrix $S_X$, the outcome with $h_Y^2$ and $S_Y$. Let $\widehat{r}_T^* = S_T\widehat{r}_T$ be the star-space residuals, $O = O_{XY}$ the $n_X \times n_Y$ overlap-matching matrix, and $O_X = OO^\top$ and $O_Y = O^\top O$ the diagonal 0/1 matrices marking the overlapping mice in each sample. After centring each residual vector by its mean over the overlapping mice (giving $\widetilde{r}_X^*$, $\widetilde{r}_Y^*$),

$$\widehat{\rho}_* = \frac{(\widetilde{r}_X^*)^\top O\,\widetilde{r}_Y^*}{\{(\widetilde{r}_X^*)^\top O_X\widetilde{r}_X^*\}^{1/2}\{(\widetilde{r}_Y^*)^\top O_Y\widetilde{r}_Y^*\}^{1/2}}, \qquad \widehat{\Sigma}_{XY} = \widehat{\rho}_*\,\widehat{\sigma}_X\widehat{\sigma}_Y\,A_X^* O A_Y^{*\top}, \tag{S21}$$

that is, $\widehat{\rho}_*$ is the Pearson correlation of the centred star-space residuals over the overlapping mice (Methods, equation 3). The implied residual cross-covariance in the original sample space is $\widehat{\rho}_*\widehat{\sigma}_X\widehat{\sigma}_Y S_X^{-1}OS_Y^{-1}$. $\widehat{\rho}_*$ carries the residual correlation of the shared mice into $\widehat{\Sigma}_{XY}$.

#### 3.2 Same, different or partly overlapping samples

The method is designed for two-sample MR: the exposure and the outcome may be measured on the same mice, on different mice or on partly overlapping sets of mice. The cross-covariance $\widehat{\Sigma}_{XY}$ in (S21) is built from the shared mice only, and is zero when there are none ($O = 0$); the second-order objective then reduces to the independent-samples form, with $\Omega_m(\theta) = \Sigma_{Y,m} + \theta^2\Sigma_{X,m}$ (Supplementary Note 1). The data analysed in this paper come from one cohort, and each trait is analysed on its own complete sample (Methods), so the exposure and outcome samples overlap but need not coincide.

#### 3.3 Exactness of the working model

Let $\varepsilon_X$ and $\varepsilon_Y$ be the residual vectors (polygenic plus environmental) of the two traits, with $\mathrm{Cov}(\varepsilon_X, \varepsilon_Y) = \Delta_{XY}$ ($n_X \times n_Y$). Because $\widehat{b}_T = A_T T = A_T^* S_T T$, the exact coefficient cross-covariance is

$$\mathrm{Cov}(\widehat{b}_X, \widehat{b}_Y) = A_X\Delta_{XY}A_Y^\top = A_X^*\big(S_X\Delta_{XY}S_Y\big)A_Y^{*\top}. \tag{S22}$$

Comparing with (S21), the working model replaces the $n_X \times n_Y$ matrix $S_X\Delta_{XY}S_Y$ by $\rho_*\sigma_X\sigma_Y O$: star-space residuals of different mice are taken to be uncorrelated, and those of the same mouse to share a single correlation $\rho_*$.

*Complete overlap.* Suppose both traits are measured on the same $n$ mice ($O = I_n$) and follow the usual bivariate polygenic model with the same kinship matrix,

$$\Delta_{XY} = \sigma_X\sigma_Y\big[\rho_g h_X h_Y(2K) + \rho_e\{(1-h_X^2)(1-h_Y^2)\}^{1/2}I_n\big], \tag{S23}$$

with genetic correlation $\rho_g$ and environmental correlation $\rho_e$, and suppose that $S_X$ and $S_Y$ are built from the true heritabilities and the same kinship matrix (in the software $h_T^2$ is the REML estimate under the null model, held fixed, so this holds only approximately). With $2K = U\Lambda U^\top$, $V_T = U\{h_T^2\Lambda + (1-h_T^2)I\}U^\top$ and $S_T = V_T^{-1/2}$, all three matrices share the eigenvectors $U$, so

$$\begin{aligned} S_X\Delta_{XY}S_Y &= \sigma_X\sigma_Y\,U\,\mathrm{diag}\{r(\lambda_1),\ldots,r(\lambda_n)\}U^\top, \\ r(\lambda) &= \frac{\rho_g h_X h_Y\lambda + \rho_e\{(1-h_X^2)(1-h_Y^2)\}^{1/2}}{\{(h_X^2\lambda + 1 - h_X^2)(h_Y^2\lambda + 1 - h_Y^2)\}^{1/2}}. \end{aligned} \tag{S24}$$

Star-space residuals are therefore uncorrelated across the eigen-directions of the kinship matrix, but their cross-correlation varies along its spectrum: $r(0) = \rho_e$ in directions without relatedness, $r(1) = \rho_g h_X h_Y + \rho_e\{(1-h_X^2)(1-h_Y^2)\}^{1/2}$ (the residual correlation of the two traits within a mouse when the diagonal of $2K$ is 1), and $r(\lambda) \to \rho_g$ as $\lambda \to \infty$ when $h_X, h_Y > 0$. The working model is exact when $r(\lambda)$ is constant, which holds in particular when

(a) $h_X^2 = h_Y^2$ and $\rho_g = \rho_e = \rho$: then $r(\lambda) \equiv \rho = \rho_*$ (the special case in Methods);

(b) $h_X^2 = h_Y^2 = 0$ (no kinship-structured residual): $S_T = I$ and $\Delta_{XY} = \rho_e \sigma_X \sigma_Y O$, exact for any overlap pattern;

(c) $\rho_g = \rho_e = 0$: $\Delta_{XY} = 0$ and $\widehat{\Sigma}_{XY}$ estimates zero.

When $0 < h_X^2, h_Y^2 < 1$, $h_X^2 \neq h_Y^2$ and $\rho_g = \rho_e = \rho \neq 0$, $r(\lambda)$ is not constant: by the Cauchy–Schwarz inequality $|r(\lambda)| \leq |\rho|$, with equality at finite $\lambda$ only when $\lambda = 0$, so the effective correlation is shrunk in structured directions by an amount that depends on how different the two heritabilities are. For the ML–pMOI analysis, the factor $\min_\lambda r(\lambda)/\rho$ computed from its heritabilities under complete overlap is in Supplementary Table 2(b). When $\rho_g \neq \rho_e$, $r(\lambda)$ varies even when the heritabilities are equal.

*Partial overlap.* When the two samples differ, $S_X$ and $S_Y$ are computed on different sets of mice and need not share eigenvectors, so (S24) does not generally apply. Cases (b) and (c) remain exact for any overlap pattern; case (a) does not generally extend to partial overlap.

##### 3.4 Validity of the joint covariance

Because $\Sigma_T = \widehat{\sigma}_T^2 A_T^* A_T^{*\mathsf{T}}$ (Supplementary Note 2), the joint covariance implied by (S21) is

$$C = \begin{pmatrix} A_X^* & 0 \\ 0 & A_Y^* \end{pmatrix} \begin{pmatrix} \widehat{\sigma}_X^2 I_{n_X} & \widehat{\rho}_* \widehat{\sigma}_X \widehat{\sigma}_Y O \\ \widehat{\rho}_* \widehat{\sigma}_X \widehat{\sigma}_Y O^{\mathsf{T}} & \widehat{\sigma}_Y^2 I_{n_Y} \end{pmatrix} \begin{pmatrix} A_X^* & 0 \\ 0 & A_Y^* \end{pmatrix}^{\mathsf{T}}. \tag{S25}$$

$O$ has at most one non-zero entry, equal to 1, in each row and column, so its singular values are 0 or 1, and the middle matrix is positive semidefinite whenever $|\widehat{\rho}_*| \leq 1$, which holds for a sample correlation. Hence $C \succeq 0$ by construction. Moreover, $P_T = \widehat{\sigma}_T \Sigma_T^{-1/2} A_T^*$ has orthonormal rows, and $\Sigma_X^{-1/2} \widehat{\Sigma}_{XY} \Sigma_Y^{-1/2} = \widehat{\rho}_* P_X O P_Y^{\mathsf{T}}$, so

$$s_{\max}\big(\Sigma_X^{-1/2} \widehat{\Sigma}_{XY} \Sigma_Y^{-1/2}\big) \leq |\widehat{\rho}_*|. \tag{S26}$$

If the two traits share their sample, covariates and heritability, $A_X^* = A_Y^*$ and the bound is attained; when the samples overlap only partly, $A_X^*$ and $A_Y^*$ are computed on different sets of mice and it need not be.

The unwhitened construction $\widehat{\Sigma}_{XY}^{\text{raw}} = \widehat{\rho}_{\text{raw}} \widehat{\sigma}_X \widehat{\sigma}_Y A_X O A_Y^{\mathsf{T}}$, with $\widehat{\rho}_{\text{raw}}$ the correlation of the unwhitened residuals, treats them as independent across mice, ignores the kinship structure of the marginal covariances and need not give a valid joint covariance; it is computed only as a diagnostic. Before use, the software computes $s_{\max}$ and the minimum eigenvalue of $C$; an excess of $s_{\max}$ over 1 at the level of rounding error is clipped, and a larger one stops the fit. By (S26) any excess is rounding error. Real-data values, including the raw-space construction, are in Supplementary Table 7.

### Supplementary Note 4. Simulations

The code of the simulation engine will be included in the R package qtl2founderSim, to support reproduction and further development.

Methods summarises the simulation designs; they are given in full below and in Supplementary Table 1a. Simulations S1–S4 were generated and analysed with the simulation engine; S5 was analysed with qtl2mr 0.7.0.

*Single-locus data-generating process (S1–S3).* The genotype structure of real mice from the *Qsox1* analysis was kept, and only founder effects and residuals were simulated. S1 and S2 used $n = 400$ mice with the founder-probability matrix $P_m$ at the lead marker JAX00272363, the kinship matrix $K$ and the covariates $C$; S3 used a separate design of $n = 500$ mice at the chromosome 1 marker UNC494697 from the same published cross. In each design $Z_m = P_m H$, and the data-generating process was

$$X = C\alpha_X + Z_m b_X + e_X, \qquad Y = C\alpha_Y + Z_m(\theta b_X + \delta) + e_Y, \qquad e_T \sim N\big(0, \sigma_T^2 V_T\big),$$

with $V_T = h_T^2(2K) + (1 - h_T^2)I$, $\sigma_T^2 = 1$ and $h_T^2 = 0.20$ unless stated. Calibration settings used $\delta = 0$. The exposure architecture was set by a raw direction $d_X \in \mathbb{R}^7$, scaled to a target first-stage strength:

$$b_X = s\, d_X, \qquad s = \left\{ \frac{7F_{\text{target}}\sigma_X^2}{d_X^{\mathsf{T}} G_X d_X} \right\}^{1/2}, \qquad G_X = Z_m^{\mathsf{T}} M_X Z_m,$$

with $M_X$ the residual-making matrix of Methods, so that the population founder-block statistic $F_{\text{target}} = \frac{1}{7} b_X^{\mathsf{T}} \Sigma_X^{-1} b_X$, with $\Sigma_X = \sigma_X^2 G_X^{-1}$, is a design quantity, not the fitted statistic $\widehat{F}_X^{\text{block}}$. With known covariances, $7\widehat{F}_X^{\text{block}}$ is non-central $\chi_7^2$ with non-centrality $7F_{\text{target}}$, so $\mathbb{E}\widehat{F}_X^{\text{block}} = F_{\text{target}} + 1$, about 2.5 at $F_{\text{target}} = 1.5$. Sample overlap was introduced by drawing exposure and outcome residuals jointly on the overlapping mice with residual correlation $\rho_{XY}$, giving an effective coefficient-error correlation $\rho_{\text{eff}} = \text{overlap fraction} \times \rho_{XY}$; otherwise residuals were independent. $(P_m, K, C, H)$, the marker, $F_{\text{target}}$ and the architecture were fixed across replicates. Each replicate was refitted with the qtl2-compatible model, with $h^2$ fixed at its design value, to obtain $\widehat{b}_X$, $\widehat{b}_Y$, $\Sigma_X$ and $\Sigma_Y$, then analysed with Founder GLS-IVW. In S2, $\Sigma_{XY}$ combined the design residual correlation with the fitted residual variances; $\rho_*$ was estimated from star-space residuals only in the S1 overlap check.

*Selection.* At $F_{\text{target}} = 1.5$, 81–84% of S1 replicates met the inclusion threshold $\widehat{F}_X^{\text{block}} \geq 1.5$. Unselected results use all replicates or blocks; selected single-locus summaries use only passing replicates, and selected multi-locus fits use only passing blocks. At $F_{\text{target}} \geq 7$ the threshold removed essentially no replicate or block, so the selected and unselected analyses agree (Supplementary Tables 1c–1g); at $F_{\text{target}} = 30$ it removed none, and no profile set was unbounded.

*Coverage and performance measures.* Coverage is membership of the profile-$Q$ set $\mathcal{C}$ of Supplementary Note 1, Section 1.4 (termed "exact" in the tables): a set that is unbounded or a union of disjoint intervals covers $\theta$ when it contains it, and the frequency of unbounded sets is reported (Supplementary Table 1b). $Q^{\min}$ is taken from the closed form (S8) where it applies (the S1 and S2 refits, the S4 sample-overlap arm and the S4 full-covariance comparator) and from a whole-line numerical search otherwise, so coverage does not depend on locating every component of $\mathcal{C}$. First-order coverage uses the Wald interval. Relative error, $e_r = (\widehat{\theta}_r - \theta)/\theta$, is not summarised for null settings; the other measures are listed in Methods.

#### 4.1 Single-locus calibration and sample overlap (S1)

*Design.* Under $b_Y = \theta b_X$ and independent samples, $\theta$ took the values −1, −0.5, −0.25, 0, 0.25, 0.5 and 1 and $F_{\text{target}} \in \{1.5, 7, 15, 30\}$, with 500 replicates per cell; SO profile-$Q$ intervals were compared with first-order intervals that treat $\widehat{b}_X$ as fixed. An overlap calibration, using a synthetic design of 400 exposure and 400 outcome mice with simulated kinship and founder probabilities, at $\theta = 0.25$ and $F_{\text{target}} \in \{1.5, 7, 15, 30\}$ (1,000 replicates per setting) compared the independence objective ($\Sigma_{XY} = 0$) with the overlap-aware objective across overlap fractions and residual correlations. A targeted check covered $h^2 \in \{0.20, 0.50, 0.80\}$, $F_{\text{target}} \in \{15, 30, 100\}$ and $\theta \in \{0.25, 1\}$, with 300 replicates per setting.

With exact proportionality and independent samples, the second-order (SO) estimate was centred on $\theta$ at $F_{\text{target}}$ of 7 and 15 (largest absolute median relative error 2.5%; Extended Data Fig. 3a), with coverage 0.936–0.960 across the seven values of $\theta$ (Fig. 3b) and 0.926–0.960 at $F_{\text{target}} = 30$ (Extended Data Fig. 3b; Supplementary Table 1c(i)). First-order intervals were near nominal coverage at $\theta = 0$ but under-covered increasingly as $|\theta|$ grew, to 0.780–0.790 at $|\theta| = 1$ (Supplementary Note 1, Section 1.5).

At $F_{\text{target}} = 1.5$, unselected SO coverage fell to 0.852–0.916 and 12.9% of sets were unbounded. Same-sample selection raised coverage to 0.877–0.942 and reduced unbounded sets to 4.2%, but moved median estimates towards zero (at $\theta = 1$, from 1.024 to 0.929; Supplementary Table 1c(i)). In the targeted check at $h^2$ up to 0.8 and $F_{\text{target}} \geq 15$, coverage of the interval found by stepping outward from $\widehat{\theta}$ to the first threshold crossing on each side was 0.910–0.977 and $Q^{\min}$ rejection 0.027–0.080 (Supplementary Table 1c(iii)).

In the single-locus overlap check, ignoring exposure–outcome cross-covariance shifted estimates upwards as the effective coefficient-error correlation grew, consistent with overlap bias in conventional MR[2]. Across 18 overlap settings at $F_{\text{target}} = 7$, mean relative error ranged from 0.021 to 0.354 for the independence objective, against $-0.064$ to 0.037 for the overlap-aware objective; the same pattern held at $F_{\text{target}} = 15$. The bias was small relative to interval width: coverage at these strengths remained 0.925–0.958 (overlap-aware) and 0.929–0.981 (independence; Supplementary Table 1c(ii); Extended Data Fig. 3c,d).

#### 4.2 Non-proportional founder contamination (S2)

*Design.* S2 added a component $\delta$ to the proportional outcome effect, with magnitude $\lambda$ relative to $\theta b_X$ and alignment $\kappa$ with the exposure founder pattern. With the metric $W_\theta = \Omega(\theta)^{-1}$, the norm $\|v\|_{W_\theta} = (v^\top W_\theta v)^{1/2}$ and the unit exposure fingerprint $q_X = b_X / \|b_X\|_{W_\theta}$,

$$\delta = \lambda \, \|\theta b_X\|_{W_\theta} \big\{ \kappa q_X + (1 - \kappa^2)^{1/2} q_\perp \big\},$$

where $q_\perp$ is a $W_\theta$-orthogonal unit vector obtained by removing the $W_\theta$-projection on $b_X$ from $u \sim N(0, I_7)$ ($\kappa = 0$ orthogonal, $\kappa = 1$ fully aligned). The projection slope $\theta^\star = \theta + b_X^\top W_\theta \delta / (b_X^\top W_\theta b_X)$ adds to $\theta$ the part of $\delta$ aligned with $b_X$ in this metric. It equals $\theta$ under orthogonal contamination and is the common slope of the contaminated pattern only under full alignment; it is a fixed-metric reference, not in general the estimand of Founder GLS-IVW. The grid was $F_{\text{target}} \in \{1.5, 7, 15, 30\}$, $\theta \in \{0.25, 1\}$, $\lambda \in \{0, 0.25, 0.5, 1\}$ and $\kappa \in \{0, 0.5, 1\}$, with $\rho_{XY} = 0.20$ and 1,000 replicates per setting. $Q^{\min}$ rejection is at $P < 0.05$ under the $\chi^2_6$ reference.

Without contamination ($\lambda = 0$), $Q^{\min}$ rejection at the nominal 0.05 level was 0.042–0.054 at $F_{\text{target}} \geq 7$ and 0.022–0.037 at 1.5 (Supplementary Table 1d(ii)). Across all S2 contamination settings at the weak strength, 14.6% of sets were unbounded without selection and 6.0% with selection (Supplementary Table 1b).

A fully aligned component ($\kappa = 1$) is indistinguishable from a change of slope. At $F_{\text{target}} = 15$, $\theta = 0.25$ and $\lambda = 1$, the median estimate was 0.499, close to the projection slope $\theta^\star = 0.500$. Coverage of $\theta$ fell to 0.28, but coverage of $\theta^\star$ was 0.94 and $Q^{\min}$ rejection remained near nominal (0.046; Supplementary Table 1d(i); Fig. 3c,d).

An orthogonal component ($\kappa = 0$) raised $Q^{\min}$ rejection with $\lambda$: at $\lambda = 1$ and $\theta = 0.25$, rejection reached 0.48 at $F_{\text{target}} = 15$ and 0.20 at $F_{\text{target}} = 7$, while coverage of $\theta$ was 0.91 at the higher strength. A large orthogonal component also shifted the SO estimate: with $\theta = 1$, $\lambda = 1$ and $F_{\text{target}} = 15$, the median estimate was 1.69 although $\theta^\star = 1.00$; coverage fell to 0.05 and $Q^{\min}$ rejected in every replicate (rate 1.00; Supplementary Table 1d(i); Extended Data Fig. 6). The weight depends on $\theta$: a larger slope increases $\theta^2\Sigma_X$ in $\Omega(\theta)$ and down-weights the orthogonal residual. Orthogonality in the metric at the generating $\theta$ therefore does not make contamination neutral for the estimate. Only the component departing from the exposure pattern is detectable.

#### 4.3 Scalar coding (S3)

*Design.* With $h^2 = 0.20$, $F_{\text{target}} = 80$ and $\theta = 0.4$, the outcome followed a non-proportional architecture with $\delta$ orthogonal to $b_X$ and $\|\delta\| = 0.35\|b_X\|$. In each of 100 replicates, the same simulated phenotypes were collapsed under four pre-specified binary codings $s$ (aligned, cancelling, partial and mismatched, i.e. aligned with $\delta$), $g_s = P_m s$, and analysed by one-dimensional qtl2-compatible mixed models, giving scalar first-stage $F$ statistics and Wald ratios. Only the non-proportional architecture was simulated. Under proportionality ($\delta = 0$) every coding with a non-zero first stage has scalar estimand $\theta$ analytically, and this is the proportional reference; under non-proportionality the estimand depends on the coding, $\theta_s = \theta + \beta^\star_{g_s\delta}/\beta^\star_{g_sX}$, where $\beta^\star$ denotes the population projection on $g_s$ (Supplementary Note 1, Section 1.7).

The aligned grouping gave median scalar $F = 213.8$ and Wald ratio 0.386 (analytical estimand $\theta_s = 0.386$). The mismatched grouping, aligned with $\delta$, gave $F = 6.5$ and ratio $-1.309$ ($\theta_s =$-1.347). The cancelling and partial groupings also changed both strength and the scalar estimand (Supplementary Table 1e(ii); Extended Data Fig. 7). The coding dependence under non-proportionality is consistent with evidence that treating a multi-allelic DO QTL as biallelic can mislead mediation analysis[5].

#### 4.4 Multi-locus stacking (S4)

*Design.* Multi-locus experiments used synthetic designs in three arms. (i) Independent pruned blocks: entries $A_{ij} \sim N(0, 1)$ were column-centred and QR-decomposed, and partitioning $Q$ into seven-column blocks gave $Z_m = \sqrt{n}\,Q_m$ with $n = 900$, so that $Z_m^\top Z_m = nI_7$ and $Z_m^\top Z_{m'} = 0$ for $m \neq m'$; directions were normalised and scaled to $F_{\text{target}}$, with $M \in \{20, 100\}$, $F_{\text{target}} \in \{1.5, 7, 15, 30\}$ and $\theta \in \{\pm0.25, \pm0.5, \pm1\}$ (100 replicates per setting). (ii) Sample overlap: seven-dimensional coefficient-summary blocks were generated directly at every $M$, with $\Sigma_X = \Sigma_Y = I_7/(7F_{\text{target}})$ and $\Sigma_{XY} = \rho_{\text{eff}} I_7/(7F_{\text{target}})$. At $F_{\text{target}} \in \{1.5, 7, 15, 30\}$ and $\theta = 0.25$, the main grid was $M \in \{20, 100, 500\}$ and overlap strength $\rho_{\text{eff}} \in \{0, 0.2, 0.4, 0.6\}$, comparing the independence and overlap-aware objectives (500 replicates per setting); an extended grid at $F_{\text{target}} = 30$ with $M \in \{1, 5, 10, 20, 50, 100, 500\}$ reports the coverage of Wald intervals from curvature standard errors (Supplementary Table 1f(iv)). (iii) Correlated local blocks: $M = 10$ and $n = 3{,}000$, with $Z_m = \sqrt{n}\{c^{1/2}U + (1-c)^{1/2}U_m\}$ for shared and block-specific components $U$ and $U_m$, giving observed maximum canonical correlations of 0.05, 0.20, 0.50, 0.80 and 0.95; at $\theta = 0.25$ and $F_{\text{target}} \in \{1.5, 7, 15, 30\}$, an unpruned block-diagonal analysis, which ignores off-diagonal coefficient covariance, was compared with an analysis of one pre-specified representative block per correlated cluster (the ‘pruned top hit’) and a full-covariance comparator (1,000 replicates per setting).

*Sample overlap.* Independence-objective coverage fell as blocks were added. At correlation 0.6 and $F_{\text{target}} = 7$, it was 0.420 with $M = 20$ blocks, 0.000 with $M = 100$ and 0.000 with $M = 500$, and at $F_{\text{target}} = 30$ it was 0.000 with $M = 500$. With the overlap covariance known, the overlap-aware objective covered 0.932–0.954 at that correlation for $M$ up to 500 and $F_{\text{target}}$ of 7 and 15 (Supplementary Table 1f(iii); Fig. 3e; Extended Data Fig. 4c). In the extended $F_{\text{target}}$ = 30 grid, Wald coverage at correlation 0.6 and $M = 500$ was 0.000 (independence) versus 0.936 (overlap-aware; Supplementary Table 1f(iv)). The estimating-equation bias behind this is derived in Supplementary Note 1, Section 1.5.

*Correlated neighbouring loci and pruning.* Treating correlated neighbouring blocks as independent reduced unpruned SO coverage to 0.415–0.468 at canonical correlation 0.95 over $F_{\text{target}}$ of 1.5, 7 and 15, and to 0.463 at $F_{\text{target}} = 30$. At strengths 7 and 15, with the block covariances known, the pruned top hit gave coverage 0.935–0.958, similar to the full-covariance comparator (0.921–0.957); the first-order block-diagonal analysis under-covered (Supplementary Table 1f(ii); Fig. 3f; Extended Data Fig. 4e). Pruning did not remove weak-instrument behaviour: at $F_{\text{target}} = 1.5$, the retained single block covered 0.855–0.921, with 7.2% unbounded sets.

*Weak instruments and same-sample selection.* At $F_{\text{target}} = 1.5$ without selection, the overlap-aware estimate of the sample-overlap arm remained approximately centred (largest absolute mean relative error 3.3%) but covered only 0.846–0.906, and the independent-block arm covered 0.875 and 0.907 at $M = 20$ and 100 (mean over $\theta$; Supplementary Tables 1f(i) and 1f(iii)). This shortfall is the finite-sample behaviour of many weak blocks, not an effect of selection. Same-sample selection removed on average 17.5% of blocks in the sample-overlap arm and introduced bias: mean relative error was −11.8% to −8.9% without overlap and +10.7% to +12.6% at correlation 0.6, where the slope of the correlated estimation errors exceeds $\theta = 0.25$. Coverage fell as blocks were added, to 0.632 and 0.294 at $M = 500$ (Supplementary Table 1f(iii)); selection also worsened coverage in the independent-block arm (Supplementary Table 1f(i)). Overlap adjustment does not remove this founder-block form of the winner's curse[6].

#### 4.5 Cross-locus diagnostics (S5)

*Design.* Seven-dimensional coefficient-summary blocks were generated directly, with random positive-definite $\Sigma_{X,m}$ and $\Sigma_{Y,m}$ and $\Sigma_{XY,m} = 0$. With $M = 20$, $\theta = 0.25$ and every locus at the same $F_{\text{target}} \in \{1.5, 7, 15, 30\}$ (200 replicates per scenario), a clean scenario ($b_{Y,m} = \theta b_{X,m}$ for all $m$) was compared with a directional-invalid scenario in which two high-information loci followed $b_{Y,m} = (\theta + c_m) b_{X,m}$ with $c_m = \theta$: internally proportional blocks with shifted local effects. Three further scenarios placed the same two loci under orthogonal within-locus contamination ($\lambda = 1$), combined a smaller directional shift ($c_m = \theta/2$) with orthogonal contamination ($\lambda = 0.75$), or used a clean proportional design with $\theta = 1$ (Extended Data Fig. 5). We evaluated global SO, per-locus SO, two-step IVW, weighted median, weighted mode, $Q_{\text{between}}$ and leave-one-locus-out influence; the representative replicate of Fig. 3g was chosen by the rule given in Methods.

At $F_{\text{target}} = 15$, the directional scenario raised global SO mean error from 0.002 to 0.052 and reduced coverage from 0.930 to 0.365. Two-step IVW also shifted, whereas weighted median and mode estimates[7,8] stayed close to $\theta$. $Q_{\text{between}}$ rejected in 0.66 of replicates (0.06 when clean), and leave-one-out refits ranked a planted locus as most influential in 0.99. Diagnostic power depended on strength: $Q_{\text{between}}$ rejection rose from 0.39 at $F_{\text{target}} = 7$ to 0.94 at 30. Within-locus contamination behaved differently: orthogonal contamination left global SO coverage at 0.940 with near-nominal $Q_{\text{between}}$ rejection (0.06), whereas mixed contamination reduced coverage to 0.690 with rejection 0.23 (Supplementary Table 1g; Extended Data Fig. 5). Greater strength did not correct invalid loci: in the directional scenario at $F_{\text{target}} = 30$, global SO coverage was 0.095 (Supplementary Table 1g).

In clean S5 at $F_{\text{target}} = 1.5$, $\theta = 0.25$ and with every locus valid, two-step IVW, weighted median and weighted mode estimates were biased towards zero both with and without selection, because each per-locus estimate is itself a weak ratio. Global SO mean error was 0.010 without selection and $-0.019$ with selection (coverage 0.885 and 0.870). At $\theta = 1$, the corresponding errors were 0.003 and $-0.086$ (Supplementary Table 1g; Extended Data Fig. 5).

##### 4.6 Accuracy of the star-space reconstruction

*Design.* A phenotype-level simulation used $n_X = n_Y = 140$ mice, a synthetic kinship matrix $K$ for the union of the samples (with exposure-by-outcome block $K_{XY}$), a simulated founder-contrast design, $\theta = 0.25$ and $F_{\text{target}} = 100$. Three known residual cross-covariances were compared: the in-model target $\Delta_* = \rho\,\sigma_X\sigma_Y S_X^{-1} O S_Y^{-1}$ of Supplementary Note 3; the raw-overlap target $\Delta_{\text{raw}} = \rho\,\sigma_X\sigma_Y O$; and the kinship-plus-residual target $\Delta_{\text{kin+raw}} = \sigma_X\sigma_Y\{(\rho/2)(2K_{XY}) + (\rho/2)O\}$. The grid crossed overlap fractions 0.3, 0.6 and 1.0, heritability pairs $(h_X^2, h_Y^2) = (0.20, 0.20)$, $(0.78, 0.50)$ and $(0.78, 0.80)$, and nominal correlations 0.05, 0.10 and 0.20, with 50 replicates per setting (81 settings). Overlap-aware SO fits using the reconstructed and the exact coefficient-level cross-covariances were compared (Supplementary Table 6). The coefficient-fitting and whitening maps used the true heritabilities, so uncertainty in estimated $h_X^2$ and $h_Y^2$ was not tested. The exposure and outcome founder-probability matrices were drawn independently, even for overlapping mice, which weakened the coefficient-level cross-covariance relative to mice measured twice: true $s_{\max}$ was 0.01–0.08 for the in-model target and at most 0.39 across all targets.

Covariance recovery was imperfect: the relative Frobenius error $\|\widehat{\Sigma}_{XY} - \Sigma_{XY}^{\text{exact}}\|_F / \|\Sigma_{XY}^{\text{exact}}\|_F$ had mean 0.58 to 1.09 across targets, and was large even for the in-model target, where small nominal correlations made $\widehat{\rho}_*$ noisy relative to its target. The resulting SO estimates nevertheless differed by at most $5.61 \times 10^{-3}$ in any replicate, and coverage by at most 0.020 in any setting.

### Supplementary Note 5. Additional methods

This Note gives the data, instrument-selection and computational details that Methods summarises.

#### 5.1 Upstream data and scans

*Data and upstream scan.* The analyses use the Diversity Outbred bone cohort of the Farber laboratory[9]. The cohort data were supplied with that laboratory's R/qtl2 genome scan, which used rank inverse-normal-transformed traits, $\Phi^{-1}\{(r - 0.5)/n\}$ for rank $r$ among $n$ mice with average ranks for ties, leave-one-chromosome-out kinship and the covariates sex, age at sacrifice, body weight, body length and generation as a numeric term, in mice with complete values for these covariates.

*Use for instrument selection, and the body-size rescan.* We used the upstream scan only to select instruments: it defined the candidate loci and their representative markers for every exposure except body weight and body length. Because the scan adjusted for body weight and body length, we rescanned these two traits without them, with leave-one-chromosome-out kinship, the covariates sex, age at sacrifice and generation as a factor, and the same rank transform, and took their candidate instruments from this rescan. Instruments selected in a scan adjusted for body size could be affected by that adjustment, a collider-type selection. The rescan shares its covariates and sample with our fits, and all of its instruments passed the inclusion threshold, so the instrument strengths of these two exposures are not comparable with those of the other exposures.

*Downstream estimates.* All downstream estimates, including instrument strength, founder support and every MR effect, come from our own linear mixed models (Methods). These models can remove candidate loci but cannot add loci that the scan did not select.

#### 5.2 Instrument selection

*Peak rule.* On each autosome, with markers in map order, peaks were called with `find_peaks` of R/qtl2[10] at `threshold = 4` and `peakdrop = 1.5`. Each peak is represented by its first maximum-LOD marker in map order.

*Alternative instrument lists.* The peak-drop list applies the same rule with `peakdrop = 2`. The contiguous-run list keeps the highest marker, removes the contiguous run of markers with LOD $\geq 4$ around it and repeats, and then visits these candidates in decreasing LOD, dropping any within 1 Mb of one already kept on the same chromosome.

*Inclusion threshold.* $\widehat{F}_X^{\text{block}} \geq 1.5$ is lenient: a block with no exposure signal passes it with probability $\Pr(\chi^2_7 \geq 10.5) \approx 0.16$.

#### 5.3 Scalar SNP effects

The SNP effects of the scalar SNP-IVW (S) in the landscape and ML–pMOI analyses were estimated for each trait separately by generalised least squares on the centred observed dosage at each instrument's marker, using the trait's own sample, covariates and LOCO kinship, with variance components estimated by REML under the null model for each trait and chromosome, and expressed per SD of the trait on its own sample. Missing dosage that remained after the missing-dosage rule of Methods was mean-imputed within each trait's own sample, and missing outcome dosage never removed a locus; founder probabilities are complete. A marker with minor allele frequency below 0.01 in a trait's own sample gave no estimate. Scalar SNP effects were not computed at markers that appear only in the contiguous-run list, so the landscape analysis with that list compared F1 and F2 only.

#### 5.4 Traits and the *Qsox1* analysis

*Traits.* Bone traits follow standard nomenclature[11]: cortical tissue mineral density (TMD), polar moment of inertia (pMOI), maximum moment of inertia (Imax), cortical area fraction (Ct.Ar/Tt.Ar), total area (Tt.Ar), medullary area (Ma.Ar) and cortical porosity (Ct.Por), all measured by micro-computed tomography ($\mu$CT), and medial–lateral femoral width (ML)[9]. The eight outcomes of the *Qsox1* analysis were TMD, ML, pMOI, Imax, Ct.Ar/Tt.Ar, Tt.Ar, Ma.Ar and Ct.Por. The 74 landscape traits span bone morphology and $\mu$CT microarchitecture, histomorphometry, biomechanics, adiposity, body size, glucose, muscle mass and marrow adipose tissue (MAT) volumes.

*Marrow adipose tissue.* The four MAT volumes, zeros included, were rank-transformed on each trait's own complete sample before the mixed models, whether the trait was the exposure or the outcome, using Blom's offsets, $\Phi^{-1}\{(r-3/8)/(n+1/4)\}$, with average ranks for ties.

*Expression. Qsox1* expression followed the processing of the original study[9]: read counts were variance-stabilised with DESeq2[12], quantile inverse-normal transformed, and 48 probabilistic estimation of expression residuals (PEER) factors were estimated from the transformed values without intercept or covariates[13]. We used the expression values and PEER factors distributed with the published cross files, without re-estimation.

*Instrument.* The lead *Qsox1* eQTL marker `JAX00272363` (155.654 Mb) had a founder-probability eQTL LOD of 24.42, above the autosomal permutation threshold of the original study (10.89)[9].

#### 5.5 Multi-locus analysis of ML and pMOI

*Refits.* Besides the leave-one-locus-out and chromosome-exclusion refits, the global SO estimate was refitted on the top-$k$ blocks by $\widehat{F}_X^{\text{block}}$ ($k = 10$, 25 and 50) and after exclusion of the five most influential blocks.

*Weighted median and weighted mode.* The weighted median is the lower weighted median, and the weighted mode is the maximum of a weighted normal-kernel density. Their Wald intervals use as standard error the standard deviation of the estimate over 500 parametric bootstrap draws, each redrawing every $\widehat{\theta}_m$ from $N\{\widehat{\theta}_m, \widehat{\text{Var}}(\widehat{\theta}_m)\}$ with loci and weights fixed. Locus-bootstrap percentile intervals, computed with a different weighted-mode bandwidth rule, are in Supplementary Table 9.

*Comparators.* The SNP effects of the scalar SNP-IVW were estimated as in Section 5.3. The observational regression used the traits standardised on their own samples; the unadjusted regression is in Supplementary Table 9.

#### 5.6 Landscape computation

For each exposure, the founder fits were computed once, in a single batch call over the union of the primary, peak-drop and contiguous-run lists without an inclusion threshold, and the fitted locus blocks were stored; the analyses with $\widehat{F}_X^{\text{block}} \geq 1.5$ or $\geq 2$, with or without the Steiger condition and for each instrument list, were re-aggregated from these blocks.

#### Supplementary Note 6. Basis-rotation decomposition

*First classification.* Before the decomposition, the 170 scalar-only pairs were classified under rules fixed before the results were inspected, and the pair with opposite-sign estimates was described locus by locus (rules and cut-offs, Supplementary Table 8). Every pair that changed at the representation step had a founder estimate less than half the scalar estimate, or of opposite sign, with a standard error less than 1.25 times that of the scalar estimate (for 51 of the 123, at least half of the scalar signal came from loci with discordant founder patterns or from a single locus). Of the 47 that changed at the overlap-aware step, the interval widened by at least 30% for 15, the estimate moved towards or past zero for 12, and the interval just reached zero with little change for 20.

The follow-up covers the 123 scalar-only pairs that lost the scalar signal at the representation step and the pair with opposite-sign estimates (Methods), with reference groups and the 47 pairs that lost it at the overlap-aware step for comparison (Section 6.6). It asks whether the scalar signal is carried by the one founder direction that the marker SNP represents, and whether the rest of the founder block carries exposure signal that the outcome does not follow. The analysis was specified before it was run; two deviations and one added flag are stated in Section 6.5. Values are in Supplementary Table 10.

For a pair, the loci $m = 1, \dots, M$ are those of its set $L$ (Methods), with summaries $\widehat{b}_{X,m}$, $\widehat{b}_{Y,m}$, $\Sigma_{X,m}$, $\Sigma_{Y,m}$ and $\Sigma_{XY,m}$. The marker of locus $m$ has strain distribution $s_m \in \{0,1\}^8$, the founder alleles at the SNP with either allele coded 1. Loci whose marker has a missing or heterozygous founder call, or is monomorphic among the founders, are excluded from the subspace analyses; the remaining $M'$ loci are the complete loci, and a pair is flagged when more than 20% of its loci are excluded. The locus index is dropped where a statement holds locus by locus.

##### 6.1 The SNP axis

The contrast vector of the marker is $c = H^\mathsf{T}(s - \bar{s}\mathbf{1}_8) \in \mathbb{R}^7$, with $\bar{s}$ the mean entry of $s$. Because $\mathbf{1}_8^\mathsf{T} a = 0$ for founder effects $a = Hb$,

$$c^\mathsf{T} b = a^\mathsf{T}(s - \bar{s}\mathbf{1}_8) = \sum_{f:\, s_f = 1} a_f = -\sum_{f:\, s_f = 0} a_f, \tag{S27}$$

the sum of the founder effects in the group coded 1, or minus the sum over the other group.

*Transformation.* Let $\Sigma_Y^{-1/2}$ be the symmetric inverse square root, $u_1 = \Sigma_Y^{-1/2} c / \|\Sigma_Y^{-1/2} c\|$, $U_2 \in \mathbb{R}^{7\times 6}$ an orthonormal basis of the orthogonal complement of $u_1$ and $R = [u_1, U_2]$. For $T \in \{X, Y\}$,

$$J = R^\mathsf{T} \Sigma_Y^{-1/2}, \qquad \gamma_T = J\widehat{b}_T, \qquad \Sigma'_T = J\Sigma_T J^\mathsf{T}, \qquad \Sigma'_{XY} = J\Sigma_{XY} J^\mathsf{T}, \tag{S28}$$

so that $\Sigma'_Y = I_7$. $J$ is whitening in the first-order metric $\Sigma_Y^{-1}$ followed by a rotation; it is not an orthogonal map of the original coordinates. Write $b = \alpha c + r$ with $c^\mathsf{T}\Sigma_Y^{-1} r = 0$, the generalised least-squares (GLS) decomposition in this metric. Then $Jb$ has coordinates

$$(Jb)_1 = \alpha \|\Sigma_Y^{-1/2} c\|, \qquad \alpha = \frac{c^\mathsf{T}\Sigma_Y^{-1} b}{c^\mathsf{T}\Sigma_Y^{-1} c}, \qquad (Jb)_{2:7} = U_2^\mathsf{T}\Sigma_Y^{-1/2} r, \qquad \|(Jb)_{2:7}\|^2 = r^\mathsf{T}\Sigma_Y^{-1} r. \tag{S29}$$

The other six coordinates carry everything in $b$ that is orthogonal to $c$ in this metric, and they vanish if and only if $b \in \operatorname{span}(c)$. No result below depends on the choice of $U_2$.

*A marker regression within the founder model.* The scalar genotype of the marker is $g_s = Ps$ (Methods). Because $s - \bar{s}\mathbf{1}_8 = Hc$, $g_s = \bar{s}P\mathbf{1}_8 + Zc$, whose first term is constant, so $M_T g_s = M_T Zc$

for the residual-making matrix $M_T$ of Methods. Fitted with the same samples, covariates and covariance as the founder block, the marker regression (S15) therefore gives, using $Z^\top M_T T = (Z^\top M_T Z)\widehat{b}_T$ and $\Sigma_T \propto (Z^\top M_T Z)^{-1}$,

$$\widehat{\beta}_{g_s T} = \frac{c^\top Z^\top M_T T}{c^\top Z^\top M_T Z c} = \frac{c^\top \Sigma_T^{-1} \widehat{b}_T}{c^\top \Sigma_T^{-1} c} \tag{S30}$$

for any founder imbalance. For the outcome this is $\alpha$ of (S29): the first axis is exactly what a marker regression within the founder model estimates.

*Ratios.* If $\Sigma_X \propto \Sigma_Y$ at a locus, with any locus-specific constant, the within-model marker ratio is

$$\theta_{\mathrm{S},m} = \frac{c_m^\top \Sigma_{Y,m}^{-1} \widehat{b}_{Y,m}}{c_m^\top \Sigma_{Y,m}^{-1} \widehat{b}_{X,m}} = \frac{\gamma_{Y1,m}}{\gamma_{X1,m}}, \tag{S31}$$

and the scalar SNP-IVW estimate of Methods, computed from these marker regressions, is $\sum_m \phi_m \gamma_{X1,m}\gamma_{Y1,m} / \sum_m \phi_m \gamma_{X1,m}^2$, where $\phi_m$ is the ratio of the founder-model to the marker-model residual variance of the outcome (the marker residual absorbs the founder signal outside $c_m$). Apart from the weights $\phi_m$, this is the first-order estimate on the SNP axis (row (i), Section 6.2).

The S analysis approximates this marker regression: it uses the observed array dosage instead of $g_s$, its own fit and scaling, and all $M$ loci rather than the $M'$ complete loci. The ratio of the row-(i) first-order estimate to the S estimate, both on the complete loci, is reported for each pair.

#### 6.2 Invariance and the subspace estimators

*Invariance.* $J$ is invertible and $J^\top J = \Sigma_Y^{-1}$, so $\widehat{b}_X^\top \Sigma_Y^{-1} \widehat{b}_Y = \gamma_X^\top \gamma_Y$ at every locus, and the first-order estimate and its standard error (S10) become

$$\widehat{\theta}_{\mathrm{FO}} = \frac{\sum_m \gamma_{X,m}^\top \gamma_{Y,m}}{\sum_m \|\gamma_{X,m}\|^2}, \qquad \widehat{\mathrm{SE}}_{\mathrm{FO}} = \Big(\sum_m \|\gamma_{X,m}\|^2\Big)^{-1/2}. \tag{S32}$$

Under $d_m \mapsto J_m d_m$ and $\Omega_m \mapsto J_m \Omega_m J_m^\top$, each term $d_m^\top \Omega_m^{-1} d_m$ of the second-order objective is unchanged for any invertible $J_m$, as for a change of contrast basis (Supplementary Note 2, Section 2.1). Hence $\widehat{\theta}$, $Q^{\min}$, the profile-$Q$ set and the curvature standard error are unchanged. The orthogonal case is the basis-invariance check of Supplementary Table 5.

*The grid.* Three estimators are applied to four sets of blocks. The rows are (i) the SNP axis: coordinate 1 of $\gamma_{X,m}$ and $\gamma_{Y,m}$, with the $[1,1]$ entries of $\Sigma'_{X,m}$, $\Sigma'_{Y,m}$ and $\Sigma'_{XY,m}$; (ii) the other six: coordinates 2–7, with the $[2{:}7, 2{:}7]$ sub-blocks; (iii) all seven coordinates on all $M$ loci, which reproduces F1 and F2; and (iii′) all seven coordinates on the $M'$ complete loci. The columns are the first-order estimator (FO, the F1 tier), the second-order estimator with $\Sigma_{XY} = 0$ (SO-independence; Supplementary Note 1, Section 1.5) and the overlap-aware second-order estimator (SO overlap-aware, the F2 tier), each with its interval as in Methods. Row (i) uses the one-dimensional objective

$$Q_1(\theta) = \sum_{m=1}^{M'} \frac{(\gamma_{Y1,m} - \theta\gamma_{X1,m})^2}{1 + \theta^2 \sigma'_{X,m} - 2\theta\sigma'_{XY,m}}, \tag{S33}$$

where $\sigma'_{X,m}$ and $\sigma'_{XY,m}$ are the $[1,1]$ entries of $\Sigma'_{X,m}$ and $\Sigma'_{XY,m}$ ($\sigma'_{XY,m} = 0$ for SO-independence), minimised over the whole real line with the profile set of Supplementary Note 1, Section 1.4. Under $b_Y = \theta b_X$ every row targets $\theta$; otherwise each row targets its own subspace ratio, $\theta_1$ for row (i) and $\theta_2$ for row (ii).

*First-order decomposition.* Because $\Sigma'_Y = I_7$, the first-order objective on the complete loci is the sum of its row-(i) and row-(ii) parts. With $D_1 = \sum_m \gamma^2_{X1,m}$, $D_2 = \sum_m \|\gamma_{X,2:7,m}\|^2$ and $\widehat{\theta}_1$, $\widehat{\theta}_2$ the first-order estimates of rows (i) and (ii),

$$\widehat{\theta}^{(\text{iii}')}_{\text{FO}} = w\,\widehat{\theta}_1 + (1-w)\,\widehat{\theta}_2, \qquad w = \frac{D_1}{D_1 + D_2}. \tag{S34}$$

This holds for row (iii′), and for F1 only when $M' = M$. Conditional on $\gamma_X$, the reported first-order variances satisfy $\operatorname{Var}(\widehat{\theta}^{(\text{iii}')}_{\text{FO}})/\operatorname{Var}(\widehat{\theta}_1) = w$ exactly, so the first-order interval from all seven coordinates is $w^{1/2}$ times as wide as that from the SNP axis alone.

*No second-order decomposition.* The off-diagonal block of $J\Omega(\theta)J^{\mathsf{T}}$, $\theta^2\Sigma'_X[1,2{:}7] - \theta(\Sigma'_{XY}[1,2{:}7] + \Sigma'_{XY}[2{:}7,1]^{\mathsf{T}})$, is non-zero in general, so the second-order objective is not the sum of its row-(i) and row-(ii) parts.

*Dependence.* Conditional on $\gamma_X$, the first-order $\widehat{\theta}_1$ and $\widehat{\theta}_2$ are uncorrelated because $\Sigma'_Y = I_7$; unconditionally they are correlated through $\Sigma'_X[1,2{:}7]$ and the cross blocks of $\Sigma'_{XY}$. Their correlation $\rho_{12}$ is therefore computed for every column (Section 6.5).

#### 6.3 Information share and concordance

*Exposure signal.* At each complete locus, the Wald statistics of the two parts are

$$W_1 = \frac{\gamma^2_{X1}}{\Sigma'_X[1,1]}, \qquad W_2 = \gamma^{\mathsf{T}}_{X,2:7}\big(\Sigma'_X[2{:}7,2{:}7]\big)^{-1}\gamma_{X,2:7}, \tag{S35}$$

non-central $\chi^2_1$ and $\chi^2_6$ variables. $W_2$ does not depend on the whitening metric: it is the Wald statistic for the restriction $b_X \in \operatorname{span}(c)$, $W_2 = \min_\alpha(\widehat{b}_X - \alpha c)^{\mathsf{T}}\Sigma_X^{-1}(\widehat{b}_X - \alpha c)$.

*First-order information.* $D_k$ is the first-order precision, or information, of part $k$ in the outcome-noise metric: conditional on $\gamma_X$, $\operatorname{Var}(\widehat{\theta}_k) = 1/D_k$ when exposure noise and $\Sigma'_{XY}$ are ignored. Let $\bar{\gamma}_X = Jb_X$ be the noise-free coordinates and $\bar{\gamma}_{X,k}$ their part $k$. Because $\mathbb{E}(D_k) = \sum_m \|\bar{\gamma}_{X,k,m}\|^2 + \sum_m \operatorname{tr}\Sigma'_{X,m}[k,k]$, the noise-corrected precisions are $D^*_k = D_k - \sum_m \operatorname{tr}\Sigma'_{X,m}[k,k]$, and the noise-corrected information share of the SNP axis is

$$\pi_D = \frac{D^*_1}{D^*_1 + D^*_2}, \quad \text{which estimates} \quad \Pi = \frac{\sum_m \bar{\gamma}^2_{X1,m}}{\sum_m \|\bar{\gamma}_{X,m}\|^2} = \frac{\sum_m \|\bar{\gamma}_{X,m}\|^2\pi_m}{\sum_m \|\bar{\gamma}_{X,m}\|^2}, \tag{S36}$$

where $\pi_m = \bar{\gamma}^2_{X1,m}/\|\bar{\gamma}_{X,m}\|^2$ is the noise-free share at locus $m$, the squared cosine between $\bar{\gamma}_{X,m}$ and the first axis. $D^*_k$ can be negative; $\pi_D$ is not truncated and is reported only when $D^*_1$ and $D^*_2$ are both positive. $\pi_D$ is the noise-corrected counterpart of the first-order weight $w$ in (S34), the uncorrected information share, and the two agree only when $D_k \gg \sum_m \operatorname{tr}\Sigma'_{X,m}[k,k]$. Exposure noise enters $D_2$ through six dimensions and $D_1$ through one, so it pulls $w$ towards 1/7; with weak blocks the gap is material.

*Concordance within each part.* For part $k$ ($k = 1$, the SNP axis, with $d_1 = 1$ coordinate; $k = 2$, the other six, with $d_2 = 6$), let $\gamma_{T,k,m}$ be the coordinates of part $k$ at complete locus $m$, and

$$\begin{gathered}\Gamma_{XY,k} = \sum_m\big\{\gamma^{\mathsf{T}}_{X,k,m}\gamma_{Y,k,m} - \operatorname{tr}\Sigma'_{XY,m}[k,k]\big\}, \qquad \Gamma_{YY,k} = \sum_m \|\gamma_{Y,k,m}\|^2 - d_kM', \\ \Gamma_{XX,k} = D^*_k, \qquad r_k = \operatorname{sign}(\widehat{\theta}_{\text{S}})\,\frac{\Gamma_{XY,k}}{(\Gamma_{XX,k}\Gamma_{YY,k})^{1/2}},\end{gathered} \tag{S37}$$

defined when $\Gamma_{XX,k} > 0$ and $\Gamma_{YY,k} > 0$. The concordance index $r_k$ is an uncentred cosine between the exposure and outcome founder-effect patterns in part $k$, corrected for noise and overlap and oriented by the sign of the S estimate: 1 means proportional in the direction of S, and 0 no proportional relation. Because of the noise correction, $|r_k|$ can exceed 1; values are not clipped, and no per-pair uncertainty is computed.

##### 6.4 Amplification of non-proportional effects, and compensation

This section compares the probability limits of the scalar and the founder first-order estimators; finite-sample weak-instrument bias and the second-order estimator are not covered. At locus $m$ let $b_{Y,m} = \theta b_{X,m} + \delta_m$, where $\delta_m$, the part of the outcome effect not proportional to the exposure effect, reflects pleiotropy or a second QTL in the block; a component proportional to $b_{X,m}$ is indistinguishable from a change in $\theta$ and is absorbed into $\theta$. The scalar is taken to be the within-model marker ratio (S31), which assumes $\Sigma_X \propto \Sigma_Y$ at the locus; for row (i) the expressions below are exact without this assumption. With $\tilde{\delta} = J\delta$,

$$\theta_{\mathrm{S},m} = \theta + \frac{\tilde{\delta}_1}{\bar{\gamma}_{X1}}, \qquad \theta_{\mathrm{F},m} = \frac{b_X^\top \Sigma_Y^{-1} b_Y}{b_X^\top \Sigma_Y^{-1} b_X} = \theta + \frac{\bar{\gamma}_X^\top \tilde{\delta}}{\|\bar{\gamma}_X\|^2}; \tag{S38}$$

the first expression is (S16) in the rotated coordinates. Since $|\bar{\gamma}_{X1}| = \pi_m^{1/2}\|\bar{\gamma}_X\|$, the scalar bias is at most $\pi_m^{-1/2}\|\tilde{\delta}\|/\|\bar{\gamma}_X\|$ and the founder bias at most $\|\tilde{\delta}\|/\|\bar{\gamma}_X\|$: over $\tilde{\delta}$ of fixed norm, the worst-case scalar bias is $\pi_m^{-1/2}$ times the worst-case founder bias. For a given $\delta$, the ratio of the two biases is unbounded in both directions.

*Random isotropic pleiotropy.* Suppose that the $\delta_m$ are independent, have mean zero, are independent of $b_X$ and satisfy $\mathbb{E}(\delta_m\delta_m^\top) = \tau^2\Sigma_{Y,m}$ with one $\tau$ for all loci (isotropy in the whitened metric; the multiplicative random-effects form). Then $\mathbb{E}(\tilde{\delta}_1^2) = \mathbb{E}\|\tilde{\delta}\|^2/7 = \mathbb{E}\{(v^\top\tilde{\delta})^2\}$ for the unit vector $v = \bar{\gamma}_X/\|\bar{\gamma}_X\|$, and the root-mean-square (RMS) ratio of the scalar to the founder IVW bias is $\Pi^{-1/2}$, with $\Pi$ of (S36): exactly for the row-(i) first-order estimate, and up to the weights $\phi_m$ for the scalar SNP-IVW estimate. If $\tau$ varies across loci, the scale is common in raw units, or the $\delta_m$ are directional, there is no general factor, so $\pi_D^{-1/2}$ is an RMS ratio under these conditions only, not a typical or maximal factor.

*Compensation.* By (S27), $c^\top b_X$ is the sum of the exposure founder effects in the group coded 1, so founder effects of opposite sign within a group cancel, and $c^\top b_X$ can be small at a locus with strong founder-level effects ($\pi_m$ is governed by $c^\top\Sigma_Y^{-1}b_X$, which need not vanish when $c^\top b_X$ does). The scalar's noise-free first-stage non-centrality is $\pi_m^{(X)} b_X^\top\Sigma_X^{-1}b_X$, with $\pi_m^{(X)}$ the share in the exposure metric, so the scalar instrument is weak when the share is small, and a $\delta$ that does not cancel in the same way is amplified by the factors above. The founder estimator uses all seven directions and is not exposed to this cancellation.

##### 6.5 Tests and classification

*Exposure signal outside the SNP direction.* $P_{\text{other}}$ refers $\sum_m W_{2,m}$ to $\chi^2_{6M'}$, under the null hypothesis that $b_{X,m} \in \operatorname{span}(c_m)$ at every complete locus; $P_{\text{SNP}}$ refers $\sum_m W_{1,m}$ to $\chi^2_{M'}$. $P_{\text{other}}$ uses exposure data only and has four limits. (1) Loci are treated as independent, but marginal blocks fitted on the same mice are correlated through haplotype linkage disequilibrium; because $\operatorname{Cov}(z_j^\top A z_j, z_k^\top B z_k) = 2\operatorname{tr}(A\Sigma_{jk}B\Sigma_{kj}) \geq 0$ for jointly Gaussian mean-zero $z_j$, $z_k$ and positive semidefinite $A$, $B$, this can only inflate the null variance of $\sum_m W_{2,m}$, so $P_{\text{other}}$ is anticonservative. (2) The plug-in $\Sigma_X$ is treated as known, which inflates each Wald statistic (by about $n/(n-p)$ for a maximum-likelihood residual variance with $p$ fixed-effect columns) and, when $\widehat{\Sigma}_X$ is biased downwards, tilts $\pi_D$ towards the other six directions. (3) A large $W_2$ means only that $b_X$ does not lie in the marker direction, which a multi-allelic QTL[5], a marker whose strain distribution is not the causal one or leakage of other loci's effects can each produce. (4) Pairs that share an exposure share, or nearly share, $P_{\text{other}}$.

*Does the outcome follow in the other six directions?* Each subspace estimate solves an estimating equation $g_k(\theta) = 0$, the first-order normal equation or the second-order score of row (i) or (ii). Linearising both in the per-locus data,

$$\mathrm{Cov}(\widehat{\theta}_1, \widehat{\theta}_2) \approx \frac{1}{h_1 h_2} \sum_m \nabla g_{1,m}^{\mathsf{T}} V_m \nabla g_{2,m}, \qquad h_k = \frac{\partial g_k}{\partial \theta}\Big|_{\widehat{\theta}_k}, \tag{S39}$$

where $\nabla g_{k,m}$ is the gradient with respect to $(\gamma_{X,m}, \gamma_{Y,m})$ and $V_m$ their joint covariance, built from $\Sigma'_{X,m}$, $I_7$ and $\Sigma'_{XY,m}$. The same linearisation gives the variances, the correlation $\rho_{12}$ and the delta-method standard deviation $\mathrm{SD}_\delta$ of $\widehat{\theta}_2 - \widehat{\theta}_1$, which includes exposure noise. The primary statistic is

$$z_c = \frac{\widehat{\theta}_2 - \widehat{\theta}_1}{\mathrm{SD}_\delta(\widehat{\theta}_2 - \widehat{\theta}_1)}. \tag{S40}$$

The pre-specified form, $z_c^{\text{pre}} = (\widehat{\theta}_2 - \widehat{\theta}_1)/(\mathrm{se}_1^2 + \mathrm{se}_2^2 - 2\rho_{12}\mathrm{se}_1\mathrm{se}_2)^{1/2}$, with each cell's own standard error (curvature for SO, $D_k^{-1/2}$ for FO) and the delta-method $\rho_{12}$, is reported alongside. The primary column is SO overlap-aware, as specified. Before the full analysis, the delta method was checked against a parametric bootstrap (200 draws per pair) in three pilot pairs; the specified criterion was agreement of the standard deviation of $\widehat{\theta}_2 - \widehat{\theta}_1$ within 15% in each column.

*Deviations from the specification.* (1) $z_c$ replaced $z_c^{\text{pre}}$ as the primary statistic, because the check validated the delta-method standard deviation, not the denominator of $z_c^{\text{pre}}$, which combines model standard errors with a delta-method covariance. In the three pilot pairs both denominators were below the robust bootstrap spread of $\widehat{\theta}_2 - \widehat{\theta}_1$, that of $z_c^{\text{pre}}$ by more than that of $z_c$. The change was made before the full and reference analyses were run, with both statistics of the pilot pairs visible; it lowered $|z_c|$ in each pilot pair, taking two of the three below 1.96. Both statistics are reported. (2) The SO columns were read against a robust bootstrap spread, estimated from the median absolute deviation, because the row-(i) SO estimator is of ratio type and its bootstrap distribution is heavy-tailed. Under the standard deviation none of the nine column-by-pair checks passed; under the robust spread the overlap-aware column passed in all three pairs and the independence column in two. The FO column failed and was set aside, because its bootstrap, centred on the noisy $\gamma_X$, adds exposure noise twice. *Added flag:* pairs whose SNP axis has no detectable exposure signal ($P_{\text{SNP}} \geq 0.05$) are flagged as having a low-reliability $z_c$; the flag was fixed after the pilot and before any full or reference classification was viewed.

*Selection.* Within the selected pairs, $z_c$ is inflated by the selection: with $M' = M$, (S34) gives $\widehat{\theta}_2 = (\widehat{\theta}_{\text{F1}} - w\widehat{\theta}_1)/(1 - w)$, so selecting an S interval that excludes zero and an F1 interval that includes it pulls $\widehat{\theta}_2$ towards zero or past it, the direction criterion of class C below. As a baseline, the same analysis was applied to 100 pairs drawn at random, with a seed fixed before the run, from each of two reference groups of the QC cohort: pairs whose S and F2 intervals excluded zero with the same sign (R1), and founder-only pairs (R2).

*Overlap amplification (OA).* On the SNP axis (row (i)), the FO or the SO-independence interval excludes zero while the SO overlap-aware profile set contains it, containment being decided by membership, $Q(0) - Q^{\min} \leq \chi^2_{1,0.95}$ (with $Q_1$), whatever the shape of the set. The containment leg is conservative to the extent that the profile set is too narrow; the exclusion leg is anticonservative, because FO standard errors omit exposure noise, the SO-independence standard error can be understated at weak strength, and two chances are taken. The net direction of the error in OA counts is not established.

*Classification.* Each pair is classified by rules fixed before the analysis was run. Compression (C) requires $P_{\text{other}} < 0.05$, $|z_c| \geq 1.96$ in the primary column, and $\widehat{\theta}_2$ towards zero or opposite ($|\widehat{\theta}_2| < |\widehat{\theta}_1|$, or opposite signs). Under the grade of the original specification, C is decisive when also $P_{\text{other}} < 0.001$ and $|z_c| \geq 3.0$, and suggestive otherwise. OA is decisive when the non-overlap fit that excludes zero has $|z| \geq 2.576$ and the overlap-aware profile $P \geq 0.10$, and suggestive otherwise. The class is compression, overlap amplification, both or neither. A pair classed neither has one label: the SNP axis does not reproduce the scalar signal (no row-(i) column excludes zero); no exposure signal is detected in the other six directions ($P_{\text{other}} \geq 0.05$); the outcome follows ($|z_c| < 1.96$); or other. When the primary-column cell of row (i) or (ii) is not a single bounded interval, $z_c$ and the C criterion are not evaluable, and the pair is counted as such. Boundary flags mark values near the cut-offs, for $|z_c|$ the band $[1.71, 2.21]$ (all flags, Supplementary Table 10(c)).

*Scope.* Curvature standard errors understate the sampling standard deviation when blocks are weak and share a genetic background, so the cuts at 1.96 and 3.0 are nominal. The classification, $z_c$ within the selected pairs and the concordance index are descriptive.

#### 6.6 Results

*Groups.* Four groups were analysed with the same code and rules: G1+O, the 123 pairs and the pair with opposite-sign estimates, O; the reference draws R1 and R2 (Section 6.5); and G2, the 47 scalar-only pairs whose F1 interval excludes zero and F2 interval includes it, added after the classification of the other three groups had been seen. $\pi_D^{-1/2}$, the ratio of the second- to the first-order standard error and the width of the F2 interval relative to its standard error were added after the other quantities, which had been defined in writing before they were computed. Every overlap-aware profile set was bounded, no pair had more than 20% of its loci excluded, and the added flag marked no pair. Group summaries, shares and concordance, and per-pair values are in Supplementary Tables 10(a), 10(b) and 10(c).

| | G1+O | R1 | R2 | G2 |
|---|---|---|---|---|
| Pairs (exposures) | 124 (48) | 100 (55) | 100 (53) | 47 (26) |
| Row-(i) first-order estimate over S: median; pairs within ±15% | 0.98; 122 | 1.00; 99 | 1.04; 64 | 0.99; 46 |
| $P_{\text{other}} < 0.001$ | 124 | 100 | 100 | 47 |
| Per-locus $W_2$ with $P < 0.05$: observed / expected | 2,807 / 186 | 2,503 / 165 | 2,202 / 147 | 1,250 / 81 |
| $\pi_D$, median (IQR) | 0.19 (0.18–0.23) | 0.19 (0.16–0.21) | 0.18 (0.16–0.21) | 0.20 (0.16–0.24) |
| Concordance index $r_1$, SNP axis: median ($n$) | 0.88 (85) | 0.91 (89) | 0.40 (57) | 0.73 (39) |
| Concordance index $r_2$, other six: median ($n$) | −0.07 (68) | 0.94 (87) | 0.57 (68) | 0.16 (37) |
| C under $z_c$ | 31 (25%) | 2 (2%) | 2 (2%) | 2 (4%) |
| C under $z_c^{\text{pre}}$ | 96 | 8 | 8 | 19 |
| OA (decisive) | 6 (0) | 16 (5) | 14 (0) | 11 (2) |
| Neither: outcome follows / SNP axis does not reproduce S | 85 / 2 | 80 / 2 | 9 / 75 | 34 / 0 |
| $\lvert z_c\rvert$ in $[1.71, 2.21]$ | 36 | 5 | 6 | 5 |

*The SNP axis and the exposure test.* Except in R2, whose S intervals include zero, the row-(i) first-order estimate reproduced the S estimate to within ±15% in nearly every pair. $P_{\text{other}}$ was below 0.001 in all 371 pairs, and per-locus $W_2$ exceeded its 5% point about 15 times as often as expected in every group, so $P_{\text{other}}$ does not separate the groups: by limit (3) above, a large $W_2$ means only that $b_X$ does not lie in the marker direction.

*Information and interval widths.* $\pi_D$ was defined in all 371 pairs, and $\pi_D^{-1/2}$, the RMS ratio of Section 6.4, had a median of 2.28 in G1+O (2.24–2.34 across the groups). The median of $w^{1/2}$ was 0.41–0.42 in every group and matched the landscape ratio of the F1 to the S standard error: on the same loci, the first-order founder interval is narrower than S because it uses all seven directions (Section 6.2). The F2 interval was wider than this (median width ratio to S, 0.53 in G1+O), because the overlap-aware second-order standard error, which includes exposure noise and the overlap term, was 1.18–1.27 times the first-order one.

*Concordance.* In G1+O, $r_1$ had an IQR of 0.60–1.39 and $r_2$ of $-0.22$ to 0.01; $r_2$ was negative in 49 of the 68 pairs with a value, and in the 55 pairs with both indices, $r_1 - r_2$ had a median of 0.85, against 0.00 in the 79 such pairs of R1. $r_1$ was negative in no pair of G1+O, R1 or G2 and in 7 pairs of R2.

*Compression and overlap amplification.* The direction part of C held in all 124 G1+O pairs, consistent with their selection (Section 6.5), and in all 47 G2 pairs, so in these groups C was decided by $|z_c|$. Of the 31 G1+O pairs in C, 30 had $\widehat{\theta}_1$ and $\widehat{\theta}_2$ of opposite signs; they came from 17 exposures, so they are not 31 independent observations. No pair reached $|z_c| \geq 3$, so under the grade of the original specification every C pair is suggestive, and no pair met both C and OA. In G2, the SNP-axis signal of 11 pairs was significant only when sample overlap was ignored (OA); 2 pairs, with $\widehat{\theta}_1$ and $\widehat{\theta}_2$ of the same sign and from 2 exposures, met C; and the other 34 were classed neither, all because the outcome follows.

*Two examples.* Two pairs were chosen by a rule fixed before the example values were computed: the O pair, and the G1 pair in C with the largest $|z_c|$. In the O pair, displacement at maximum load → osteoclast surface per bone surface (Oc.S/BS), the S estimate was $-0.30$; the first-order estimates were $\widehat{\theta}_1 = -0.31$ on the SNP axis and $\widehat{\theta}_2 = 0.13$ in the other six directions, so that F1 was 0.10; $z_c = 2.36$ ($z_c^{\text{pre}} = 2.98$). In the G1 pair, post-yield deflection → bone formation rate per total volume (BFR/TV), the S estimate was 0.51, $\widehat{\theta}_1 = 0.52$ and $\widehat{\theta}_2 = -0.07$, so that F1 was 0.06; $z_c = -2.85$ ($z_c^{\text{pre}} = -3.24$; Fig. 4d,e of the main text). Both pairs are in C. Supplementary Fig. 1 shows both pairs locus by locus (a, the O pair; b, the G1 pair) and, at two loci of the O pair, the founder effects split into the part the marker SNP keeps and the part it drops (c, d).

*Limits.* First, the group contrasts partly reflect selection: entry to G1 pulls $\widehat{\theta}_2$ towards zero or past it (Section 6.5), which by itself implies concordance on the SNP axis and discordance in the other six directions, and entry to R1 favours $\widehat{\theta}_2$ following $\widehat{\theta}_1$. The only pre-specified quantity free of the selection, $P_{\text{other}}$, is saturated, and asymmetric leakage of other loci's effects between exposure and outcome could produce apparent compression that is not separated here. Second, $r_k$ is missing for 39 ($r_1$) and 56 ($r_2$) of the 124 G1+O pairs, in which $\Gamma_{YY,k}$ was not positive because the outcome signal is close to its noise level, so the medians use a subset selected towards larger outcome signals; and the two indices, with $M'$ and $6M'$ coordinates, have different noise levels. Third, the C count depends on the denominator of $z_c$ (31 G1+O pairs under $z_c$, 96 under $z_c^{\text{pre}}$), and 36 G1+O pairs, including 14 of the 31 C pairs, lie in the boundary band of $|z_c|$; because the denominator of $z_c$ was itself below the robust bootstrap spread in the pilot, a wider denominator would lower the count further.

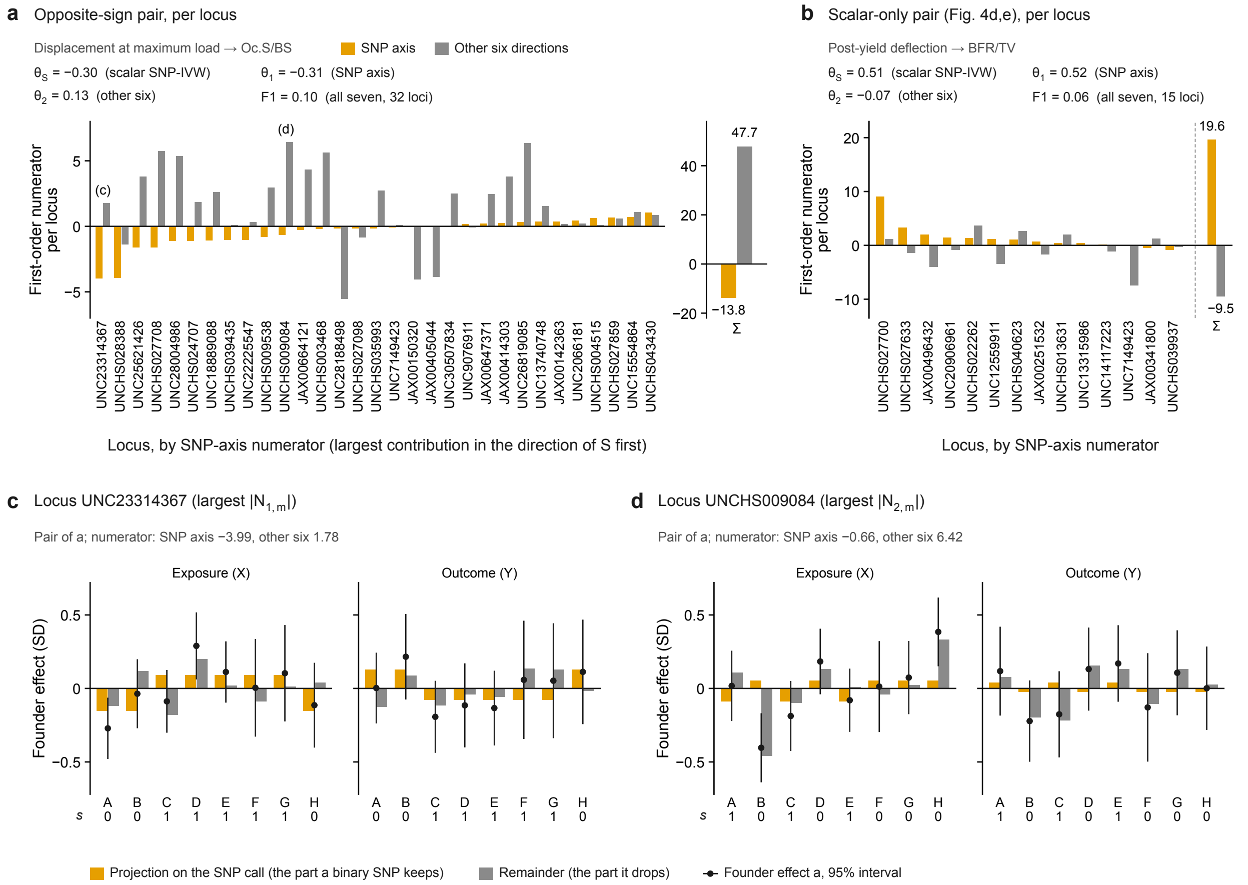


Supplementary Fig. 1: **Per-locus contributions for the two example pairs, and founder effects for the pair with opposite-sign estimates.** **a**, Displacement at maximum load → osteoclast surface per bone surface (Oc.S/BS), the one pair of the QC cohort whose S and F2 intervals both exclude zero, with opposite signs (Fig. 4b). For each of its 31 complete loci (of 32), the first-order numerator on the SNP axis ($N_{1,m}$, orange) and in the other six founder directions ($N_{2,m}$, grey), with loci ordered by $N_{1,m}$; right, their sums over loci ($\Sigma$, on its own scale). With the matching first-order denominators $D_{1,m}$ and $D_{2,m}$, $\widehat{\theta}_1 = \sum_m N_{1,m}/\sum_m D_{1,m}$ and $\widehat{\theta}_2 = \sum_m N_{2,m}/\sum_m D_{2,m}$ are the first-order estimates of rows (i) and (ii) (Section 6.2). Printed above the panel: $\widehat{\theta}_{\mathrm{S}}$, the scalar SNP-IVW estimate; $\widehat{\theta}_1$; $\widehat{\theta}_2$; and F1, the first-order estimate on all seven directions and all 32 loci. (c) and (d) mark the loci shown in **c** and **d**. **b**, The same display for post-yield deflection → bone formation rate per total volume (BFR/TV), the scalar-only pair of Fig. 4d,e, at its 14 complete loci (of 15); the sums follow the dashed line, on the same scale. F1 uses all 15 loci. **c**, **d**, Two loci of the pair in **a**: UNC23314367, with the largest $|N_{1,m}|$, and UNCHS009084, with the largest $|N_{2,m}|$. For the exposure (X) and the outcome (Y), the centred founder effects $a_f$ of founders A–H in trait standard deviations (dots; lines, $a_f \pm 1.96$ standard errors), split into the projection on the SNP call, $\alpha(s_f - \bar{s})$ (orange), the part that a binary SNP genotype keeps, and the remainder, $a_f - \alpha(s_f - \bar{s})$ (grey), the part that it drops. These are the founder-effect forms of $\alpha c$ and $r$ in $b = \alpha c + r$, with $\alpha = c^{\mathsf{T}}\Sigma_Y^{-1}b/c^{\mathsf{T}}\Sigma_Y^{-1}c$ for $b = b_X$ and for $b = b_Y$, both in the outcome metric $\Sigma_Y^{-1}$ (Section 6.1). Under each founder, $s_f$: 1 for founders with the second of the marker's two alleles in the R/qtl2 genotype data, 0 for those with the first (coding the other allele as 1 leaves the bars unchanged; Section 6.1). Founders as in Supplementary Note 2.
